# Deciphering the temporal heterogeneity of cancer-associated fibroblast subpopulations in breast cancer

**DOI:** 10.1101/2020.05.05.070607

**Authors:** Freja Albjerg Venning, Kamilla Westarp Zornhagen, Lena Wullkopf, Morteza Chalabi Hajkarim, Kyoung Jae Won, Chris Denis Madsen, Janine Tera Erler

## Abstract

Cancer-associated fibroblasts (CAFs) comprise a heterogeneous population of stromal cells within the tumour microenvironment. CAFs exhibit both tumour-promoting and tumour-suppressing functions, making them exciting targets for improving cancer treatments. A careful identification and characterisation of the CAF heterogeneity is thus necessary before implementing CAF-targeted strategies in cancer. With that in mind, we developed a flow cytometry strategy based on exclusion of non-CAF cells and successfully employed it to explore the temporal heterogeneity of CAFs in two models of triple-negative breast cancer (4T1 and 4T07). Analysing 128 murine tumours we identified 5-6 main CAF subpopulations and numerous minor ones based on the analysis of alpha smooth muscle actin, fibroblast activation protein alpha, platelet derived growth factor receptor alpha and beta, CD26/DPP4 and podoplanin. All markers showed temporal changes, and CD26+ CAFs emerged as a large novel subpopulation. These results form the foundation needed for the future elucidation of tumour-promoting CAF subpopulations.

## Introduction

Cancer-associated fibroblasts (CAFs) partake in a plethora of pro-tumorigenic functions, such as supporting cancer stem cells, providing protection against chemotherapy, and creating an immunosuppressive tumour microenvironment (TME) (recently reviewed in (Gieniec et al., 2019)). This has sparked great interest in targeting CAFs to treat various types of solid cancers (Chen and Song, 2019), as CAFs are generally thought to be genetically stable, in comparison to cancer cells, and thus appear as promising drug targets that may not develop resistance due to mutations. On the other hand, resident fibroblasts within the tissue can present a barrier to oncogenic transformation of epithelial cells (Dotto et al., 1988) and limit the proliferation of neoplastic cells (Kaukonen et al., 2016), exemplifying desirable fibroblast functions. Emerging evidence also supports anti-tumorigenic potentials of CAFs (Rhim et al., 2014; Özdemir et al., 2014), adding a layer of complexity to CAF-targeting strategies: who are the proverbial bad guys, and how do they distinguish themselves from the tumour-suppressive CAFs? While studies reporting on tumour-suppressing functions of CAFs are clearly the minority, they nevertheless highlight the need for gaining more knowledge regarding CAF subpopulations with potentially opposite functions, if targeting of CAFs is to be a successful add-on to cancer treatments.

Intra-tumoural heterogeneity of cancer cells in breast cancer is well recognised, and emerging evidence points toward breast cancer CAFs being equally heterogeneous (Bartoschek et al., 2018; Busch et al., 2017; Calvo et al., 2013; Costa et al., 2018; Cremasco et al., 2018; Raz et al., 2018; Su et al., 2018). While cancer cell heterogeneity is believed to arise through clonal evolution, the origins of CAFs, and how this influences the CAF heterogeneity, are still being intensely studied. Evidence points toward multiple origins rather than one, with CAF-progenitor cells heralding from as diverse sources as adipose stem cells, endothelial cells, resident fibroblasts, bone marrow (BM)-derived mesenchymal stromal cell (MSCs), and even from malignant cancer cells themselves (reviewed in (Gieniec et al., 2019) and (Chen and Song, 2019)). A study using the MMTV-PyMT mouse model of breast cancer provides evidence that the origin of CAFs is reflected into functionally different CAF subpopulations: CAFs derived from BM-derived MSCs promote angiogenesis, while CAFs deriving from resident fibroblasts help recruit tumour associated macrophages (Raz et al., 2018). Another study used single cell RNA sequencing (scRNA-seq) to define three CAF subpopulations within late-stage MMTV-PyMT breast tumours (Bartoschek et al., 2018). The authors suggested the use of certain single markers to discriminate between the three populations, and while they were able to show some intriguing spatial locations of these markers within mouse breast tumours, they did not look into co-expression of these or other CAF markers on the protein level. Heterogeneity of breast cancer CAFs is not just a phenomenon in experimental cancer models, as human breast tumours also have been demonstrated to encompass at least four CAF subpopulations, linking one of these subpopulations to the development of an immunosuppressive TME (Costa et al., 2018).

Currently no study has looked at the temporal composition of CAF subpopulations during the development of the breast tumour, and how such a co-evolution of CAF subpopulations might depend on the metastatic potential of the tumour cells. To investigate these questions, we performed orthotopic implantation of syngeneic 4T1 or 4T07 cancer cells into the mammary fat pad of female mice. The two cell lines are both models of triple-negative breast cancer (TNBC) and originate from the same spontaneous breast tumour in a BALB/c mouse, but differ in their metastatic potential. 4T1 cells form primary tumours in syngeneic mice that readily spread and form macro-metastases in the lungs, liver, bone, and brain. 4T07 cells form primary tumours, but these tumours only manage to spread to the lymph nodes and to form micro-metastases in the lungs of syngeneic mice (Aslakson and Miller, 1992). When isolating CAFs from different time points during tumour development, these two models offer the opportunity to study cancer cell intrinsic abilities to educate and recruit different CAF subpopulations at different time points, in the setting of a fully functioning immune system.

Seeking to provide the CAF research community with an unbiased starting point for investigating the CAF heterogeneity in the context of any type of solid tumours in mouse cancer models, we have developed a multicolour flow cytometry (FCM) workflow, based on the principle of negative selection. By excluding essentially all other cell types in the TME, a CAF-enriched population is left behind ready for further *ex vivo* analysis. In this way, we purify living cells and avoid limiting the CAF population to predefined markers that are not all encompassing. Our set-up also avoids the ‘plastic-education’ typically observed when CAFs are isolated by letting the CAFs migrate out of a small piece of tumour to subsequently adhere to the culture dishes for days. As a proof of concept, we here investigate for the first time the co-expression of six CAF markers on primary breast cancer CAFs over time, and show how these markers define numerous specific CAF subpopulations that change in abundance as the tumours grow.

## Results

### Unbiased detection of CAFs through negative selection strategy

We carried out three independent biological repeats of the orthotopic breast tumour model shown in Fig 1A, implanting either 5×10^5^ 4T1 or 4T07 cells into the mammary fat pad of the αSMA-RFP BALB/c mice, analysing a total of 128 tumours and 12 healthy mammary fad pads. We chose to employ a negative selection strategy in order not to limit our dataset to one specific CAF marker, giving us an unrestricted starting point for investigating CAFs in 4T1 and 4T07 tumours. We designed an antibody cocktail of lineage markers to exclude non-fibroblast cells from the single cell suspension of tumours analysed by FCM (Fig 1A) (see Materials and Methods for a list of antibodies used). To exclude cancer cells we initially used eGFP labelled cancer cells, however pilot experiments suggested that eGFP expression was either lost or greatly diminished from the tumours around 14 days after implantation (data not shown), prompting us to choose another way of excluding the cancer cells. The expression of EpCAM, CD24, and CD49f have been used to detect different subtypes of breast cancer cells (Keller et al., 2010), thus we tested the expression of these markers in the 4T1 and 4T07 cell lines (Fig S1). The more metastatic 4T1 cell line expressed all three markers, while the less aggressive 4T07 only expressed CD24 and CD49f. As it was previously reported that multiple different populations can be found in cultured breast cancer cell lines (Keller et al., 2010), we decided to include all three markers in our lineage cocktail to exclude as many cancer cells as possible. The population of sorted live cells that did not express any of the lineage markers was termed lineage negative (Lin-) (Fig 1B).

**Figure 1.**
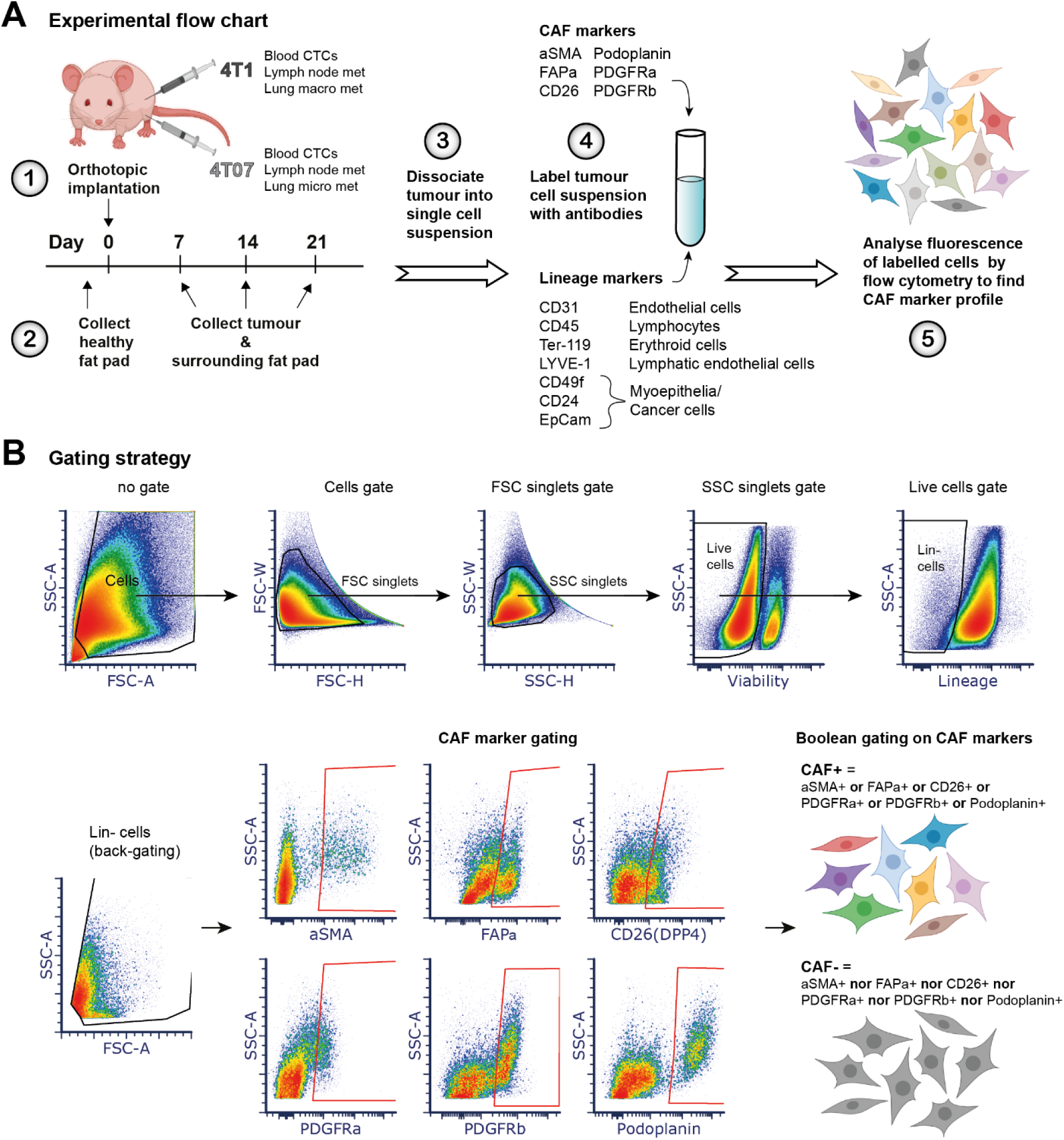
Experimental set-up and gating strategy for flow cytometry analysis. **A)** Schematics of experimental setup. 4T1 or 4T07 breast cancer cells were injected into the mammary fat pad of aSMA-RFP mice. 4T1 and 4T07 are sister cell lines derived from the same tumour, but have different metastatic profiles as denoted next to them in the schematic. CTCs= circulating tumour cells, met= metastases. **B)** Flow cytometry gating strategy to obtain live cells devoid of lineage markers (Lin-), and subsequent gating on each CAF marker used for Boolean gating to define the CAF- and CAF+ populations. The gates for all markers were set using the fluorescent minus one (FMO) approach. The density plots show one representative 4T1 D7 tumour (D7: collected at day 7). FSC = forward scatter, SSC = side scatter.

### Fibroblast population size depend on time and tumour type

Most reports point towards CAFs having a tumour supportive role, regardless of the CAF marker chosen to represent the CAF population. Thus, the initial question we sought to answer was what percentages of live cells in the TME are CAFs? Due to our negative selection strategy, the Lin-population is highly enriched for fibroblasts and CAFs, and we investigated whether this population differed in size during tumour progression and between the two tumour types. We first evaluated the percentage of purified Lin-cells in healthy mammary fat pad and observed an average of ∼10% Lin-cells in 8-16 weeks old mice without tumours. Not surprisingly, this percentage decreased as the tumours developed (day 7 (D7), day 14 (D14), day 21 (D21)), simply because the Lin-cells were being outnumbered by the expanding and proliferative tumour cells in the mammary fat pad (Fig 2A). Despite this, the percentage of Lin-cells still seemed to increase during tumour growth, as depicted by the linear fit of the data from both 4T1 (p=0.0653) and 4T07 (p=0.0628) tumours. Another interesting observation was that 4T07 tumours had a significantly higher percentage of Lin-cells as compared to 4T1 tumours, irrespective of tumour age (Fig 2A and Suppl. Fig 3B). These observations demonstrate that the numbers of Lin-cells are rising as the tumour grow, and that the two tumour types have different levels of Lin-cells.

**Figure 2.**
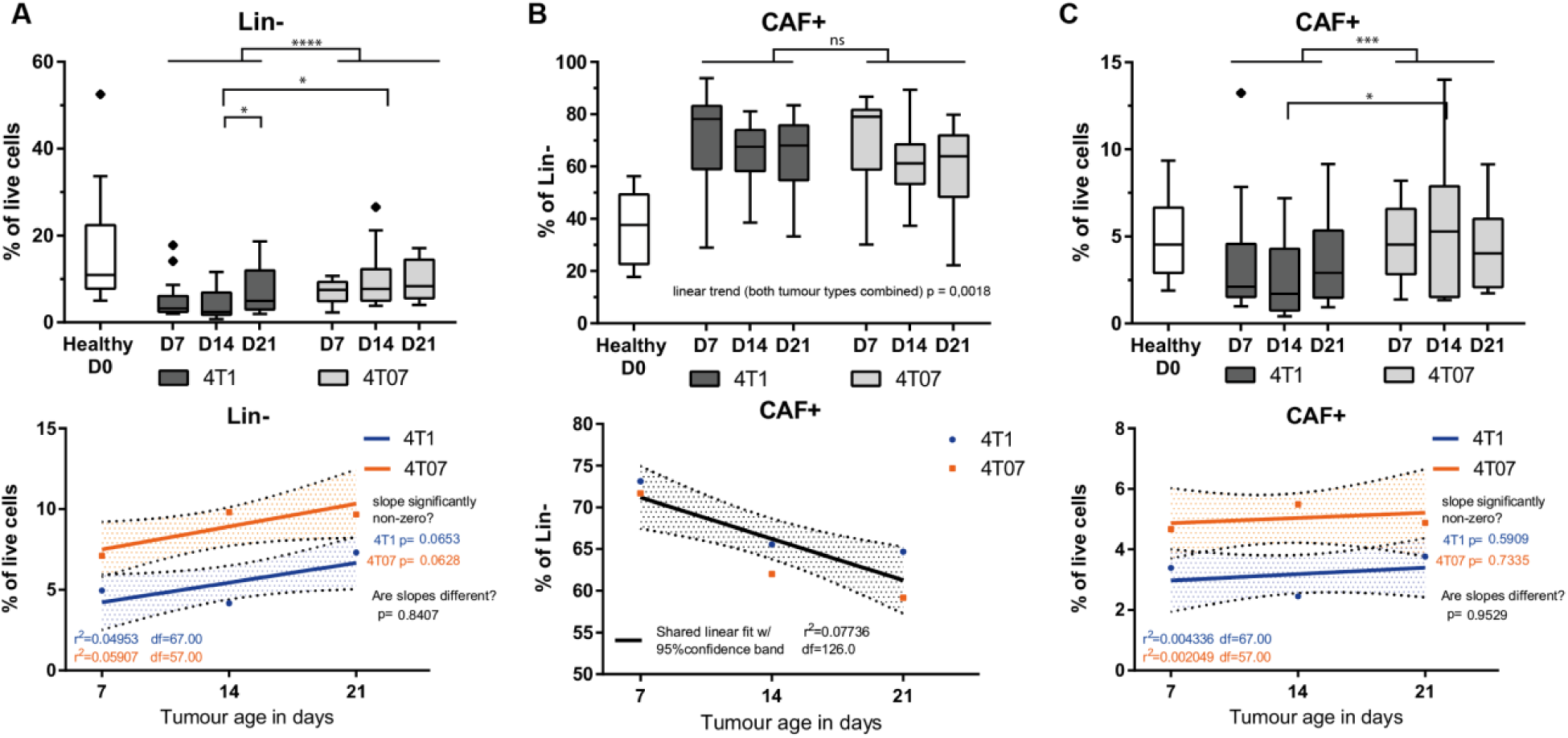
Analysis of 128 tumours reveal temporal dynamics and an effect of tumour type. **A-C)** Percentage of cells in healthy mammary fat pad (D0) and 4T1 and 4T07 tumours across the different time points. Tukey box plots of two (healthy samples) or three independent repeats (tumour samples) combined. The boundaries of the Tukey style box go from the 25^th^ to the 75^th^ percentile, and the median is depicted by a line. The healthy samples provide a D0 reference point, but all the statistics are only concerning the tumour samples. Fitting a straight line to the tumour data, testing if one line fits both 4T1 and 4T07 tumours to indicate no difference, or if slope and/or y-intercept differ if two lines better capture the dataset. Fitted lines are shown with hashed out 95% confidence bands (CB95), with dots indicating the observed mean at the respective time point. **A)** 2-way ANOVA analysis finds that both tumour type and day affects the size of the Lin-population, in relation to live cells. Multiple comparisons post-test finds 4T1 D21 to be significantly larger than 4T1 D14, and multiple unpaired t-test show a significant difference between tumour types at D14 (False discovery rate (FDR) approach, Q=1%). Linear fitting of the dataset shows two different fits are needed, both with similar slopes and their CB95 containing 0, but with different y-intercepts, i.e. 4T1 tumours have fewer Lin-cells than 4T07 tumours in general with a possible trend towards an increase in both tumours. **B)** Mean percent CAF+ population across the different time points, showing that even the combination of all 6 CAF markers only captures about 60-80% of the cells in the Lin-population. Linear fitting shows that one line with a negative slope describes the dataset better than two different lines, i.e. the relative CAF+ population within the Lin-population decreases with age in the same manner in both tumour types. 1-way ANOVA analysis with linear trend post-test reveals that there is a significant linear decrease in the CAF+ population as tumours age. **C)** CAF+ population size in relation to live cells in the tumour, here only tumour type effects the population size (2-way ANOVA), and multiple unpaired t-tests with FDR correction reveals that the tumour types differ at D14. Linear fitting shows that two separate lines are needed to describe the dataset, both with similar slopes and their CB95 containing 0. However, the fits do have different y-intercepts, with 4T07 tumours having more CAF+ cells than 4T1 tumours. To determine statistical significance 2-way ANOVA with Tukey’s multiple comparisons post-test was used for intra-tumour comparisons, and multiple, two-tailed, t-tests with FDR correction (Q=1%) was used for inter-tumour comparisons. *= p<0.05, **=p<0.01, ***=p<0.001, **** p<0.0001. Healthy n=12, 4T1 D7 n=21, 4T1 D14 n=24, 4T1 D21 n=24, 4T07 D7 n=24, 4T07 n=21, 4T07 D21 n=14.

### Six CAF markers combined capture the majority of Lin-cells

A multitude of markers have been proposed as CAF markers, including but not limited to: alpha smooth muscle actin (αSMA), fibroblast activation protein alpha (FAPα), fibroblast specific protein-1 (FSP1/S100A4), vimentin, Podoplanin (PDPN), platelet derived growth factor receptor alpha and beta (PDGFRα and PDGFRβ), CD90 (Thy.1), neuron glial antigen 2 (NG2), Caveolin-1 (Kalluri, 2016), and recently integrin a11 (Primac et al., 2019; Schnittert et al., 2019). As these markers are also expressed on other cell types in the TME, it remains unclear to what extend these markers truly represent the total level of CAFs within the TME. To allow for future FACS-based isolation and subsequent live-cell studies we focused our analysis on cell-surface markers. We therefore took advantage of the αSMA-RFP mouse that expresses red fluorescent protein (RFP) under the alpha smooth muscle actin (αSMA) promotor (LeBleu et al., 2013), considered to be the *bona fide* marker of activated myo-fibroblasts and myofibroblastic CAFs. We then designed an antibody panel consisting of 5 cell surface CAF markers: FAPα, PDGFRβ, PDGFRβ, PDPN and dipeptidyl petidease-4 (DPP4/CD26). FSP1/S100A4 is a commonly used CAF marker, however it is an intracellular calcium binding protein and not believed to be routinely present on the cell surface, and thus we did not include it in our panel. We included the atypical CAF marker CD26, also known as DPP4, as it is a lineage marker of fibrogenic dermal fibroblasts (Rinkevich et al., 2015) and a close homologue of the well-known CAF marker FAPα (Scanlan et al., 1994).

To simplify the analysis we chose to make the presence of each CAF marker a binary question, assigning each cell to either the positive or negative population based on fluorescent minus one (FMO) gating. We did this for each of the 6 markers, and using Boolean criteria we could define two Lin-populations: a) Lin-cells that do not express any of the 6 selection markers (we termed this population CAF-) and b) Lin-cells that express at least one of the 6 markers (we termed this population CAF+) (Fig 1B). As anticipated, our experimental setup with 6 CAF markers did not lead to full coverage of the Lin-population, although still capturing 60-80% of the Lin-cells in tumours (Fig 2B). The analysis also demonstrated that the percentage of CAF+ cells in relation to live cells was relatively stable over time in tumours (Fig 2C, lower panel). Interestingly, the overall percentage of CAF+ cells in relation to Lin-cells was actually decreasing during cancer development regardless of tumour type (Fig 2B, linear trend p=0,0018), meaning that the CAF-population was increasing. The CAF-population, not expressing any of the 6 markers, may therefore represent other CAF subpopulations expressing markers not used in this panel. In summary, our workflow using 5 well-established markers and one new CAF marker (CD26) captured 60-80% of all Lin-cells in the primary tumours, and it is thus reasonable to assume that our workflow captured the majority of *bona fide* CAFs.

### CAF marker prevalence depends on tumour age and type

We then sought to explore how each individual CAF marker was temporally present in each tumour type. The first observation was a dramatic switch in all of the 6 CAF populations when comparing the healthy mammary fat pad to the breast cancer tissue. The healthy tissue had a strong percentage of ∼90% PDGFRα+ cells that drastically decreased as the tumour grew (Fig 3A). Oppositely, αSMA, PDGFRβ, FAPα and CD26+ CAF populations were all significantly appearing/expanding as soon as a tumour was established (Fig 3A). Importantly, these alterations were independent on tumour type as they were observed in both 4T1 and 4T07 tumours. The second observation was that the percentage of each individual CAF marker was not stable over time. For instance, the αSMA, PDPN and PDGFRβ+ populations showed a decrease over time in the CAF+ population, with linear fitting of the data indicating similar temporal dynamics of all three markers in both 4T1 and 4T07 tumours (Fig 3). In the case of the other three markers, FAPα, CD26 and PDGFRα, these showed divergent dynamics across time in the two tumour types. This was revealed by the 2-way ANOVA analysis demonstrating a significant interaction between the two independent variables (time and tumour type) as illustrated by the linear fitting showing the two fitted lines crossing each other (Fig 3). Interestingly, CD26 appeared to have completely opposite dynamics in the two tumour types. In 4T07 tumours, there was a significant decrease over time from D7 to D14/D21 (Fig 3A). In 4T1 tumours, comparisons between each time points found no significant differences, yet linear fitting suggested an increase over time (slope significantly non-zero), culminating in a significant difference between tumour types at D21 (4T1 > 4T07). Similar dynamics were seen when CAF markers are examined within the Lin-population (Fig S4).

**Figure 3.**
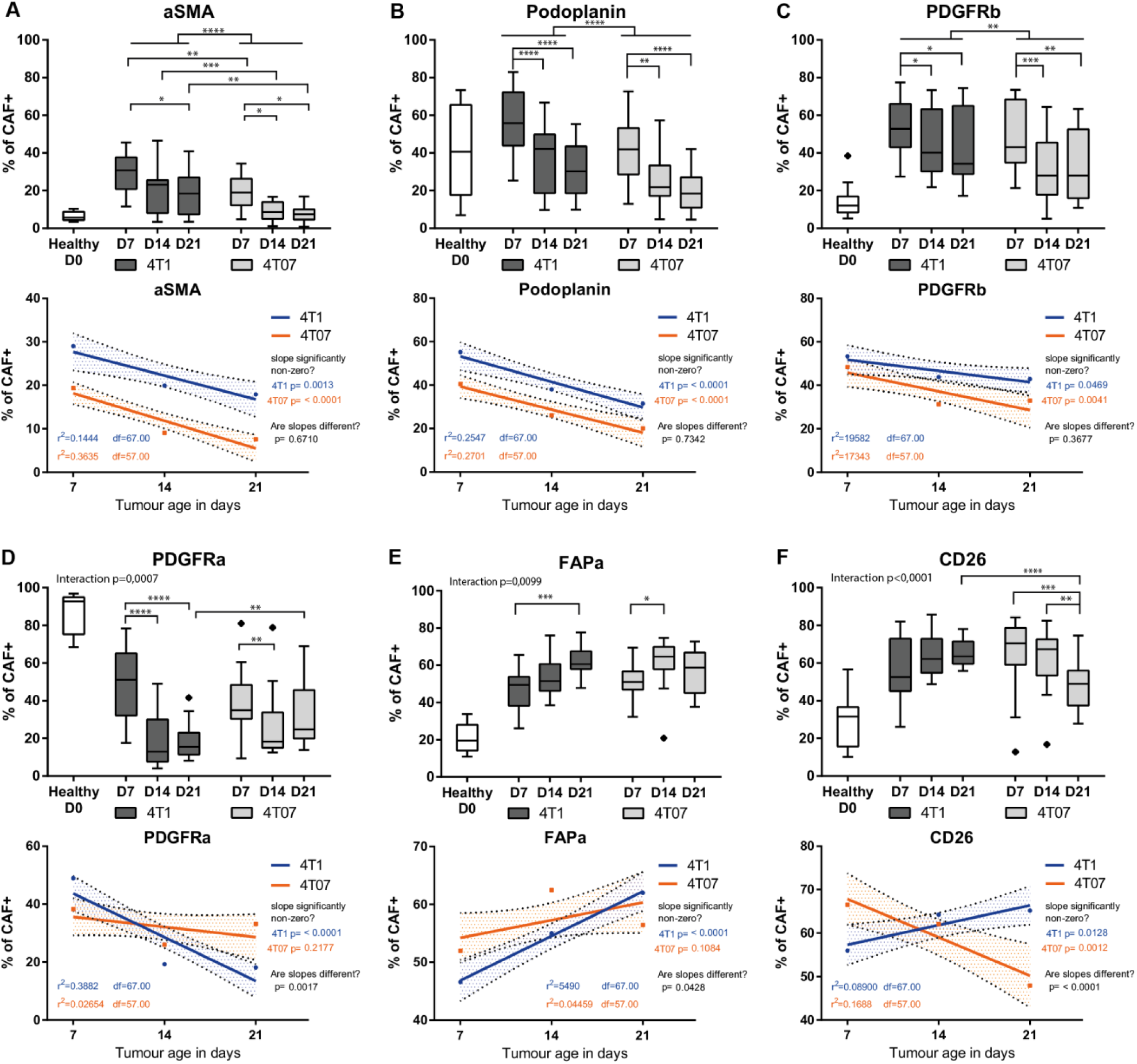
All 6 CAF markers show temporal dynamism and in some cases tumour type dependent dynamics. Analysis of the CAF+ population of 128 tumours**. A-F)** Percentage of cells in healthy mammary fat pad (D0) and 4T1 and 4T07 tumours across the different time points positive for the respective CAF marker. Tukey box plots of two (healthy samples) or three independent repeats (tumours samples) combined. The boundaries of the Tukey style box go from the 25^th^ to the 75^th^ percentile, and the median is depicted by a line. The healthy samples provide a D0 reference point, but all the statistics are only concerning the tumour samples. Fitting a straight line to the data, testing if one line fits both 4T1 and 4T07 tumours to indicate no difference, or if slope and/or y-intercept differ if two lines better capture the dataset. Fitted lines are shown with hashed out 95% confidence bands (CB95), with dots indicating the observed mean at the respective time point. 2-way ANOVA was run once for each marker separately to look for interaction and over-all effect of tumour type and/or day. To determine statistical significance between time points within each tumour type (intra-tumour time comparisons) a 2-way ANOVA with Tukey’s multiple comparisons post-test was run on all the CAF markers combined, once for 4T1 tumours and once for 4T07 tumours. To determine if markers differed between tumour types, multiple unpaired, two-tailed t-tests without assuming equal variance and with FDR correction (Q=1%) was run for each of the three time points (inter-tumour comparison). *= p<0.05, **=p<0.01, ***=p<0.001, **** p<0.0001. Healthy n=12, 4T1 D7 n=21, 4T1 D14 n=24, 4T1 D21 n=24, 4T07 D7 n=24, 4T07 n=21, 4T07 D21 n=14.

Taken together, data from the 128 tumours demonstrate that the development of a malignant breast tumour dramatically shifts the population of fibroblasts/CAFs in the mammary fat pad, and that once the tumour is established the dynamics of each individual CAF marker keeps changing over time. αSMA, PDPN, and PDGFRβ behave similarly and all decrease over time in both tumour types, while the newest CAF marker, CD26, stands out as the only marker with intriguing opposite dynamics over time when comparing the two different tumours: increasing in the aggressive 4T1 while decreasing in 4T07. The common denominator in these analyses is that the biggest changes seem to take place between D7 to D14.

### A few CAF subpopulations dominate the TME

With the presence of 6 CAF markers recorded for each cell in the CAF+ population, the number of theoretically possible CAF subpopulations is 63 when using Boolean criteria. To investigate which of these theoretical subpopulations actually exist in the breast tumours, we plotted the CAF+ population of each tumour as a heat map, with each row representing one of the theoretical CAF subpopulations and the colouring representing the percentage of such subset in the tumour. It is clear from the heat map (Fig 4A and B) that many of the 63 theoretical Boolean CAF subpopulations were actually present in both tumour types and that a few CAF subpopulations were more prevalent than others were. It is noteworthy that many of the subpopulations are the same in both tumour types and remain prominent across time (Fig 4A and B). The three independent repeats in some cases seemed to be distinctly different, suggesting a degree of systemic technical noise from one repeat to the next (batch effect), likely due to a combination of differences in antibody staining intensities, as well as instrumental performance from day to day. We assessed this and found only a very minor batch effect within the combined dataset (see Fig S5 and its legend).

**Figure 4.**
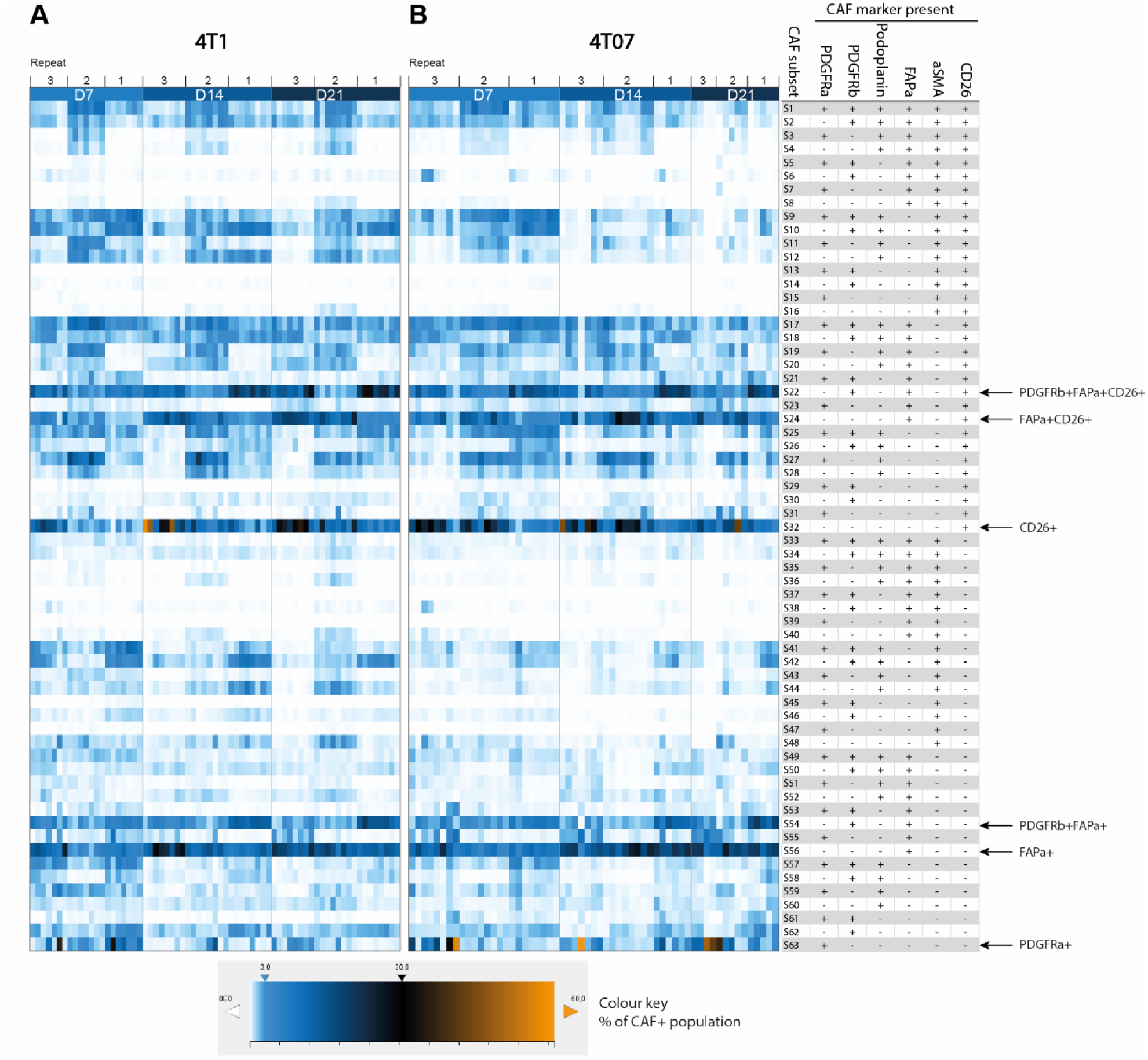
A few CAF subsets dominate the TME. **A**) and **B**) heat map visualisation of the prevalence of each theoretical Boolean CAF subset across time and tumour type within the CAF+ population. Each lane corresponds to one tumour and three independent repeats are combined as denoted above the heat maps (128 tumours in total). Each square shows the percent this CAF subset constitutes out of the CAF+ population, see colour key for reference. The heat maps clearly show that far from all the theoretical subsets are present and that a few CAF subsets dominate the TME across time and tumour type. 4T1 D7 n=21, 4T1 D14 n=24, 4T1 D21 n=24, 4T07 D7 n=24, 4T07 n=21, 4T07 D21 n=14.

### CAF subpopulations expressing FAPα and/or CD26 are the most prevalent

A commonality of the most abundant CAF subpopulations across both time and tumour type was the expression of either FAPα or CD26 as seen in Fig 5. The CAF+ population was more heterogeneous in 4T1 than in 4T07 tumours at D7. However, as the TME in both tumour types matured with time, it appears that the CAF+ population becomes slightly more homogeneous over time in both 4T1 and 4T07, with the top 5 CAF subpopulations increasing in abundance to finally cover more than 60% (Fig 5). Including the top 10 CAF subpopulations only added 10-20% more to each time point (Table S8), thus demonstrating how relative rare most of the theoretical subpopulations are.

**Figure 5.**
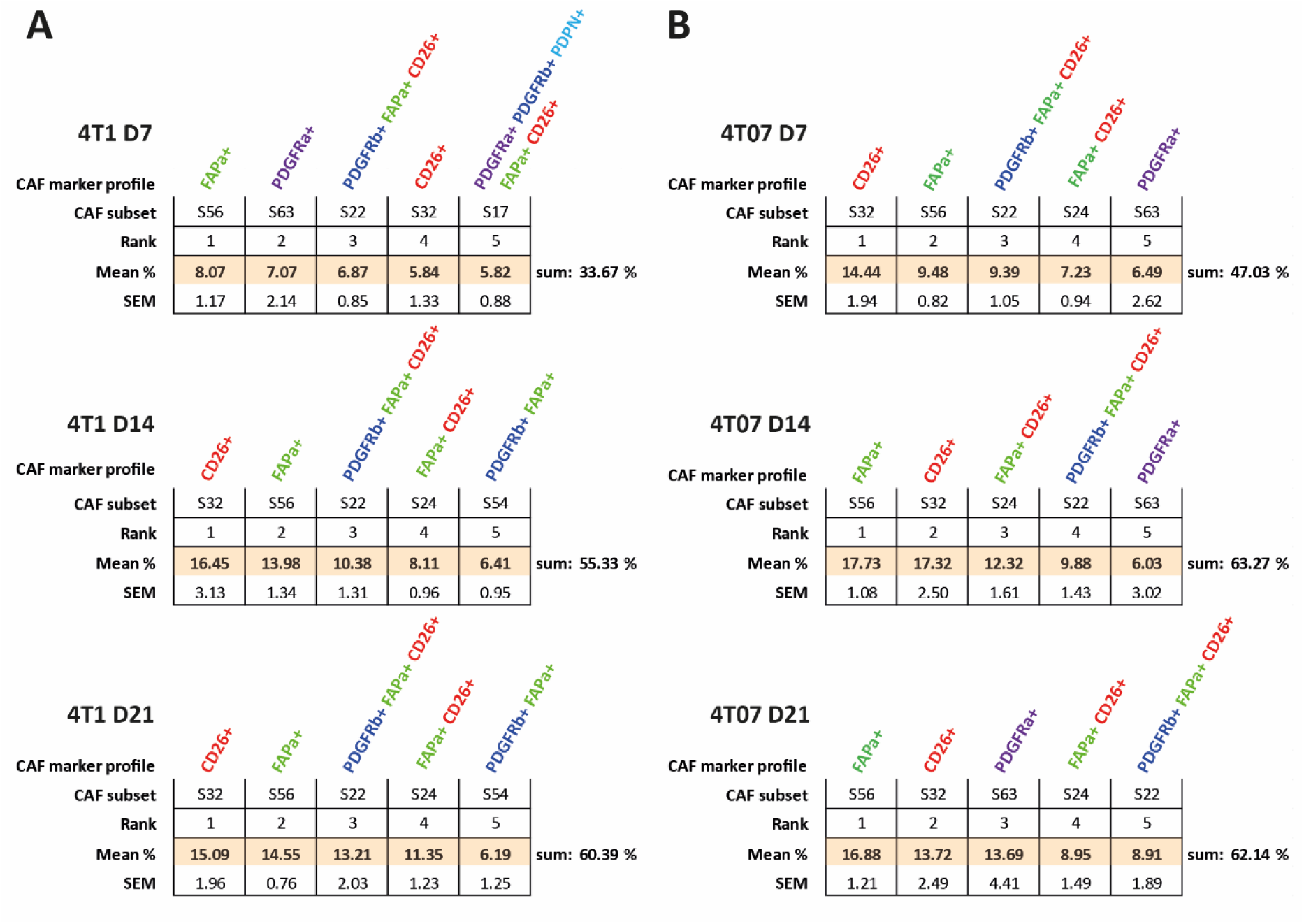
FAPα+ and CD26+ CAF subpopulations dominate the TME at all times. Top 5 most abundant Boolean CAF subpopulations (subsets) in **A)** 4T1 and **B)** 4T07 tumours at a given time point (within the CAF+ population). The CAF population becomes less heterogeneous over time, and FAPα (green) and CD26 (red) expressing subsets dominate the TME. Mean % and SEM from three independent repeats combined. 128 tumours in total. 4T1 D7 n=21, 4T1 D14 n=24, 4T1 D21 n=24, 4T07 D7 n=24, 4T07 n=21, 4T07 D21 n=14. See supplementary materials for complete abundance ranking of all 63 CAF subsets.

In addition to using Boolean criteria to define CAF subpopulations, we analysed the Lin-population using UMAP (Uniform Manifold Approximation and Projection) dimension reduction (https://arxiv.org/abs/1802.03426) (McInnes et al., 2018) (Fig 6). This type of analysis does not rely on manual gating of the CAF markers into binary +/- populations, instead the intensity values of each of the 6 markers are used to guide the reduction in dimensions. Consistently, we saw only a few large clusters (subpopulations) and many smaller ones in both tumour types (Fig 6), in agreement with the Boolean analysis in figure 4. This analysis also confirmed that the largest clusters expressed a high level of FAPa and/or CD26 (Fig 6A & B).

**Figure 6.**
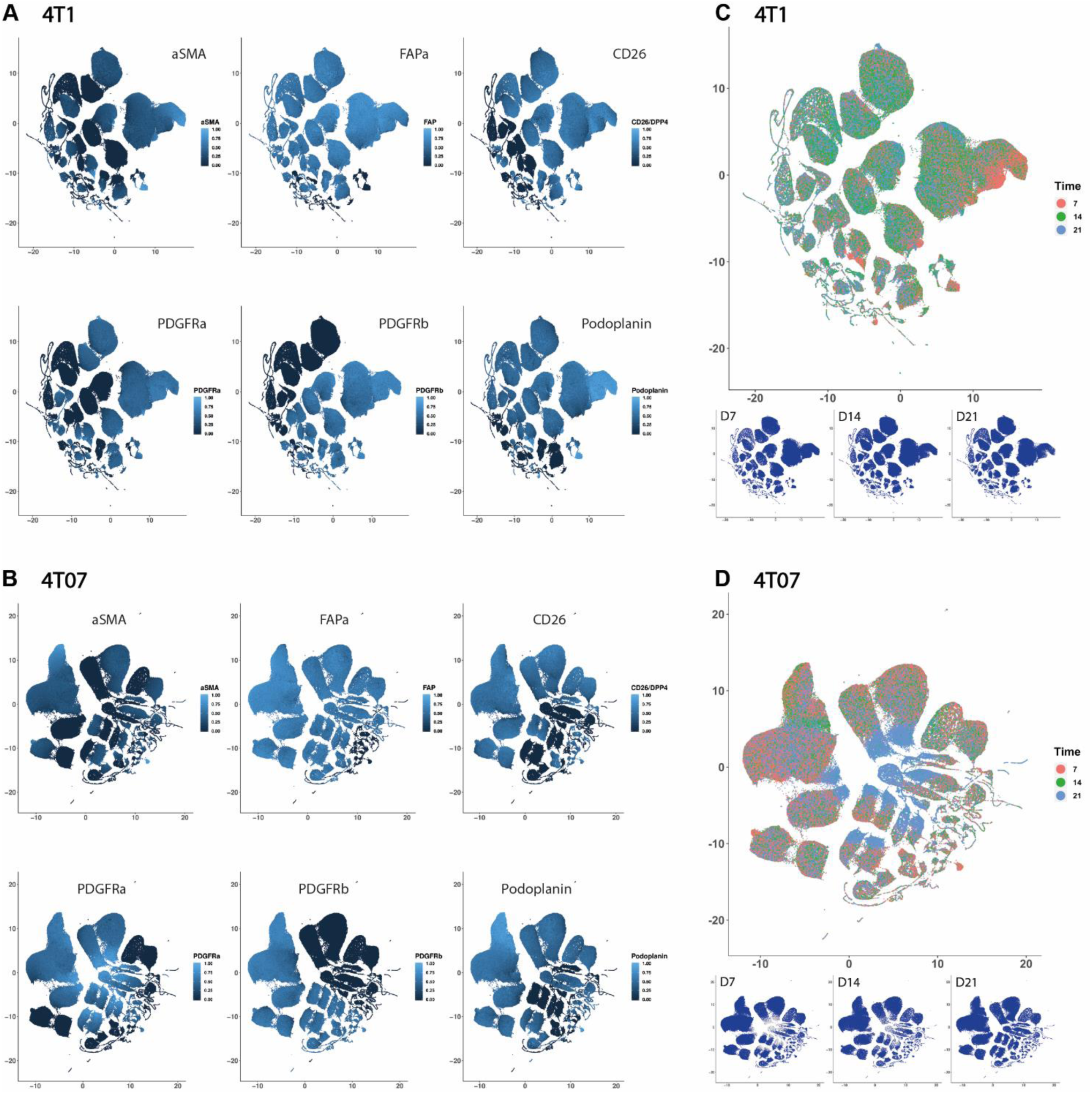
UMAP show a handful of large and many small CAF subpopulations. Dimension reduction by UMAP of all 6 CAF markers within the Lin- population. The intensity levels were normalised into a range from 0=no signal to 1=strongest signal. A and B) UMAP plots of each of the 6 CAF markers. C and D) UMAP plots of cells from day 7, 14, and 21 separately (small plots) and combined (large plot), showing how some clusters are more abundant at a given time point. Data shown here in this figure are from one biological repeat (rep 3), 4T1 n=23 and 4T07 n=19.

In summary, a minority of the 63 theoretical subpopulations are present at great numbers in both tumours, and many of these are shared, and express FAPα and/or CD26, hinting at the presence of a common core of CAFs between the two breast tumour types regardless of the metastatic potential of the tumour. 4T1 tumours appear to initially have a more heterogeneous CAF+ population than 4T07 tumours, yet over time, the CAF+ population in both tumours becomes less heterogeneous. Although the data clearly indicates that a handful of subpopulations is prevailing in the TME, it is important not to forget that many of the 63 theoretical combinations are actually present in the TME, albeit in low numbers, and that 20-40% of the Lin-population is not detected by our markers (Fig 2B).

### CAF+ population composition changes dramatically over time

The UMAP plots revealed that there was a change in abundance of some clusters over time (Fig 6C and D), thus we next wanted to see how CAF subpopulations co-existed in each tumour type at the different time points. To get a graphical overview of the complex composition of all 63 possible Boolean CAF subpopulations, we took advantage of the freeware SPICE (https://niaid.github.io/spice/), developed by the Roederer laboratory, to visualise our polychromatic FCM dataset (Roederer et al., 2011). The pie charts generated by SPICE are presented in Fig 7, with each of the possible CAF subpopulations being represented as a slice in the chart. From the first glance, it is evident that the composition of subpopulations making up the CAF+ population changes from D7 to D14 in both tumour types, and that the tumour types also appear to be different from one another. Using a permutation test to compare the pie charts it was possible to determine if the compositions of CAF subpopulation are statistically different from one another. As the dynamics of some of the single CAF markers (Fig 3) and the UMAP plots (Fig 6) suggested, a clear change in the relative abundances of the CAF subpopulations was taking place from D7 to D14 in both tumour types, while on the broad scale the total composition did not change significantly from D14 to D21 (Fig 7). When comparing the two tumour types the total CAF composition in 4T1 tumours was significantly different from that of 4T07 tumours at D7. The CAF composition then became more similar at D14, before diverging significantly again at D21. What the permutation test and the pie charts did not reveal was which CAF subpopulations specifically differ across time and/or tumour type, and we next moved on to look into this.

**Figure 7.**
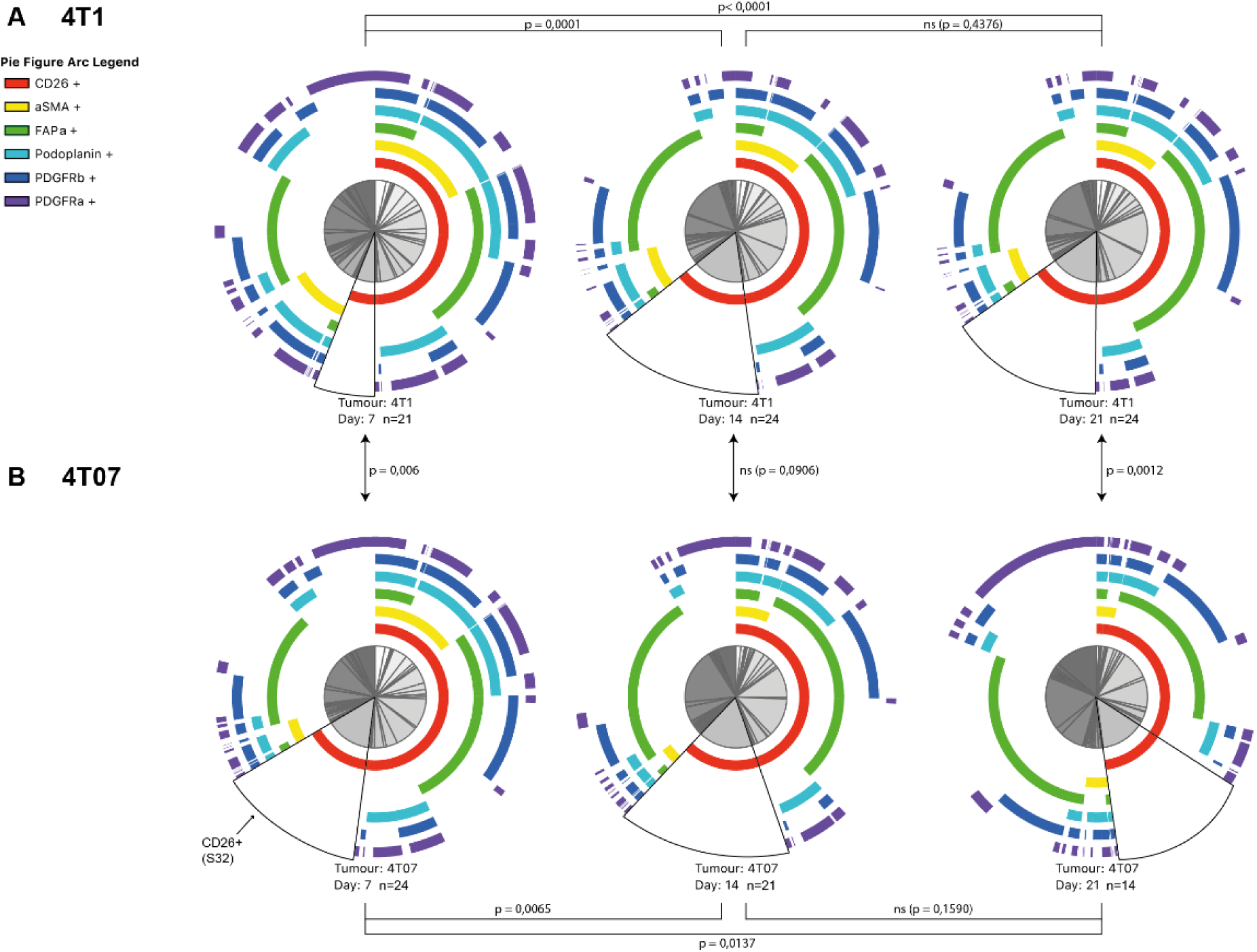
SPICE visualisation of Boolean CAF subsets. **A)** 4T1 tumours and **B)** 4T07 tumours, three independent repeats combined. The arcs around the pie charts corresponds to each of the 6 CAF markers, and the pie slices are read by looking at which arcs that are present outside of it. In this way, each pie slice corresponds to one theoretical Boolean subset of CAFs, subset S32 is marked on each pie with a black border as an example. It is evident that not all of the theoretical subsets are present in the tumours, and that the overall composition of CAF subsets change as tumours matures. Furthermore, the initial difference between tumour types seen at D7 is translated into a difference at the latest stage at D21. Statistical p-values are from permutation testing comparing the CAF subset composition within each pie to that of another pie. 128 tumours in total, 4T1 D7 n=21, 4T1 D14 n=24, 4T1 D21 n=24, 4T07 D7 n=24, 4T07 n=21, 4T07 D21 n=14.

To identify the CAF subpopulations that change significantly in size across time we performed a 2-way ANOVA on the 63 theoretical Boolean subpopulations, followed by Tukey’s multiple comparisons post-test. All the subpopulations found to change significantly with time are listed in Table 1. While both 4T1 and 4T07 tumours changed from D7 to D14 (Fig 7), 12 subpopulations were changing in 4T1 tumours compared to only 7 in 4T07 tumours (Table 1). However, many of the shifting subpopulations (S9, S17, S24, S32, and S56) were the same in both tumour types, indicating that both tumours mature their CAF+ population composition in a similar way. Comparing across tumour types at each time point it became clear that the tumours were significantly different regarding the overall composition of CAF subpopulations (see pie charts in Fig 7), but that this compositional difference was mostly made up by tiny subpopulations (Table 2). However, there were two noteworthy substantial differences between 4T1 and 4T07 tumours; at D7, CD26 single positive CAFs (S32) were almost 2.5 times as abundant in 4T07 compared to 4T1 tumours (14.4% vs 5.8%), and at D21 the PDGFRα single positive CAFs (S63) subpopulation was more than 9 times as abundant in 4T07 compared to 4T1 tumours (13.7% vs 1.5%) (Table 2).

**Table 1.**
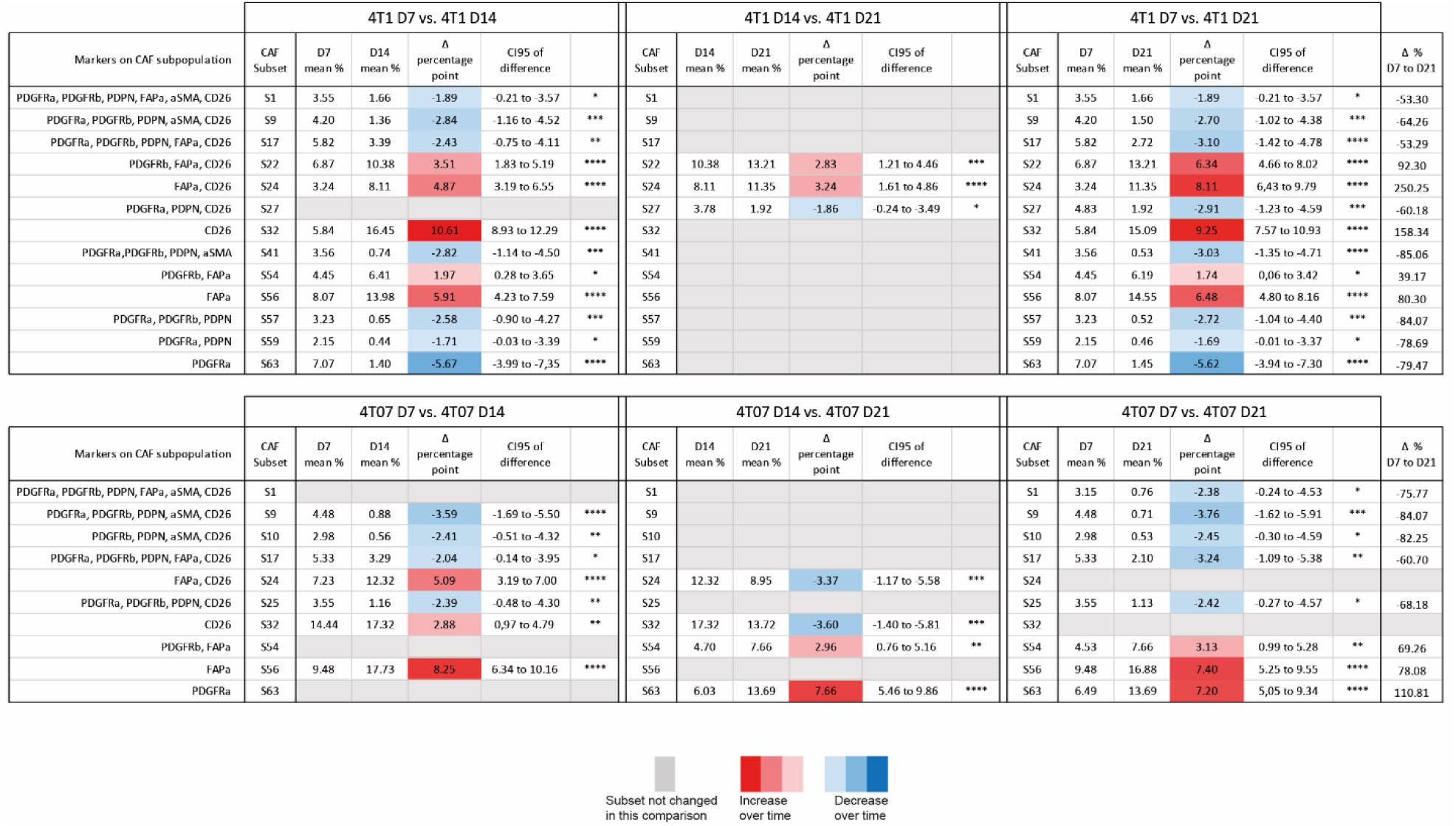
CAF subsets that change over time within each tumour type. All CAF subpopulations that were found to change significantly by 2-way ANOVA and Tukey’s multiple comparisons post-test are listed below. The mean difference in percentage point is colour-coded with red indicating an increase from the earlier to the later time point, and blue indicating a likewise decrease over time. Grey fields indicate that the subpopulations did not change over time in that particular comparison. Δ % = change in percent from D7 to D21. 128 tumours in total, 4T1 D7 n=21, 4T1 D14 n=24, 4T1 D21 n=24, 4T07 D7 n=24, 4T07 n=21, 4T07 D21 n=14.

**Table 2.** All CAF subsets that differ in size between 4T1 and 4T07 tumours at day 7, 14 or 21. CAF subpopulations that differ between the tumour types were identified by multiple t-tests without assuming equal variance and using the FDR approach (Q=1%) to correct for multiple comparisons. The mean difference in percentage point is colour-coded with red indicating the CAF subpopulation to be larger in 4T1 than in 4T07, and vice versa with blue. 128 tumours in total, 4T1 D7 n=21, 4T1 D14 n=24, 4T1 D21 n=24, 4T07 D7 n=24, 4T07 n=21, 4T07 D21 n=14.

| Significantly different CAF subsets D7 |  |  |  |  |  |  |
| --- | --- | --- | --- | --- | --- | --- |
| Markers on CAF subpopulation | CAF subset | P value | Mean % 4T1 | Mean % 4T07 | Difference in percentage point | SE of difference |
| PDPN, FAPa, aSMA, CD26 | S12 | 0.001220 | 1.95 | 0.69 | 1.26 | 0.36 |
| FAPa, CD26 | S24 | 0.001268 | 3.24 | 7.23 | -3.99 | 1.16 |
| CD26 | S32 | 0.000918 | 5.84 | 14.44 | -8.60 | 2.41 |
| PDGFRa, PDGFRb, PDPN, FAPa, aSMA | S33 | 0.000139 | 0.51 | 0.25 | 0.26 | 0.06 |
| PDGFRa, PDGFRb, PDPN, aSMA | S41 | 0.000789 | 3.56 | 1.04 | 2.52 | 0.70 |
| PDGFRb, PDPN, aSMA | S42 | 0.000322 | 2.44 | 0.74 | 1.69 | 0.43 |
| PDGFRa, PDPN, aSMA | S43 | 0.000008 | 0.67 | 0.22 | 0.45 | 0.09 |
| PDPN, aSMA | S44 | 0.000001 | 0.81 | 0.26 | 0.55 | 0.10 |
| PDGFRa, PDGFRb, PDPN, FAPa | S49 | 0.001231 | 1.28 | 0.77 | 0.51 | 0.15 |

| Significantly different CAF subsets D14 |  |  |  |  |  |  |
| --- | --- | --- | --- | --- | --- | --- |
| Markers on CAF subpopulation | CAF subset | P value | Mean % 4T1 | Mean % 4T07 | Difference in percentage point | SE of difference |
| PDGFRb, PDPN, aSMA, CD26 | S10 | 0.000009 | 3.84 | 0.56 | 3.28 | 0.65 |
| PDPN, FAPa, aSMA, CD26 | S12 | 0.000014 | 2.07 | 0.36 | 1.70 | 0.35 |
| PDGFRb, aSMA, CD26 | S14 | 0.000159 | 0.05 | 0.01 | 0.04 | 0.01 |
| PDGFRa, FAPa, CD26 | S23 | 0.000080 | 0.17 | 0.53 | -0.36 | 0.08 |
| PDGFRb, PDPN, CD26 | S26 | 0.000081 | 0.85 | 0.26 | 0.59 | 0.13 |
| PDGFRb, PDPN, aSMA | S42 | 0.001271 | 2.72 | 0.55 | 2.18 | 0.63 |
| PDGFRb, aSMA | S46 | 0.000007 | 0.45 | 0.06 | 0.39 | 0.08 |
| aSMA | S48 | 0.000469 | 0.90 | 0.29 | 0.61 | 0.16 |
| PDGFRa, PDGFRb, FAPa | S53 | 0.001104 | 0.09 | 0.24 | -0.15 | 0.04 |
| PDGFRb, PDPN | S58 | 0.001065 | 0.67 | 0.23 | 0.43 | 0.12 |

| Significantly different CAF subsets D21 |  |  |  |  |  |  |
| --- | --- | --- | --- | --- | --- | --- |
| Markers on CAF subpopulation | CAF subset | P value | Mean % 4T1 | Mean % 4T07 | Difference in percentage point | SE of difference |
| PDGFRb, PDPN, FAPa, aSMA, CD26 | S2 | 0.000003 | 1.97 | 0.39 | 1.58 | 0.29 |
| PDPN, FAPa, aSMA, CD26 | S12 | 0.000914 | 1.35 | 0.25 | 1.10 | 0.31 |
| PDGFRa, PDGFRb, FAPa, CD26 | S21 | 0.000303 | 0.38 | 1.17 | -0.79 | 0.20 |
| PDGFRa, PDGFRb, FAPa | S53 | 0.000005 | 0.10 | 0.56 | -0.46 | 0.09 |
| PDGFRa, PDGFRb | S61 | 0.000009 | 0.07 | 0.80 | -0.73 | 0.14 |
| PDGFRa | S63 | 0.000849 | 1.45 | 13.69 | -12.24 | 3.36 |

In summary, we find that the CAF population is co-evolving along with the TME. Analysis of the 128 tumours revealed temporal changes in abundance of many CAF subpopulations in both 4T1 and 4T07 tumours, and several of these were common to both tumour types. Two CAF subpopulations were markedly different in abundance when comparing tumour types (S32 and S63), supporting the hypothesis that cancer cell intrinsic factors affect the relative size of some CAF subpopulations.

## Discussion

In this study, we developed a multicolour FCM workflow allowing us to dissect the heterogeneity of CAFs and employ it to show for the first time how 6 CAF markers can be used to detect numerous CAF subpopulations within murine TNBC tumours. We provide evidence for the temporal co-evolution of the CAF population as the tumours mature, observing that the expression of the 6 CAF markers all change over time and the dynamics for each marker differ depending on tumour type and time. Of note, we see a striking shift from a prevalence of PDGFRα+ fibroblasts in healthy mammary fat pads, to PDGFRβ+ CAFs in tumour tissue. Moreover, we also see examples of tumour type affecting the relative size of some of the subpopulations making up the total CAF pool, supporting the hypothesis of cancer cell intrinsic factors being important for shaping the CAF compartment. Lastly we identify CD26 as a novel and abundant CAF marker only matched by FAPα as the most prevalent CAF marker in our murine models of TNBC (see Fig 8 for key findings). By employing a negative selection strategy and focusing on surface markers our approach is readily applicable to other mouse models of solid cancer, and it provides an excellent starting point for functional studies of CAF subpopulations, as well as for the continued search for novel CAF markers.

**Figure 8.**
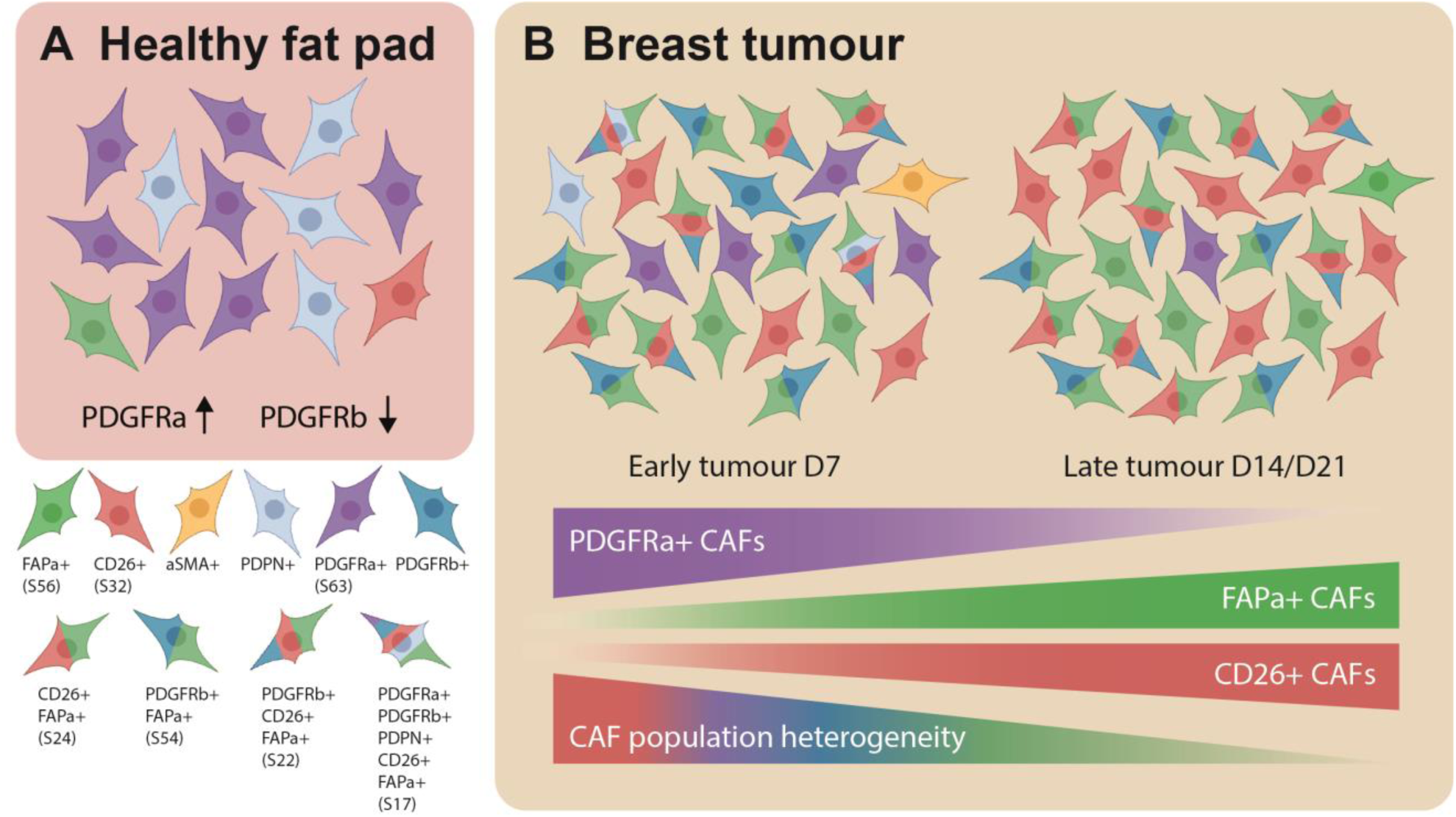
The fibroblast compartment changes dramatically from a healthy fat pad to a breast tumour. In normal mammary fat pads fibroblasts expressing PDGFRα are the most abundant, while PDGFRβ+ fibroblasts are very scarce. Once a tumour forms this picture is reversed. As the tumour progresses the heterogeneity decreases and CAFs expressing combinations of CD26, FAPα and PDGFRβ become the dominant subpopulations. S32= CAF subset 32 etc. see Fig. 4.

### The vast heterogeneity of CAFs

We chose a negative selection approach in order to acquire as diverse a population of fibroblasts/CAFs from mammary fat pads and tumours as possible, and found this Lin-population to be stable over time in tumours (Fig 2). It is important to remember that while the percentages of Lin-cells may remain stable over time, the overall cellularity increases as the tumour grows, primarily due to rapid proliferating cancer cells, and thus the total number of Lin-is increasing drastically. This fact explains why healthy fat pads have a higher percentage of Lin-cells than tumours (10% in healthy FP, 5.5% in 4T1, and 8.7% in 4T07) (Fig 2 & S3), since the fibroblasts are not being ‘diluted’ by cancer cells. The CAF population we detect is slightly larger than previously reported for this tumour model (Avgustinova et al., 2016), however the discrepancy is at least in part explained by the differences in the purification and isolation of single cells, as well as by a substantial disparity in the gating strategies.

The lack of a specific pan CAF marker (Gieniec et al., 2019; LeBleu and Kalluri, 2018) makes it difficult to directly compare the minute results of studies on CAFs, since most research groups have their own definition of what the total CAF population is. The CAF research field in general would benefit immensely from the discovery of pan CAF-specific markers or some consensus marker profiles.

### A switch in PDGFR subtype between normal and tumour tissue

PDGFRα has been reported as a marker of resident mammary fibroblasts (Bartoschek et al., 2018; Raz et al., 2018; Östman, 2017), as a general fibroblast marker (Driskell et al., 2013), and as a CAF marker (Sharon et al., 2013; Öhlund et al., 2017). Our study clearly supports PDGFRα as a marker of resident mammary fibroblasts; we see about 90% PDGFRα+ cells in the CAF+ population (35% in the Lin-population) in normal fat pads, while only around 5% are PDGFRβ+ cells (Fig 3 and S4). On the other hand, as the tumour develops, the tumour becomes highly enriched in PDGFRβ+ CAFs, while dramatically decreasing the numbers of PDGFRα+ CAFs (Fig 3). Our study is consistent with other studies showing a relative loss of PDGFRα+ CAFs during breast cancer progression in mice (Bartoschek et al., 2018; Raz et al., 2018). Our findings support the notion that PDGFRβ may primarily be a marker of CAFs while PDGFRα may mark more resting fibroblasts (Paulsson et al., 2009; Primac et al., 2019; Östman, 2017).

However, PDGFRα seems to be a prominent CAF marker in skin cancer (Arwert et al., 2012; Erez et al., 2010) and recently PDGFRα was suggested as a pan-CAF marker in pancreatic cancer as it was detected in all 4 transcriptomically defined CAF subpopulations (Neuzillet et al., 2019). In a follow-up study of pancreatic cancer, RNA-sequencing discovered that PDPN was a widely expressed CAF marker, and PDGFRα only present on a subpopulation of the CAFs (Elyada et al., 2019). We also see PDPN expression in around 40-50% of CAF+ cells in early tumours (Fig 3), however, there is a significant decrease in PDPN+ CAFs in both 4T1 and 4T07 tumours as they mature, leaving FAPα+ and CD26+ CAFs as more prevalent in late tumours. Combined this suggests that cancer type heavily influences which CAF subpopulations that dominate in the TME. Future FCM studies of CAFs from various cancers will help reveal this, and perhaps point towards the best CAF surface markers for specific cancer types.

### CD26+ CAFs are abundant new members of the CAF family

In our study, we see that FAPα+ CAFs increase over time in both the aggressive 4T1 tumours, and the less aggressive 4T07 tumours, and that the population always cover 40-60% of the total CAF+ population in these tumours (Fig 3 & 4). CD26, also known as DPP4, is a close homologue to FAPα and these two family members can form heterodimers creating complexes with both FAPα and CD26 catalytic activity (Scanlan et al., 1994). CD26 has been reported as a lineage marker in fibrogenic dermal fibroblasts (Rinkevich et al., 2015), but little is known about CD26 in breast cancer. We see a clear population of CD26+ fibroblasts in healthy fat pads, shown previously to be interlobular fibroblasts in human mammary tissue (Morsing et al., 2016), but interestingly the percentage of CD26+ CAFs increases dramatically as tumours start to grow, and remain above normal levels throughout the experiment. We find that CD26 is present on around 60% of all CAF+ cells of both 4T1 and 4T07 tumours if you pool both early and mature tumours (Fig S3). CAF subpopulations only positive for CD26 (S32) or FAPα (S56) and none of the other markers, are two of the most prevalent CAF subpopulations in both 4T1 and 4T07 tumours at all times (Fig 5 and 8), making CD26 as prevalent a CAF marker as FAPα in our murine models of TNBC. Intriguingly, in late stage tumours (D14 and D21), 3 of the 5 most abundant CAF subpopulations in both 4T1 and 4T07 tumours are CD26+ (Fig 5), and we observe a trend towards an increase over time of CD26+ CAFs in 4T1 tumours, and a decrease over time of the same population in the less aggressive 4T07 tumours (Fig 3), warranting further studies into the functional phenotype of these new CD26+ CAF subpopulations.

### Co-existence of numerous CAF subpopulations in breast cancer

Since the initial report on different subpopulations of αSMA+ myofibroblasts in breast cancer (Lazard et al., 1993), it is now becoming clear that CAFs come with different flavours and that a heterogeneous population exists within a single breast tumour (Bartoschek et al., 2018; Busch et al., 2017; Calvo et al., 2013; Costa et al., 2018; Cremasco et al., 2018; Raz et al., 2018; Su et al., 2018). Here, we have presented data showing the existence of numerous CAF subpopulations in murine models of TNBC, expanding on previous findings within breast cancer that have proposed the existence of only 2 to 4 different CAF subpopulations (Bartoschek et al., 2018; Costa et al., 2018; Cremasco et al., 2018; Raz et al., 2018).

Bartoschek et al. propose the existence of at least 3 distinct CAF subpopulations in the MMTV-PyMT mouse model of breast cancer defined by scRNA-seq; vascular (vCAFs), matrix (mCAFs), and developmental (dCAFs) CAFs (Bartoschek et al., 2018). The majority of the analysed CAFs in the MMTV-PyMT model clustered to the vCAFs. These CAFs were by scRNA-seq found to be PDGFRα^-^PDGFRβ^+^PDPN^-^FAPα^+/-^αSMA^+/-^, but the actual expression of the proteins was not analysed, precluding a direct comparison to the abundance of our FCM defined CAF subpopulations. Importantly, the scRNA-seq study of the MMTV-PyMT breast tumours did not mention CD26 as a marker, while we see this marker present on a large proportion of CAFs in our model. This discrepancy could be due to the different murine breast cancer models, i.e. spontaneous MMTV-PyMT vs implantable 4T1/4T07, but it also shows the potential limitation of relying solely on RNA detection. Combining multicolour FCM studies with scRNA-seq and/or proteomics would provide a more solid basis for both discovering new potential markers/targets (via scRNA-seq and proteomics), and validating the cellular presence of these (via FCM).

### Temporal co-evolution of the CAF population as tumours grow

The TME changes as tumours progress, supporting the cancer cells in overcoming new challenges such as securing sufficient blood supply or keeping the immune cells at bay. CAFs have been implicated in many of these tumour-promoting processes (Kalluri, 2016; LeBleu and Kalluri, 2018), yet reports studying the temporal co-evolution of CAFs as tumours progress are very limited. By comparing transcriptomic profiles from breast cancer CAFs and normal mammary fibroblasts Busch et al. suggest that CAFs gradually arise from normal fibroblasts via a primed state, as they find that a part of the fibroblasts isolated from histologically normal lumpectomy resection areas cluster together with CAFs (Busch et al., 2017). A study in the MMTV-PyMT breast cancer model briefly touched upon the concept of CAF co-evolution, and found that fibroblasts isolated from progressive stages of tumour development showed increasing extracellular matrix (ECM) remodelling abilities (Calvo et al., 2013). The remarkable expansion from healthy fad pads to D7 tumours of fibroblasts positive for PDGFRβ and FAPα attests to this cancer cell-driven co-evolution. While our implantable TNBC model naturally skips the pre-malignant stages of tumour development, it offers the possibility to focus on faster temporal co-evolution dynamics occurring as tumours take hold (D7), and grow (D14 and D21).

By comparing all possible Boolean CAF subpopulations, we reveal that early stage tumours differ significantly from late stage tumours in the composition of their CAF population. This was true both for 4T1 and 4T07 tumours, hinting at massive early recruitment/education of CAFs in our models of TNBC (Fig 7). This is further supported by the substantial transformations seen between a healthy mammary fat pad and an early D7 tumour (Fig 2 & 3). Interestingly, when going deeper into those differences, i.e. which CAF subpopulations that change in relative abundance over time, many of these changes are the same in both tumour types (Table 1), suggesting a common way of educating/recruiting the core population of CAFs as TNBC tumours mature and grow in size. In general, very few subpopulations differ between the two tumour types, and most of these changes occur in subpopulations of low abundance (Table 2). The small degree of disparity between the two cell lines may not be that surprising, as they share the same ancestry (Aslakson and Miller, 1992). Nevertheless, two CAF subpopulations do show large differences; CD26 single positive CAFs (S32) are almost 2.5 times as abundant early on in 4T07 tumours (D7), and in late stage tumours (D21) PDGFRα single positive CAFs (S63) are more than 9 times as abundant in 4T07 compared to 4T1 tumours (Table 2). This shows that cancer cell intrinsic factors do affect the size of CAF subpopulations, and it is interesting to speculate whether the differences reflects small but important dissimilarities in the ratio of education of resident fibroblasts vs. de novo recruitment of e.g. BM-derived CAF precursors. Further studies will help figure out if the subpopulations that differ between 4T1 and 4T07 tumours have functions that contribute to the metastatic potential of the tumour; e.g. whether CD26+ CAFs are generating an immune suppressive TME in TNBC as suggested previously (Costa et al., 2018). A remaining open question is whether PDGFRα single positive CAFs in 4T07 are resident fibroblast that have yet to become fully activated CAFs, or maybe represent a CAF population with anti-tumorigenic properties?

## Conclusion

In conclusion, we are the first to use an FCM strategy to extensively investigate CAF heterogeneity and the temporal dynamics of CAF markers in TNBC progression. Our study adds to the body of growing evidence against a simplistic single-marker definition of CAFs by identifying the co-existence of multiple CAF subpopulations in TNBC based on the detection of 6 CAF markers. While a few CAF subpopulations dominate the TME and are common to both 4T1 and 4T07 tumours, many subtler subpopulations are nevertheless detectable and may have important functions. CD26 has emerged as the “new kid on the fibroblast marker block” in murine TNBC, and future studies should dissect the function of CD26+ CAFs and other prominent subpopulations.

## Material and methods

### Mouse model of TNBC

All animal experiments were approved by the Danish Animal Experiments Inspectorate, following the guidelines in the EU Directive 2010/63/EU.

#### Breast cancer cell lines

The mouse breast cancer cell lines 4T1 (RRID:CVCL_0125) and 4T07 (RRID:CVCL_B383) are both derived from the same, spontaneous tumour from a BALB/c mouse (Aslakson and Miller, 1992). Both cancer cell lines were acquired from the Karmanos Cancer Institute, USA, and cultured in DMEM w/L-Glutamine, high glucose and pyruvate, + 10% FBS+ 1% (100U/ml) P/S (Gibco, ThermoFisher Scientific, Copenhagen, Denmark) in a 37°C humid incubator with 5% CO2. The cells were split 1:10 when they reached 70-80% confluency. Cell lines were routinely checked to be free of mycoplasma and cell line purity and identity was validated by short tandem repeats (STR)-profiling at the start of the project.

#### αSMA-RFP mice

To facilitate detection of the intracellular cytoskeletal protein αSMA in living cells such as CAFs, we used BALB/c mice genetically engineered to express a red fluorescent protein (RFP) simultaneously as the αSMA protein. These BALB/c αSMA-RFP transgenic mice express the DsRed-Express RFP protein under the mouse *Acta2* gene promoter (LeBleu et al., 2013). These mice were obtained from Raghu Kalluri and bred hemizygously in-house at our institute, but are now commercially available from The Jackson Laboratories (RRID:IMSR_JAX:031160).

#### Orthotopic breast tumour implantation model

When preparing cells for orthotopic implantation, the trypsinised cells were washed in PBS (Gibco, ThermoFisher Scientific, Copenhagen, Denmark) to remove remaining media and trypsin, and then re-suspended in PBS at 10^7^ cells/ml. The cell suspension was kept on ice until injection. 50µl cell suspension was injected into the lower left and right mammary fat pad of either 8-16 weeks old wild type BALB/c mice (for FCM control purposes) or hemizygous αSMA-RFP mice, using a 27G disposable needle, depositing 5×10^5^ cells per injection. Genetically identical cells, i.e. either 4T1 or 4T07, were implanted in both sides of each mouse in order to minimise the total number of mice. Tumour growth and animal welfare was monitored twice a week, following regulations stipulated by the Danish Animal Experiments Inspectorate. At 7, 14 or 21 days (D7, D14, D21) post injection the resulting primary tumours and remaining surrounding fat pad were collected in PBS on ice after euthanasia of the animals, and single cell suspensions prepared as described below.

#### Sample size

Three independent biological repeats of the orthotopic tumour models were carried out, and the sample size (number of tumours = technical repeats) within each biological repeat is listed below. Additionally, 12 healthy mammary fat pads were also collected and analysed in the same way as the tumour samples.

|  | 4T1 |  |  |  | 4T07 |  |  |  |  | Healthy mammary fat pad |  |  |
| --- | --- | --- | --- | --- | --- | --- | --- | --- | --- | --- | --- | --- |
|  | Rep 1 | Rep 2 | Rep 3 | Total | Rep 1 | Rep 2 | Rep 3 | Total |  | Rep 2 | Rep 3 | Total |
| D7 | 7 | 7 | 7 | 21 | 8 | 8 | 8 | 24 | D0 | 8 | 4 | 12 |
| D14 | 8 | 8 | 8 | 24 | 6 | 8 | 7 | 21 |  |  |  |  |
| D21 | 8 | 8 | 8 | 24 | 5 | 5 | 4 | 14 |  |  |  |  |
| Total |  |  |  | 69 |  |  |  | 59 |  |  |  | 12 |

Sample size was determined by the maximum number of samples it was possible to process in each biological repeat. Hemizygous aSMA-RFP mice were randomly allocated to be part of either the 4T1 or the 4T07 group, and injected with the respective tumour cells. On each collection day four animals from each tumour group were randomly selected for euthanasia and subsequent tumour collection. Due to paucity of cells in some tumour samples, the final number of tumours (technical repeats) analysed varies from 4 to the planned maximum of 8, with a total of 128 tumours analysed. The analysis of the tumour samples was not blinded.

### Flow cytometry

#### Dissociation of tumours into single cells

Tumours and cell suspensions were kept on ice between steps. Tissue was minced into roughly 2×2 mm pieces using disposable scalpels, and treated with the digestion enzyme mix from the mouse tumour dissociation kit by Miltenyi (Miltenyi Biotec Norden AB, Lund, Sweden, cat. # 130-096-730). Following the directions on the kit, the sample was then incubated in c-tubes (Miltenyi Biotec Norden AB, Lund, Sweden, cat. # 130-096-334) on the gentleMACS Octo tissue homogeniser w/ heaters (Miltenyi Biotec Norden AB, Lund, Sweden) to keep the mixture at 37°C, using the pre-defined tumour_TDK2 program, running for 41 min. The sample was then washed with PBS and strained through a 70-micron mesh strainer to obtain a single cell suspension. Red blood cells (RBCs) were lysed using 1x RBC lysis solution from BD (Becton Dickinson Denmark A/S, Lyngby, Denmark, cat. # 555899), and cellular debris was removed according to directions in the Miltenyi Debris Removal Kit (Miltenyi Biotec Norden AB, Lund, Sweden, cat. # 130-109-398). The final single cell suspension was frozen in freezing media containing 50% DMEM 40% FBS and 10% DMSO, and kept frozen until the day of FCM analysis.

#### Sample preparation and antibody labelling

To minimize the technical noise and differences in antibody labelling, all frozen single cell suspensions of 4T1 and 4T07 tumours from a biological repeat were thawed and prepped for FCM analysis on the same day. For all washing steps and sample suspension, cold FACS buffer containing PBS + 2mM EDTA + 1% BSA + 25mM HEPES, pH 7 was used unless otherwise noted. Thawed samples were counted and a maximum of 10^7^ cells resuspended in 100µl PBS and incubated on ice for 20 min with 1µl Viobility-405/520 amine reactive viability dye (Miltenyi Biotec Norden AB, Lund, Sweden, cat. # 130-110-206) per 100µl cell suspension. Excess viability dye was washed off using FACS buffer, and samples were incubated in 200µl biotin labelled lineage marker antibody cocktail for 30 min at 4°C followed by washing in FACS buffer. Lastly, samples were incubated 30 min in the dark at 4°C in 100µl CAF marker antibody cocktail per 2×10^6^ cells, then washed 3 times in FACS buffer and kept in the dark on ice until acquisition on the BD LSRII flow cytometer. The CAF marker antibody cocktail included the streptavidin-BV711 to simultaneous detect the biotin-labelled lineage positive cells.

### Reagents and multicolour FCM panel

#### Lineage marker cocktail

The lineage cocktail consisted of biotin labelled, primary antibodies detecting endothelial cells (CD31), immune cells (CD45), basal myoepithelial cells of the mammary ducts (CD49f) (Koukoulis et al., 1991; Villadsen et al., 2007), lymphatic endothelial cells (LYVE-1), and erythroid lineage cells (TER-119). Cancer cells were excluded using EpCAM/CD236, CD24 and CD49f.

**Reagent Table 1.** Lineage marker antibody cocktail.

| Lineage marker antibody cocktail |  |  |  |  |  |
| --- | --- | --- | --- | --- | --- |
| Antigen | Clone | Conjugation | Dilution | Supplier | Cat. # |
| CD24 | M1/69 | Biotin | 1:10 | Miltenyi | 130-101-982 |
| CD31 | 390 | Biotin | 1:10 | Miltenyi | 130-101-955 |
| CD45.2 | 104 | Biotin | 1:100 | BioLegend | 109804 |
| CD49f/Itga6 | GoH3 | Biotin | 1:100 | BioLegend | 313604 |
| EpCam/CD326 | G8.8 | Biotin | 1:50 | BioLegend | 118204 |
| TER-119 | Ter-119 | Biotin | 1:10 | Miltenyi | 130-101-882 |
| LYVE-1 | ALY7 | Biotin | 1:100 | eBioscience | 13-0443-82 |

**Reagent Table 2.** CAF marker antibody cocktail.

| CAF marker antibodies and multi colour FCM panel |  |  |  |  |  | Excitation laser | Filter set |  |
| --- | --- | --- | --- | --- | --- | --- | --- | --- |
| Cellular marker/<br>Antigen | Clone | Conjugation/<br>Fluorophore | Dilution | Supplier | Cat # |  | LP | BP |
| CD140a / PDGFRa | APA5 | BV421 | 1:25 | BD Bioscience | 562774 | 405nm | - | 405/50 |
| CD140b / PDGFRb | REA634 | APC-Vio770 | 1:25 | Miltenyi | 130-109-870 | 633nm | 755 | 780/60 |
| FAPa | polyclonal | AF488 | 1:50 | Nordic Biosite | bs-5758R-A488 | 488nm | 505 | 530/30 |
| Podoplanin | 8.1.1 | APC | 1:50 | SONY | 1237050 | 633nm | - | 660/20 |
| CD26 / DPP4 | H194-112 | PE-Vio770 | 1:25 | Miltenyi | 130-106-322 | 488nm | 755 | 780/60 |
| aSMA<sup>#</sup> | - | DsRed-Express | - | - | - | 488nm | 550 | 575/26 |
| Lineage markers (Biotin) | Streptavidin | BV711 | 1:500 | BD Bioscience | 563262 | 633nm | 690 | 720/30 |
| Viability dye | - | Viability 405/520 | - | Miltenyi | 130-110-206 | 405nm | 475 | 525/50 |

<sup>#</sup> DsRed is ectopically expressed in cells also expressing αSMA, no antibody needed.
The Streptavidin is included in this table as it was admixed with the CAF marker antibodies to a final cocktail applied to the samples after incubation with the lineage marker cocktail. LP= long pass filter, BP = band pass filter.

#### Data acquisition

All FCM data were acquired on a BD LSRII analyser using DIVA software. The fluorescent antibodies on the cells were excited by either a 405nm, 488nm or a 633nm laser in the LSRII analyser. BD cytometer setup and tracking (CS&T) beads were run daily on the analyser to calibrate the laser setup. The PMT voltages were checked before each run, and adjusted if necessary to assure that the positive population of the single stained controls were within detection range.

#### Compensation controls

For compensation purposes, at least 5000 positive events were acquired from single colour controls, one for each fluorophore. These single stained controls were prepared by adding the fluorophore labelled antibodies to a drop of UltraComp Beads (eBioscience, ThermoFisher Scientific, Copenhagen, Denmark, cat # 01-2222-42), followed by 30 min incubation in the dark at 4°C, and subsequent washing with FACS buffer. An exception to this was the CD140b-APC-Vio770 antibody, which does not bind to the UltraComp beads, and thus here an aliquot of single cell suspension from a tumour grown in a wild type (RFP^-/-^) mouse was used instead of the beads. Fibroblasts isolated from a healthy hemizygous αSMA-RFP mouse were grown on plastic to induce αSMA expression, and used as single colour control for the DsRed fluorophore. A sample of a tumour grown in a wild type (RFP^-/-^) mouse was used as a negative control for the APC-Vio770 antibody and the DsRed fluorophore, while the other single colour controls on the UltraComp beads carried their own internal negative control in the form of beads not binding to any antibodies. A compensation matrix was created based on the compensation controls, applied to the FCS files in the FCS Express 6 software, and compensated FCS files were used for all subsequent gating and analysis.

### FCM data analysis

All FCS files were analysed using the FCS Express 6 software.

#### Gating controls

No isotype controls were used in this study as this type of control has very limited usage in helping proper gating in multicolour FCM, where most of the background is due to spill-over from fluorophores with overlapping spectra (Maecker and Trotter, 2006). Fluorescent minus one (FMO) controls were used for each fluorophore to help guide proper gating on each marker (Roederer, 2002) (see Fig S2). The FMOs were prepared by staining tumour samples with a mix of all the conjugated-antibodies within the panel except one antibody, ending up with multiple FMO tubes, each tube lacking a specific fluorophore. The gates for each of the CAF markers was set so no more than 0.03% of the events in the FMO control sample fell within the gate.

#### Gating strategy

Cells were gated on forward and side scatter (FSC and SSC), and doublets were excluded by both FSC-H vs FSC-W and SSC-H vs SSC-W bivariate plots. Live cells were gated as Viobility 405/20-cells, and Lineage negative (Lin-) cells were gated on Live cells as BV711-cells (EpCAM-CD45-CD31-TERT119-CD24-CD49f- and LYVE-1-). CAF+ cells were then gated on Lin-cells as cells positive for at least one of the 6 CAF markers using Boolean criteria (αSMA+ OR FAPα+ OR CD26+ OR PDGFRα+ OR PDGFRβ+ OR PDPN+). See Fig 1B for visual representation of gating strategy.

#### FCM data visualisation

Percentages from Boolean gating on the 6 CAF markers performed in FCS express 6 was exported to an excel spreadsheet and visualised using the freeware SPICE (https://niaid.github.io/spice/) (Roederer et al., 2011). Each of the possible Boolean combinations (Boolean CAF subpopulations) were plotted as slices in a pie chart, with the arcs around the pie charts corresponding to one CAF marker (Fig 7). All samples within a group defined by tumour and day, e.g. 4T1 D7, were grouped together and averaged (relative scaling), so the pie slices show the mean percentage of each CAF subpopulation within the total CAF+ population. The heat-map over the ‘Boolean CAF subpopulation abundance’ across time and tumour type (Fig 4) was created using the clustering freeware TreeView3.0 (https://bitbucket.org/TreeView3Dev/treeview3/src/master/). All bar plots, graphs, and box and whisker plots were created using Graph Pad Prism version 6.

### UMAP analysis

The intensity levels of each of the 6 CAF markers were normalised into a range from 0=no signal to 1=strongest signal. The UMAP dimension reduction of the Lin-population was performed as previously described (https://arxiv.org/abs/1802.03426) (McInnes et al., 2018).

### Statistics

All statistical tests were performed using Graph Pad Prism version 6. When comparing CAF populations across time within each tumour type (intra-tumour time differences) 2-way ANOVA was used to analyse the dataset, followed by Tukey’s multiple comparisons post-test with statistical significance set to α=0,05. In case the 2-way ANOVA found tumour type not to be a significant variable, all tumours were pooled and analysed by one-way ANOVA to assess the impact of time and test for a possible linear trend. Multiple unpaired, two-tailed t-tests without assuming equal variance were used to compare time points across tumour type (inter-tumour differences), using the false discovery rate (FDR) approach with false discovery rate Q=1% to correct for multiple comparisons. When comparing pie charts generated in SPICE a build-in permutation test was used. The permutation test asks how often, given the samples that comprise the compared pies, the difference observed would happen simply by chance (https://niaid.github.io/spice/help/analysis-comparingoverlays).

## List of Abbreviations

ANOVA: Analysis of variance
aSMA: Alpha-smooth muscle actin
BM: Bone marrow
CAF(s): Cancer-associated fibroblast(s)
CB95: 95% confidence band
CD49f: Integrin alpha 6 / Itga6
CI95: 95% confidence interval
CTCs: Circulating tumour cells
D14: Day 14
D21: Day 21
D7: Day 7
DPPVI: Di-peptidyl peptidase 4 / CD26
ECM: Extracellular Matrix
eGFP: Enhanced green fluorescent protein
EpCAM: Epithelial cell adhesion molecule
FACS: Fluorescent activated cell sorting
FAPa: Fibroblast activation protein alpha
FCM: Flow cytometry
FDR: False discovery rate
FSC: Forward scatter
FSP1: Fibroblast specific protein 1 / S100A4
Lin-: Lineage negative (devoid of lineage markers)
MMTV-PyMT: Mouse mammary tumour virus Polyoma middle T-antigen
MSC(s): Mesenchymal stromal cell(s)
PDGFRa: Platelet derived growth factor receptor alpha / CD140a
PDGFRb: Platelet derived growth factor receptor beta / CD140b
PDPN: Podoplanin
PMT(s): Photomultiplier tubes(s)
RFP: Red fluorescent protein
scRNA-seq: single cell RNA sequencing
SEM: Standard error of the mean
SSC: Side scatter
TME: Tumour microenvironment
TNBC: Triple-negative breast cancer
UMAP: Uniform Manifold Approximation and Projection

## Author contributions

F.A.V. conceived the study, planned, and carried out all the experiments and performed the majority of the data analysis. K.W.Z. and L.W. helped with the *in vivo* experiments, with the preparation of tumours into FCM ready samples, and edited the manuscript. M.C.H. and K.J.W. analysed the data and produced the UMAP plots. F.A.V. wrote the manuscript. J.T.E. and C.D.M. supervised the study, helping to conceive and plan the study, as well as editing the manuscript.

