## Supplementary figures and tables for "Deciphering the temporal heterogeneity of cancer-associated fibroblast subpopulations in breast cancer"

### Supplementary material

#### Supplementary Figures

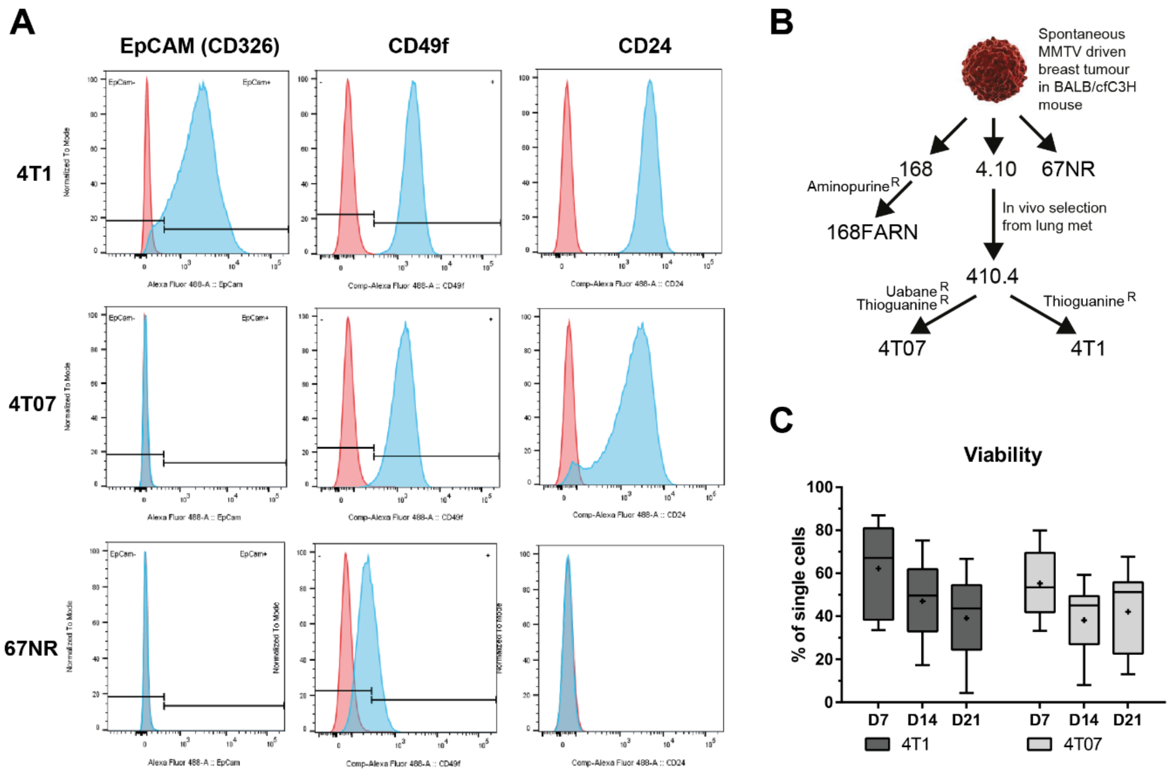

**Supplementary Figure 1. Cell surface marker analysis of mouse breast cancer cell lines. A**) Cell surface marker FCM analysis of 4T1, 4T07 and 67NR cell lines. The red graph shows the unstained control and blue the stained cell sample, n=1. **B**) Derivation of triple-negative breast cancer cell lines from a single MMTV-driven spontaneous tumour. **C**) Viability of tumour single cell suspensions across time and tumour type, all three repeats combined, Tukey style box and whisker plots with line at the median and '+' at the mean. Initially the plan was to use the 4T1 and 67NR cell lines for investigating the relation between CAF subpopulations and the metastatic potential of tumours as 67NR do not metastasise at all, however we could not find a suitable cell surface marker present on 67NR cells that was not also likely present on fibroblasts. We thus chose to use 4T07 cells which are less metastatic than 4T1 cells, albeit more metastatic than 67NR cells (Aslakson and Miller, 1992).

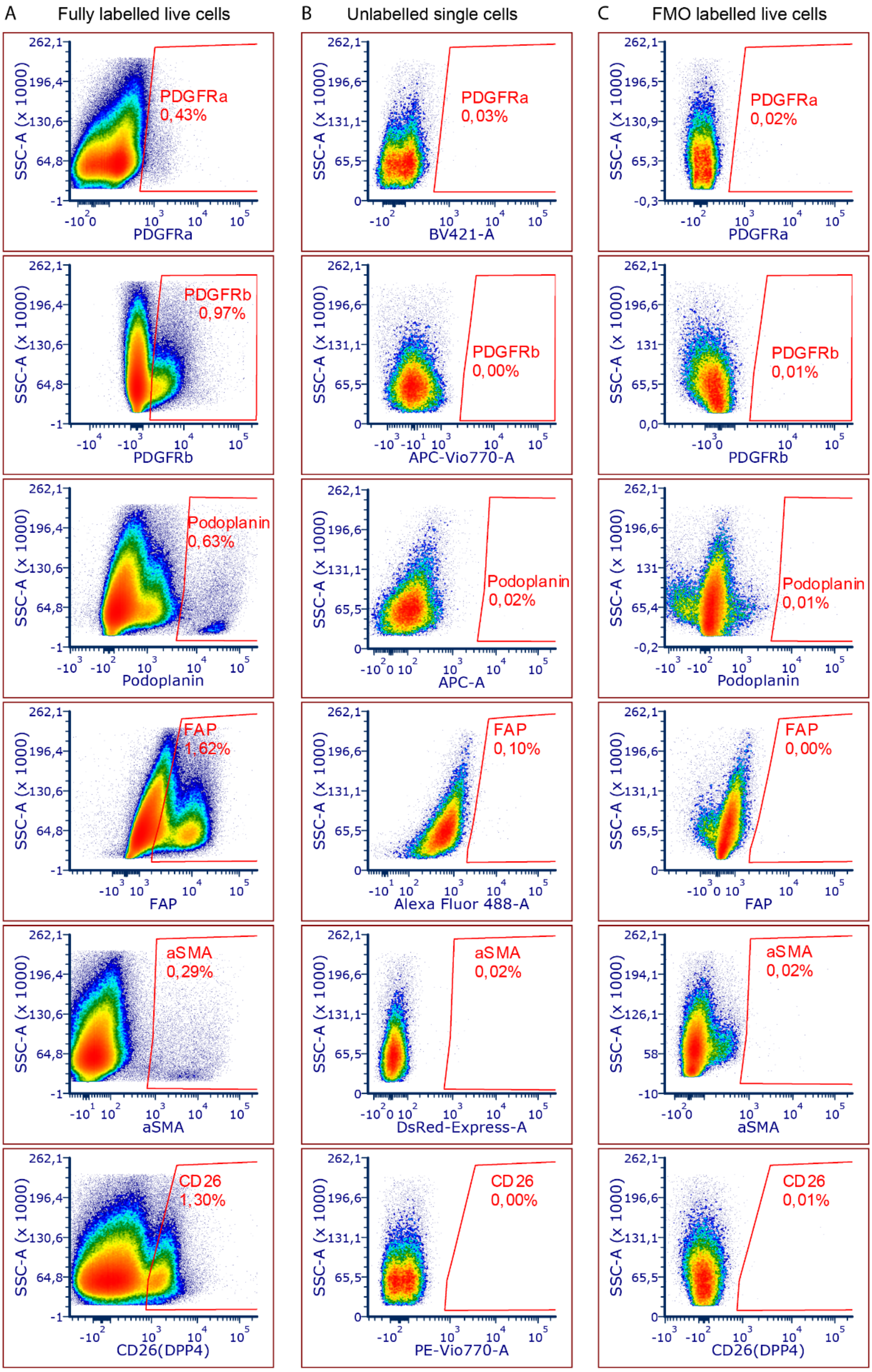

**Supplementary Figure 2. Setting of CAF marker gates using fluorescent minus one (FMO) controls. A**) CAF marker gates shown on a fully stained tumour sample, and **B**) the same gates shown on an unlabelled tumour sample. **C**) The gates are set on a sample labelled with all but the one marker the gates is being set for (FMO labelled), and placed so that less than 0,03% percent of the events fall within the gate. For each of the three repeats FMO controls were prepared fresh and used to set the CAF marker gates, here an example from repeat 3 (LSR06) is shown.

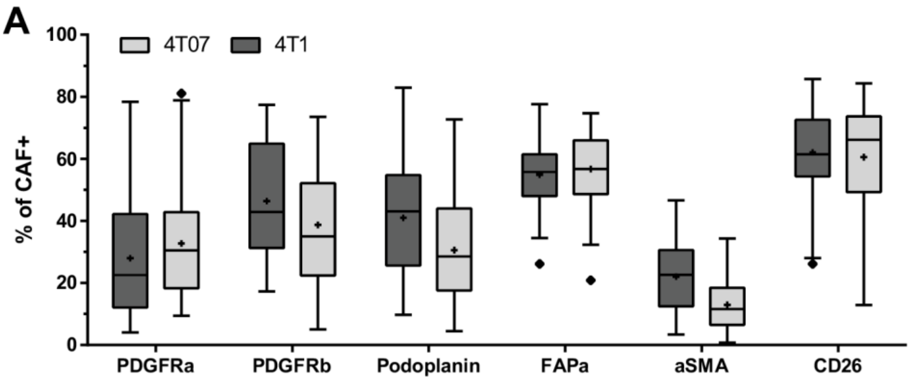

| | 4T1 | | | 4T07 | | |
|---|---|---|---|---|---|---|
| | Median | Upper Limit | Lower Limit | Median | Upper Limit | Lower Limit |
| PDGFRa | 22.5 | 78.4 | 4.1 | 30.5 | 81.1 | 9.4 |
| PDGFRb | 42.8 | 77.4 | 17.2 | 35.0 | 73.5 | 5.1 |
| Podoplanin | 43.0 | 83.0 | 9.7 | 28.6 | 72.7 | 4.5 |
| FAPa | 55.8 | 77.6 | 26.1 | 56.7 | 74.7 | 20.8 |
| aSMA | 22.7 | 46.6 | 3.4 | 11.5 | 34.3 | 0.8 |
| CD26 | 61.5 | 85.7 | 26.1 | 66.2 | 84.3 | 12.8 |

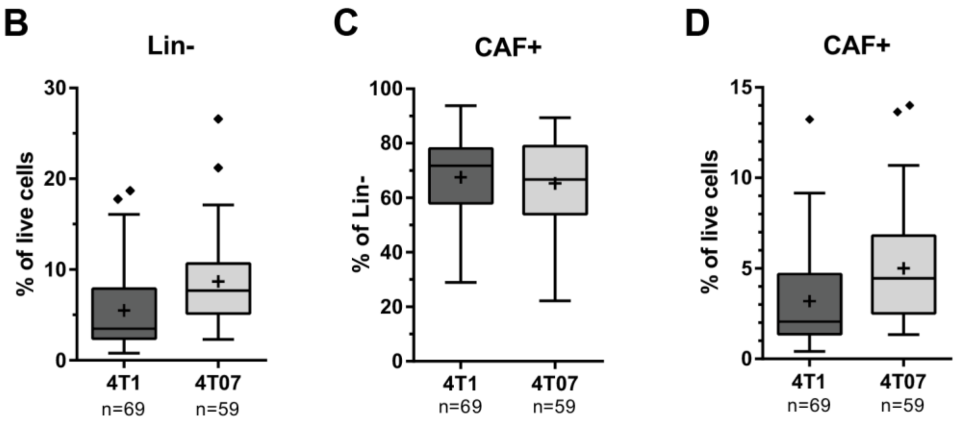

**Supplementary Figure 3. Cell population sizes across tumour type without time dimension to see biological variance.** A) Lin- population out of live cells, B) CAF+ population out of Lin- cells, C) CAF+ out of live cells. D) Percentage of CAF+ cells expressing the respective CAF marker. All plots show three independent repeats combined as Tukey style box and whisker plots, with a line denoting the median and a '+'denoting the mean.

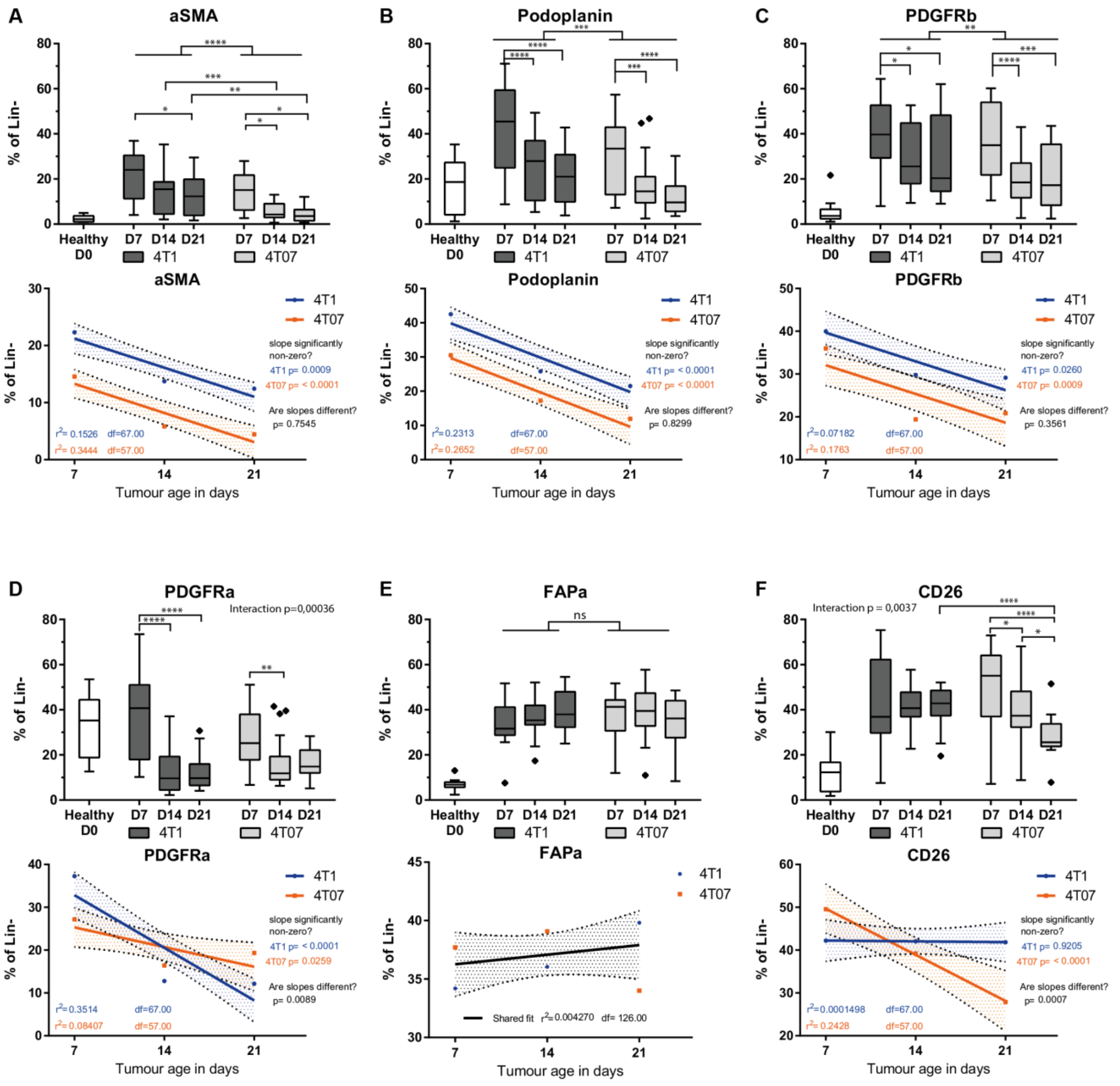

**Supplementary Figure 4. CAF marker dynamics across time and tumour type within the Lin- population. A-F)** Percent of cells positive for the respective CAF marker in healthy mammary fat pad (D0) and 4T1 and 4T07 tumours across the different time points. Tukey box plots of two (healthy samples) or three independent repeats (tumours samples) combined. The boundaries of the Tukey style box goes from the 25$^{th}$ to the 75$^{th}$ percentile, and the median is depicted by a line. The whiskers extend from the value closest to the 25$^{th}$ percentile minus 1.5 times the interquartile range (IQR = difference between 25$^{th}$ to 75$^{th}$ percentile), to the value closest to the 75$^{th}$ percentile plus 1.5 x IQR. Any values outside this range is plotted as individual points. The healthy samples provide a D0 reference point, but all the statistics are only concerning the tumour samples. Fitting a straight line to the data, testing if one line fits both 4T1 and 4T07 tumours to indicate no difference, or if slope and/or y-intercept differ if two lines better capture the dataset. Fitted lines are shown with hashed out 95% confidence bands (CB95), with dots indicating the observed mean at the respective time point. 2-way ANOVA was run once for each marker separately to look for interaction and over-all effect of tumour type and/or day. To determine statistical significance between time points within each tumour type (intra-tumour time comparisons) a 2-way ANOVA with Tukey's multiple comparisons post-test was run on all the CAF markers combined, once for 4T1 tumours and once for 4T07 tumours. To determine if markers differed between tumour types, multiple unpaired, two-tailed t-tests without assuming equal variance and with FDR correction (Q=1%) was run for each of the three time points (inter-tumour comparison). *= p<0.05, **=p<0.01, ***=p<0.001, **** p<0.0001. Healthy n=12, 4T1 D7 n=21, 4T1 D14 n=24, 4T1 D21 n=24, 4T07 D7 n=24, 4T07 n=21, 4T07 D21 n=14.

| Raw percentages 2-way ANOVA | | | | | | Difference between raw and normalized data ? | Normalized to 4T1 D7 mean 2-way ANOVA | | | | | |
|---|---|---|---|---|---|---|---|---|---|---|---|---|
| Tukey's multiple comparisons test | Mean Diff. percentage point | 95% CI of diff, | Signifi-cant? | * | Adjusted P Value | | Tukey's multiple comparisons test | Mean Diff. norm. ratios | 95% CI of diff, | Signifi-cant? | * | Adjusted P Value |
| **4T1** | | | | | | | **4T1** | | | | | |
| **PDGFRa** | | | | | | | **PDGFRa** | | | | | |
| 4T1Day 7 vs. 4T1day 14 | 29.72 | 20,23 to 39,22 | Yes | **** | < 0,0001 | | 4T1Day 7 vs. 4T1day 14 | 0.65 | 0,4848 to 0,8072 | Yes | **** | < 0,0001 |
| 4T1Day 7 vs. 4T1day 21 | 30.85 | 21,35 to 40,34 | Yes | **** | < 0,0001 | | 4T1Day 7 vs. 4T1day 21 | 0.63 | 0,4673 to 0,7897 | Yes | **** | < 0,0001 |
| 4T1day 14 vs. 4T1day 21 | 1.12 | -8,047 to 10,29 | No | ns | 0.9552 | | 4T1day 14 vs. 4T1day 21 | -0.02 | -0,1732 to 0,1382 | No | ns | 0.9622 |
| **PDGFRb** | | | | | | | **PDGFRb** | | | | | |
| 4T1Day 7 vs. 4T1day 14 | 9.55 | 0,05991 to 19,05 | Yes | * | 0.0482 | | 4T1Day 7 vs. 4T1day 14 | 0.20 | 0,04247 to 0,3649 | Yes | ** | 0.0088 |
| 4T1Day 7 vs. 4T1day 21 | 10.50 | 1,011 to 20,00 | Yes | * | 0.0259 | | 4T1Day 7 vs. 4T1day 21 | 0.23 | 0,06594 to 0,3883 | Yes | ** | 0.0029 |
| 4T1day 14 vs. 4T1day 21 | 0.95 | -8,219 to 10,12 | No | ns | 0.9677 | | 4T1day 14 vs. 4T1day 21 | 0.02 | -0,1323 to 0,1792 | No | ns | 0.9331 |
| **Podoplanin** | | | | | | | **Podoplanin** | | | | | |
| 4T1Day 7 vs. 4T1day 14 | 17.27 | 7,772 to 26,76 | Yes | **** | < 0,0001 | | 4T1Day 7 vs. 4T1day 14 | 0.34 | 0,1831 to 0,5055 | Yes | **** | < 0,0001 |
| 4T1Day 7 vs. 4T1day 21 | 23.78 | 14,29 to 33,27 | Yes | **** | < 0,0001 | | 4T1Day 7 vs. 4T1day 21 | 0.44 | 0,2836 to 0,6059 | Yes | **** | < 0,0001 |
| 4T1day 14 vs. 4T1day 21 | 6.52 | -2,656 to 15,69 | No | ns | 0.2175 | | 4T1day 14 vs. 4T1day 21 | 0.10 | -0,05530 to 0,2562 | No | ns | 0.2839 |
| **FAPa** | | | | | | | **FAPa** | | | | | |
| 4T1Day 7 vs. 4T1day 14 | -8.43 | -17,92 to 1,067 | No | ns | 0.0936 | yes | 4T1Day 7 vs. 4T1day 14 | -0.21 | -0,3662 to -0,04386 | Yes | ** | 0.0082 |
| 4T1Day 7 vs. 4T1day 21 | -15.40 | -24,89 to -5,905 | Yes | *** | 0.0005 | | 4T1Day 7 vs. 4T1day 21 | -0.36 | -0,5261 to -0,2037 | Yes | **** | < 0,0001 |
| 4T1day 14 vs. 4T1day 21 | -6.97 | -16,14 to 2,199 | No | ns | 0.1748 | yes | 4T1day 14 vs. 4T1day 21 | -0.16 | -0,3156 to -0,004108 | Yes | * | 0.0427 |
| **aSMA** | | | | | | | **aSMA** | | | | | |
| 4T1Day 7 vs. 4T1day 14 | 9.11 | -0,3859 to 18,60 | No | ns | 0.0632 | yes | 4T1Day 7 vs. 4T1day 14 | 0.36 | 0,1987 to 0,5210 | Yes | **** | < 0,0001 |
| 4T1Day 7 vs. 4T1day 21 | 11.10 | 1,603 to 20,59 | Yes | * | 0.0171 | | 4T1Day 7 vs. 4T1day 21 | 0.42 | 0,2622 to 0,5846 | Yes | **** | < 0,0001 |
| 4T1day 14 vs. 4T1day 21 | 1.99 | -7,182 to 11,16 | No | ns | 0.8664 | | 4T1day 14 vs. 4T1day 21 | 0.06 | -0,09214 to 0,2193 | No | ns | 0.6022 |
| **CD26** | | | | | | | **CD26** | | | | | |
| 4T1Day 7 vs. 4T1day 14 | -8.33 | -17,82 to 1,166 | No | ns | 0.0989 | yes | 4T1Day 7 vs. 4T1day 14 | -0.20 | -0,3607 to -0,03828 | Yes | * | 0.0106 |
| 4T1Day 7 vs. 4T1day 21 | -9.24 | -18,73 to 0,2539 | No | ns | 0.0584 | yes | 4T1Day 7 vs. 4T1day 21 | -0.24 | -0,4018 to -0,07943 | Yes | ** | 0.0014 |
| 4T1day 14 vs. 4T1day 21 | -0.91 | -10,08 to 8,258 | No | ns | 0.9702 | | 4T1day 14 vs. 4T1day 21 | -0.04 | -0,1969 to 0,1146 | No | ns | 0.8084 |
| **4T07** | | | | | | | **4T07** | | | | | |
| **PDGFRa** | | | | | | | **PDGFRa** | | | | | |
| 4T07 day 7 vs. 4T07 day 14 | 12.25 | 2,537 to 21,97 | Yes | ** | 0.009 | | 4T07 day 7 vs. 4T07 day 14 | 0.25 | 0,02512 to 0,4834 | Yes | * | 0.0254 |
| 4T07 day 7 vs. 4T07 day 21 | 5.10 | -5,840 to 16,03 | No | ns | 0.5167 | | 4T07 day 7 vs. 4T07 day 21 | 0.10 | -0,1592 to 0,3565 | No | ns | 0.6401 |
| 4T07 day 14 vs. 4T07 day 21 | -7.16 | -18,38 to 4,061 | No | ns | 0.2912 | | 4T07 day 14 vs. 4T07 day 21 | -0.16 | -0,4201 to 0,1090 | No | ns | 0.3504 |
| **PDGFRb** | | | | | | | **PDGFRb** | | | | | |
| 4T07 day 7 vs. 4T07 day 14 | 17.08 | 7,361 to 26,79 | Yes | *** | 0.0001 | | 4T07 day 7 vs. 4T07 day 14 | 0.31 | 0,08233 to 0,5406 | Yes | ** | 0.0043 |
| 4T07 day 7 vs. 4T07 day 21 | 15.41 | 4,473 to 26,34 | Yes | ** | 0.0029 | | 4T07 day 7 vs. 4T07 day 21 | 0.31 | 0,04906 to 0,5648 | Yes | * | 0.0148 |
| 4T07 day 14 vs. 4T07 day 21 | -1.67 | -12,89 to 9,550 | No | ns | 0.9346 | | 4T07 day 14 vs. 4T07 day 21 | 0.00 | -0,2691 to 0,2600 | No | ns | 0.9991 |
| **Podoplanin** | | | | | | | **Podoplanin** | | | | | |
| 4T07 day 7 vs. 4T07 day 14 | 14.38 | 4,664 to 24,10 | Yes | ** | 0.0016 | | 4T07 day 7 vs. 4T07 day 14 | 0.26 | 0,02872 to 0,4869 | Yes | * | 0.0229 |
| 4T07 day 7 vs. 4T07 day 21 | 20.38 | 9,446 to 31,32 | Yes | **** | < 0,0001 | | 4T07 day 7 vs. 4T07 day 21 | 0.38 | 0,1239 to 0,6396 | Yes | ** | 0.0016 |
| 4T07 day 14 vs. 4T07 day 21 | 6.00 | -5,218 to 17,22 | No | ns | 0.4194 | | 4T07 day 14 vs. 4T07 day 21 | 0.12 | -0,1407 to 0,3884 | No | ns | 0.5133 |
| **FAP** | | | | | | | **FAP** | | | | | |
| 4T07 day 7 vs. 4T07 day 14 | -10.50 | -20,22 to -0,7858 | Yes | * | 0.0305 | yes | 4T07 day 7 vs. 4T07 day 14 | -0.22 | -0,4513 to 0,006963 | No | ns | 0.0596 |
| 4T07 day 7 vs. 4T07 day 21 | -4.48 | -15,42 to 6,452 | No | ns | 0.5994 | | 4T07 day 7 vs. 4T07 day 21 | -0.13 | -0,3872 to 0,1285 | No | ns | 0.4655 |
| 4T07 day 14 vs. 4T07 day 21 | 6.02 | -5,201 to 17,24 | No | ns | 0.4173 | | 4T07 day 14 vs. 4T07 day 21 | 0.09 | -0,1717 to 0,3574 | No | ns | 0.6872 |
| **aSMA** | | | | | | | **aSMA** | | | | | |
| 4T07 day 7 vs. 4T07 day 14 | 10.36 | 0,6476 to 20,08 | Yes | * | 0.0334 | | 4T07 day 7 vs. 4T07 day 14 | 0.34 | 0,1111 to 0,5694 | Yes | ** | 0.0016 |
| 4T07 day 7 vs. 4T07 day 21 | 11.80 | 0,8686 to 22,74 | Yes | * | 0.0308 | | 4T07 day 7 vs. 4T07 day 21 | 0.42 | 0,1609 to 0,6766 | Yes | *** | 0.0005 |
| 4T07 day 14 vs. 4T07 day 21 | 1.44 | -9,779 to 12,66 | No | ns | 0.9509 | | 4T07 day 14 vs. 4T07 day 21 | 0.08 | -0,1860 to 0,3431 | No | ns | 0.7643 |
| **CD26** | | | | | | | **CD26** | | | | | |
| 4T07 day 7 vs. 4T07 day 14 | 4.44 | -5,279 to 14,15 | No | ns | 0.5302 | | 4T07 day 7 vs. 4T07 day 14 | 0.13 | -0,09785 to 0,3604 | No | ns | 0.3693 |
| 4T07 day 7 vs. 4T07 day 21 | 18.60 | 7,668 to 29,54 | Yes | *** | 0.0002 | | 4T07 day 7 vs. 4T07 day 21 | 0.36 | 0,09721 to 0,6129 | Yes | ** | 0.0037 |
| 4T07 day 14 vs. 4T07 day 21 | 14.17 | 2,946 to 25,38 | Yes | ** | 0.0089 | yes | 4T07 day 14 vs. 4T07 day 21 | 0.22 | -0,04075 to 0,4884 | No | ns | 0.1158 |

Decrease over time

Increase over time

**Supplementary Figure 5** To assess if a problematic batch effect was present in the dataset combined of the three repeats, we made a data normalization within each repeat by using the mean value of each CAF marker in the 4T1 D7 group, and then combined the resultant ratios in a new normalised dataset. We then performed a 2-way ANOVA with Tukey's multiple comparisons post-test like the one done on the raw combined dataset in Fig 3. If there were prominent batch effects within the combined dataset, then statistical comparisons on the ratios would yield different results than the same analyses performed on the raw percentages. Two out of 36 comparisons (5.6%) went from being significant in the raw dataset to not-significant in the normalised dataset, and five comparisons (13.9%) turned out statistically significant in the normalized dataset compared to the raw dataset (Fig S5). In the remaining 86% of the comparisons, nothing or only the level of statistical significance changed. In summary, only a minor batch effect was detected within the combined dataset.

#### Supplementary Tables

**Supplementary Table 1.** Details of 2-way ANOVA and Tukey's multiple comparisons post-test on the Lin- population out of live cells reported in main Fig 2, all repeats combined.

| Table Analyzed | Linneg all repeats | | | | |
|---|---|---|---|---|---|
| Two-way ANOVA | Ordinary | | | | |
| Alpha | 0.05 | | | | |
| Source of Variation | % of total variation | P value | P value summary | Significant? | |
| Interaction | 2.866 | 0.1202 | ns | No | |
| day | 4.131 | 0.0482 | * | Yes | |
| tumour type | 12.2 | < 0,0001 | **** | Yes | |
| ANOVA table | SS | DF | MS | F (DFn, DFd) | P value |
| Interaction | 82.99 | 2 | 41.49 | F (2, 122) = 2,156 | P = 0,1202 |
| day | 119.6 | 2 | 59.81 | F (2, 122) = 3,108 | P = 0,0482 |
| tumour type | 353.4 | 1 | 353.4 | F (1, 122) = 18,36 | P < 0,0001 |
| Residual | 2348 | 122 | 19.25 | | |

| Within each column, compare rows (simple effects within columns) | | | | | | | | |
|---|---|---|---|---|---|---|---|---|
| Number of families | 2 | | | | | | | |
| Number of comparisons per family | 3 | | | | | | | |
| Alpha | 0.05 | | | | | | | |
| Tukey's multiple comparisons test | Mean Diff, | 95% CI of diff, | Significant? | Summary | Adjusted P Value | | | |
| 4T1 | | | | | | | | |
| D7 vs. D14 | 0.7838 | -2,326 to 3,894 | No | ns | 0.8215 | | | |
| D7 vs. D21 | -2.35 | -5,461 to 0,7599 | No | ns | 0.1762 | | | |
| D14 vs. D21 | -3.134 | -6,139 to -0,1294 | Yes | * | 0.0388 | | | |
| 4T07 | | | | | | | | |
| D7 vs. D14 | -2.703 | -5,813 to 0,4071 | No | ns | 0.1021 | | | |
| D7 vs. D21 | -2.561 | -6,061 to 0,9394 | No | ns | 0.1961 | | | |
| D14 vs. D21 | 0.1421 | -3,449 to 3,734 | No | ns | 0.9952 | | | |
| Test details | Mean 1 | Mean 2 | Mean Diff, | SE of diff, | N1 | N2 | q | DF |
| 4T1 | | | | | | | | |
| D7 vs. D14 | 4.95 | 4.17 | 0.78 | 1.31 | 21 | 24 | 0.8456 | 122 |
| D7 vs. D21 | 4.95 | 7.30 | -2.35 | 1.31 | 21 | 24 | 2.536 | 122 |
| D14 vs. D21 | 4.17 | 7.30 | -3.13 | 1.27 | 24 | 24 | 3.5 | 122 |
| 4T07 | | | | | | | | |
| D7 vs. D14 | 7.11 | 9.81 | -2.70 | 1.31 | 24 | 21 | 2.916 | 122 |
| D7 vs. D21 | 7.11 | 9.67 | -2.56 | 1.48 | 24 | 14 | 2.455 | 122 |
| D14 vs. D21 | 9.81 | 9.67 | 0.14 | 1.51 | 21 | 14 | 0.1328 | 122 |

**Supplementary Table 2.** Details of 2-way ANOVA and Tukey's multiple comparisons post-test on the CAF+ population out of Lin- cells reported in main Fig 2, all repeats combined

| | | | | | |
|---|---|---|---|---|---|
| **Table Analyzed** | **CAF+ out of linneg all repeats** | | | | |
| **Two-way ANOVA** | **Ordinary** | | | | |
| **Alpha** | **0.05** | | | | |
| **Source of Variation** | **% of total variation** | **P value** | **P value summary** | **Significant?** | |
| Interaction | 0.32 | 0.8064 | ns | No | |
| day | 10.02 | 0.0015 | ** | Yes | |
| tumour type | 1.45 | 0.1623 | ns | No | |
| **ANOVA table** | **SS** | **DF** | **MS** | **F (DFn, DFd)** | **P value** |
| Interaction | 82.83 | 2 | 41.41 | F (2, 122) = 0,2155 | P = 0,8064 |
| day | 2636 | 2 | 1318 | F (2, 122) = 6,858 | P = 0,0015 |
| tumour type | 379.9 | 1 | 379.9 | F (1, 122) = 1,977 | P = 0,1623 |
| Residual | 23445 | 122 | 192.2 | | |

| | | | | | | | | |
|---|---|---|---|---|---|---|---|---|
| **Within each column, compare rows (simple effects within columns)** | | | | | | | | |
| **Number of families** | **2** | | | | | | | |
| **Number of comparisons per family** | **3** | | | | | | | |
| **Alpha** | **0.05** | | | | | | | |
| **Tukey's multiple comparisons test** | **Mean Diff,** | **95% CI of diff,** | **Significant?** | **Summary** | **Adjusted P Value** | | | |
| 4T1 | | | | | | | | |
| D7 vs. D14 | 7.59 | -2,239 to 17,42 | No | ns | 0.1635 | | | |
| D7 vs. D21 | 8.45 | -1,383 to 18,27 | No | ns | 0.1073 | | | |
| D14 vs. D21 | 0.86 | -8,639 to 10,35 | No | ns | 0.9751 | | | |
| 4T07 | | | | | | | | |
| D7 vs. D14 | 9.69 | -0,1382 to 19,52 | No | ns | 0.0542 | | | |
| D7 vs. D21 | 12.52 | 1,462 to 23,58 | Yes | * | 0.0223 | | | |
| D14 vs. D21 | 2.83 | -8,515 to 14,18 | No | ns | 0.8245 | | | |
| **Test details** | **Mean 1** | **Mean 2** | **Mean Diff,** | **SE of diff,** | **N1** | **N2** | **q** | **DF** |
| 4T1 | | | | | | | | |
| D7 vs. D14 | 73.14 | 65.55 | 7.59 | 4.14 | 21 | 24 | 2.591 | 122 |
| D7 vs. D21 | 73.14 | 64.70 | 8.45 | 4.14 | 21 | 24 | 2.883 | 122 |
| D14 vs. D21 | 65.55 | 64.70 | 0.86 | 4.00 | 24 | 24 | 0.3024 | 122 |
| 4T07 | | | | | | | | |
| D7 vs. D14 | 71.69 | 62.00 | 9.69 | 4.14 | 24 | 21 | 3.308 | 122 |
| D7 vs. D21 | 71.69 | 59.17 | 12.52 | 4.66 | 24 | 14 | 3.799 | 122 |
| D14 vs. D21 | 62.00 | 59.17 | 2.83 | 4.78 | 21 | 14 | 0.8377 | 122 |

**Supplementary Table 3.** Details of 2-way ANOVA and Tukey's multiple comparisons post-test on the CAF+ population out of live cells reported in main Fig 2, all repeats combined

| Table Analyzed | CAF+ out of live cells all repeats | | | | |
|---|---|---|---|---|---|
| Two-way ANOVA | Ordinary | | | | |
| Alpha | 0.05 | | | | |
| Source of Variation | % of total variation | P value | P value summary | Significant? | |
| Interaction | 2.21 | 0.2195 | ns | No | |
| day | 0.24 | 0.8467 | ns | No | |
| tumour type | 9.14 | 0.0005 | *** | Yes | |
| ANOVA table | SS | DF | MS | F (DFn, DFd) | P value |
| Interaction | 24.46 | 2 | 12.23 | F (2, 122) = 1,535 | P = 0,2195 |
| day | 2.655 | 2 | 1.327 | F (2, 122) = 0,1667 | P = 0,8467 |
| tumour type | 101.1 | 1 | 101.1 | F (1, 122) = 12,69 | P = 0,0005 |
| Residual | 971.7 | 122 | 7.965 | | |

| Within each column, compare rows (simple effects within columns) | | | | | | | | |
|---|---|---|---|---|---|---|---|---|
| Number of families | 2 | | | | | | | |
| Number of comparisons per family | 3 | | | | | | | |
| Alpha | 0.05 | | | | | | | |
| Tukey's multiple comparisons test | Mean Diff, | 95% CI of diff, | Significant? | Summary | Adjusted P Value | | | |
| 4T1 | | | | | | | | |
| D7 vs. D14 | 0.94 | -1,058 to 2,944 | No | ns | 0.5046 | | | |
| D7 vs. D21 | -0.37 | -2,372 to 1,630 | No | ns | 0.8989 | | | |
| D14 vs. D21 | -1.31 | -3,247 to 0,6188 | No | ns | 0.244 | | | |
| 4T07 | | | | | | | | |
| D7 vs. D14 | -0.82 | -2,825 to 1,176 | No | ns | 0.5923 | | | |
| D7 vs. D21 | -0.21 | -2,459 to 2,045 | No | ns | 0.9741 | | | |
| D14 vs. D21 | 0.62 | -1,693 to 2,928 | No | ns | 0.8017 | | | |
| Test details | Mean 1 | Mean 2 | Mean Diff, | SE of diff, | N1 | N2 | q | DF |
| 4T1 | | | | | | | | |
| D7 vs. D14 | 3.39 | 2.45 | 0.94 | 0.8433 | 21 | 24 | 1.582 | 122 |
| D7 vs. D21 | 3.39 | 3.76 | -0.37 | 0.8433 | 21 | 24 | 0.6221 | 122 |
| D14 vs. D21 | 2.45 | 3.76 | -1.31 | 0.8147 | 24 | 24 | 2.281 | 122 |
| 4T07 | | | | | | | | |
| D7 vs. D14 | 4.67 | 5.49 | -0.82 | 0.8433 | 24 | 21 | 1.383 | 122 |
| D7 vs. D21 | 4.67 | 4.88 | -0.21 | 0.9491 | 24 | 14 | 0.3086 | 122 |
| D14 vs. D21 | 5.49 | 4.88 | 0.62 | 0.9737 | 21 | 14 | 0.8966 | 122 |

**Supplementary Table 4.** Multiple t-tests pertaining to main Fig 2, all repeats combined. Percentage cells of parent population.

| Multiple t-tests 4T1 vs 4T07 Lin- out of Live cells | | | | | | | | |
|---|---|---|---|---|---|---|---|---|
| Day | Discovery? | P value | Mean1 4T1 | Mean2 4T07 | Difference | SE of difference | t ratio | df |
| D7 | | 0.039520 | 4.95 | 7.11 | -2.15 | 1.01 | 2.12 | 43 |
| D14 | * | 0.000243 | 4.17 | 9.81 | -5.64 | 1.41 | 4.00 | 43 |
| D21 | | 0.168074 | 7.30 | 9.67 | -2.36 | 1.68 | 1.41 | 36 |

| Multiple t-tests 4T1 vs 4T07 CAF+ out of Lin- cells | | | | | | | | |
|---|---|---|---|---|---|---|---|---|
| Day | Discovery? | P value | Mean1 4T1 | Mean2 4T07 | Difference | SE of difference | t ratio | df |
| D7 | | 0.761085 | 73.14 | 71.69 | 1.45 | 4.74 | 0.31 | 43 |
| D14 | | 0.273681 | 65.55 | 62.00 | 3.55 | 3.20 | 1.11 | 43 |
| D21 | | 0.268840 | 64.70 | 59.17 | 5.53 | 4.92 | 1.12 | 36 |

| Multiple t-tests 4T1 vs 4T07 CAF+ out of Live cells | | | | | | | | |
|---|---|---|---|---|---|---|---|---|
| Day | Discovery? | P value | Mean1 4T1 | Mean2 4T07 | Difference | SE of difference | t ratio | df |
| D7 | | 0.116181 | 3.39 | 4.67 | -1.28 | 0.80 | 1.60 | 43 |
| D14 | * | 0.001006 | 2.45 | 5.49 | -3.04 | 0.86 | 3.53 | 43 |
| D21 | | 0.266583 | 3.76 | 4.88 | -1.11 | 0.99 | 1.13 | 36 |

**Supplementary Table 5.** Multiple t-test pertaining to main Fig 3, all repeats combined. Percentage cells positive for each CAF marker within the CAF+ population.

| Multiple t-tests 4T1 vs 4T07 D7 | | | | | | | | |
|---|---|---|---|---|---|---|---|---|
| CAF marker | Discovery? | P value | Mean1 4T1 | Mean2 4T07 | Difference | SE of difference | t ratio | df |
| PDGFRa | | 0.0363678 | 48.99 | 38.28 | 10.71 | 4.96 | 2.16 | 43 |
| PDGFRb | | 0.278426 | 53.38 | 48.40 | 4.98 | 4.54 | 1.10 | 43 |
| Podoplanin | | 0.0041234 | 55.27 | 40.49 | 14.78 | 4.88 | 3.03 | 43 |
| FAP | | 0.0482698 | 46.58 | 51.95 | -5.37 | 2.64 | 2.03 | 43 |
| aSMA | * | 0.00114589 | 29.03 | 19.40 | 9.63 | 2.76 | 3.48 | 43 |
| CD26 | | 0.0419432 | 55.93 | 66.52 | -10.58 | 5.05 | 2.10 | 43 |

| Multiple t-tests 4T1 vs 4T07 D14 | | | | | | | | |
|---|---|---|---|---|---|---|---|---|
| CAF marker | Discovery? | P value | Mean1 4T1 | Mean2 4T07 | Difference | SE of difference | t ratio | df |
| PDGFRa | | 0.147984 | 26.03 | 19.27 | 6.76 | 4.59 | 1.47 | 43 |
| PDGFRb | | 0.0167179 | 31.32 | 43.83 | -12.51 | 5.02 | 2.49 | 43 |
| Podoplanin | | 0.0160842 | 26.11 | 38.00 | -11.89 | 4.75 | 2.51 | 43 |
| FAP | | 0.0290361 | 62.45 | 55.01 | 7.45 | 3.30 | 2.26 | 43 |
| aSMA | * | 0.00013461 | 9.03 | 19.92 | -10.89 | 2.60 | 4.19 | 43 |
| CD26 | | 0.571223 | 62.08 | 64.26 | -2.18 | 3.82 | 0.57 | 43 |

| Multiple t-tests 4T1 vs 4T07 D21 | | | | | | | | |
|---|---|---|---|---|---|---|---|---|
| CAF marker | Discovery? | P value | Mean1 4T1 | Mean2 4T07 | Difference | SE of difference | t ratio | df |
| PDGFRa | * | 0.00138182 | 18.15 | 33.19 | -15.04 | 4.34 | 3.47 | 36 |
| PDGFRb | | 0.132781 | 42.88 | 32.99 | 9.89 | 6.43 | 1.54 | 36 |
| Podoplanin | | 0.0135633 | 31.49 | 20.11 | 11.38 | 4.38 | 2.60 | 36 |
| FAP | | 0.0904156 | 61.98 | 56.44 | 5.54 | 3.19 | 1.74 | 36 |
| aSMA | * | 0.00213947 | 17.93 | 7.59 | 10.34 | 3.12 | 3.31 | 36 |
| CD26 | * | 8.4759E-06 | 65.17 | 47.91 | 17.26 | 3.33 | 5.19 | 36 |

**Supplementary Table 6** Tukey's multiple comparisons from 2-way ANOVA on % of single CAF markers within the CAF+ populations. Data from three independent repeats combined. Significant results reported in main Fig 3.

| | | | | | | | | |
|---|---|---|---|---|---|---|---|---|
| Number of families | 6 | | | | | | | |
| Number of comparisons per family | 3 | | | | Combined | | | |
| Alpha | 0.05 | | | | Repeat 1, 2, 3 | | | |
| Tukey's multiple comparisons test | Mean Diff, | 95% CI of diff, | Significant? | Summary | Adjusted P Value | | | |
| PDGFRa | | | | | | | | |
| 4T1Day 7 vs. 4T1day 14 | 29.72 | 20,23 to 39,22 | Yes | **** | < 0,0001 | | | |
| 4T1Day 7 vs. 4T1day 21 | 30.85 | 21,35 to 40,34 | Yes | **** | < 0,0001 | | | |
| 4T1day 14 vs. 4T1day 21 | 1.124 | -8,047 to 10,29 | No | ns | 0.9552 | | | |
| PDGFRb | | | | | | | | |
| 4T1Day 7 vs. 4T1day 14 | 9.553 | 0,05991 to 19,05 | Yes | * | 0.0482 | | | |
| 4T1Day 7 vs. 4T1day 21 | 10.5 | 1,011 to 20,00 | Yes | * | 0.0259 | | | |
| 4T1day 14 vs. 4T1day 21 | 0.9512 | -8,219 to 10,12 | No | ns | 0.9677 | | | |
| Podoplanin | | | | | | | | |
| 4T1Day 7 vs. 4T1day 14 | 17.27 | 7,772 to 26,76 | Yes | **** | < 0,0001 | | | |
| 4T1Day 7 vs. 4T1day 21 | 23.78 | 14,29 to 33,27 | Yes | **** | < 0,0001 | | | |
| 4T1day 14 vs. 4T1day 21 | 6.515 | -2,656 to 15,69 | No | ns | 0.2175 | | | |
| FAP | | | | | | | | |
| 4T1Day 7 vs. 4T1day 14 | -8.426 | -17,92 to 1,067 | No | ns | 0.0936 | | | |
| 4T1Day 7 vs. 4T1day 21 | -15.4 | -24,89 to -5,905 | Yes | *** | 0.0005 | | | |
| 4T1day 14 vs. 4T1day 21 | -6.972 | -16,14 to 2,199 | No | ns | 0.1748 | | | |
| aSMA | | | | | | | | |
| 4T1Day 7 vs. 4T1day 14 | 9.107 | -0,3859 to 18,60 | No | ns | 0.0632 | | | |
| 4T1Day 7 vs. 4T1day 21 | 11.1 | 1,603 to 20,59 | Yes | * | 0.0171 | | | |
| 4T1day 14 vs. 4T1day 21 | 1.989 | -7,182 to 11,16 | No | ns | 0.8664 | | | |
| CD26 | | | | | | | | |
| 4T1Day 7 vs. 4T1day 14 | -8.326 | -17,82 to 1,166 | No | ns | 0.0989 | | | |
| 4T1Day 7 vs. 4T1day 21 | -9.239 | -18,73 to 0,2539 | No | ns | 0.0584 | | | |
| 4T1day 14 vs. 4T1day 21 | -0.9125 | -10,08 to 8,258 | No | ns | 0.9702 | | | |
| Test details | Mean 1 | Mean 2 | Mean Diff, | SE of diff, | N1 | N2 | q | DF |
| PDGFRa | | | | | | | | |
| 4T1Day 7 vs. 4T1day 14 | 48.99 | 19.27 | 29.72 | 4.035 | 21 | 24 | 10.42 | 396 |
| 4T1Day 7 vs. 4T1day 21 | 48.99 | 18.15 | 30.85 | 4.035 | 21 | 24 | 10.81 | 396 |
| 4T1day 14 vs. 4T1day 21 | 19.27 | 18.15 | 1.124 | 3.898 | 24 | 24 | 0.4077 | 396 |
| PDGFRb | | | | | | | | |
| 4T1Day 7 vs. 4T1day 14 | 53.38 | 43.83 | 9.553 | 4.035 | 21 | 24 | 3.348 | 396 |
| 4T1Day 7 vs. 4T1day 21 | 53.38 | 42.88 | 10.5 | 4.035 | 21 | 24 | 3.681 | 396 |
| 4T1day 14 vs. 4T1day 21 | 43.83 | 42.88 | 0.9512 | 3.898 | 24 | 24 | 0.3451 | 396 |
| Podoplanin | | | | | | | | |
| 4T1Day 7 vs. 4T1day 14 | 55.27 | 38 | 17.27 | 4.035 | 21 | 24 | 6.051 | 396 |
| 4T1Day 7 vs. 4T1day 21 | 55.27 | 31.49 | 23.78 | 4.035 | 21 | 24 | 8.335 | 396 |
| 4T1day 14 vs. 4T1day 21 | 38 | 31.49 | 6.515 | 3.898 | 24 | 24 | 2.364 | 396 |
| FAP | | | | | | | | |
| 4T1Day 7 vs. 4T1day 14 | 46.58 | 55.01 | -8.426 | 4.035 | 21 | 24 | 2.953 | 396 |
| 4T1Day 7 vs. 4T1day 21 | 46.58 | 61.98 | -15.4 | 4.035 | 21 | 24 | 5.397 | 396 |
| 4T1day 14 vs. 4T1day 21 | 55.01 | 61.98 | -6.972 | 3.898 | 24 | 24 | 2.529 | 396 |
| aSMA | | | | | | | | |
| 4T1Day 7 vs. 4T1day 14 | 29.03 | 19.92 | 9.107 | 4.035 | 21 | 24 | 3.192 | 396 |
| 4T1Day 7 vs. 4T1day 21 | 29.03 | 17.93 | 11.1 | 4.035 | 21 | 24 | 3.889 | 396 |
| 4T1day 14 vs. 4T1day 21 | 19.92 | 17.93 | 1.989 | 3.898 | 24 | 24 | 0.7215 | 396 |
| CD26 | | | | | | | | |
| 4T1Day 7 vs. 4T1day 14 | 55.93 | 64.26 | -8.326 | 4.035 | 21 | 24 | 2.918 | 396 |
| 4T1Day 7 vs. 4T1day 21 | 55.93 | 65.17 | -9.239 | 4.035 | 21 | 24 | 3.238 | 396 |
| 4T1day 14 vs. 4T1day 21 | 64.26 | 65.17 | -0.9125 | 3.898 | 24 | 24 | 0.331 | 396 |

| | | | | | | | | |
|---|---|---|---|---|---|---|---|---|
| Number of families | 6 | | | | | | | |
| Number of comparisons per family | 3 | | | | Combined | | | |
| Alpha | 0.05 | | | | Repeat 1, 2, 3 | | | |
| Tukey's multiple comparisons test | Mean Diff, | 95% CI of diff, | Significant? | Summary | Adjusted P Value | | | |
| PDGFRa | | | | | | | | |
| 4T07 day 7 vs. 4T07 day 14 | 12.25 | 2,537 to 21,97 | Yes | ** | 0.009 | | | |
| 4T07 day 7 vs. 4T07 day 21 | 5.095 | -5,840 to 16,03 | No | ns | 0.5167 | | | |
| 4T07 day 14 vs. 4T07 day 21 | -7.158 | -18,38 to 4,061 | No | ns | 0.2912 | | | |
| PDGFRb | | | | | | | | |
| 4T07 day 7 vs. 4T07 day 14 | 17.08 | 7,361 to 26,79 | Yes | *** | 0.0001 | | | |
| 4T07 day 7 vs. 4T07 day 21 | 15.41 | 4,473 to 26,34 | Yes | ** | 0.0029 | | | |
| 4T07 day 14 vs. 4T07 day 21 | -1.669 | -12,89 to 9,550 | No | ns | 0.9346 | | | |
| Podoplanin | | | | | | | | |
| 4T07 day 7 vs. 4T07 day 14 | 14.38 | 4,664 to 24,10 | Yes | ** | 0.0016 | | | |
| 4T07 day 7 vs. 4T07 day 21 | 20.38 | 9,446 to 31,32 | Yes | **** | < 0,0001 | | | |
| 4T07 day 14 vs. 4T07 day 21 | 6.001 | -5,218 to 17,22 | No | ns | 0.4194 | | | |
| FAP | | | | | | | | |
| 4T07 day 7 vs. 4T07 day 14 | -10.5 | -20,22 to -0,7858 | Yes | * | 0.0305 | | | |
| 4T07 day 7 vs. 4T07 day 21 | -4.484 | -15,42 to 6,452 | No | ns | 0.5994 | | | |
| 4T07 day 14 vs. 4T07 day 21 | 6.018 | -5,201 to 17,24 | No | ns | 0.4173 | | | |
| aSMA | | | | | | | | |
| 4T07 day 7 vs. 4T07 day 14 | 10.36 | 0,6476 to 20,08 | Yes | * | 0.0334 | | | |
| 4T07 day 7 vs. 4T07 day 21 | 11.8 | 0,8686 to 22,74 | Yes | * | 0.0308 | | | |
| 4T07 day 14 vs. 4T07 day 21 | 1.44 | -9,779 to 12,66 | No | ns | 0.9509 | | | |
| CD26 | | | | | | | | |
| 4T07 day 7 vs. 4T07 day 14 | 4.437 | -5,279 to 14,15 | No | ns | 0.5302 | | | |
| 4T07 day 7 vs. 4T07 day 21 | 18.6 | 7,668 to 29,54 | Yes | *** | 0.0002 | | | |
| 4T07 day 14 vs. 4T07 day 21 | 14.17 | 2,946 to 25,38 | Yes | ** | 0.0089 | | | |
| Test details | Mean 1 | Mean 2 | Mean Diff, | SE of diff, | N1 | N2 | q | DF |
| PDGFRa | | | | | | | | |
| 4T07 day 7 vs. 4T07 day 14 | 38.28 | 26.03 | 12.25 | 4.127 | 24 | 21 | 4.199 | 336 |
| 4T07 day 7 vs. 4T07 day 21 | 38.28 | 33.19 | 5.095 | 4.645 | 24 | 14 | 1.551 | 336 |
| 4T07 day 14 vs. 4T07 day 21 | 26.03 | 33.19 | -7.158 | 4.766 | 21 | 14 | 2.124 | 336 |
| PDGFRb | | | | | | | | |
| 4T07 day 7 vs. 4T07 day 14 | 48.4 | 31.32 | 17.08 | 4.127 | 24 | 21 | 5.852 | 336 |
| 4T07 day 7 vs. 4T07 day 21 | 48.4 | 32.99 | 15.41 | 4.645 | 24 | 14 | 4.691 | 336 |
| 4T07 day 14 vs. 4T07 day 21 | 31.32 | 32.99 | -1.669 | 4.766 | 21 | 14 | 0.4953 | 336 |
| Podoplanin | | | | | | | | |
| 4T07 day 7 vs. 4T07 day 14 | 40.49 | 26.11 | 14.38 | 4.127 | 24 | 21 | 4.928 | 336 |
| 4T07 day 7 vs. 4T07 day 21 | 40.49 | 20.11 | 20.38 | 4.645 | 24 | 14 | 6.205 | 336 |
| 4T07 day 14 vs. 4T07 day 21 | 26.11 | 20.11 | 6.001 | 4.766 | 21 | 14 | 1.781 | 336 |
| FAP | | | | | | | | |
| 4T07 day 7 vs. 4T07 day 14 | 51.95 | 62.45 | -10.5 | 4.127 | 24 | 21 | 3.599 | 336 |
| 4T07 day 7 vs. 4T07 day 21 | 51.95 | 56.44 | -4.484 | 4.645 | 24 | 14 | 1.365 | 336 |
| 4T07 day 14 vs. 4T07 day 21 | 62.45 | 56.44 | 6.018 | 4.766 | 21 | 14 | 1.786 | 336 |
| aSMA | | | | | | | | |
| 4T07 day 7 vs. 4T07 day 14 | 19.4 | 9.034 | 10.36 | 4.127 | 24 | 21 | 3.551 | 336 |
| 4T07 day 7 vs. 4T07 day 21 | 19.4 | 7.594 | 11.8 | 4.645 | 24 | 14 | 3.594 | 336 |
| 4T07 day 14 vs. 4T07 day 21 | 9.034 | 7.594 | 1.44 | 4.766 | 21 | 14 | 0.4273 | 336 |
| CD26 | | | | | | | | |
| 4T07 day 7 vs. 4T07 day 14 | 66.52 | 62.08 | 4.437 | 4.127 | 24 | 21 | 1.521 | 336 |
| 4T07 day 7 vs. 4T07 day 21 | 66.52 | 47.91 | 18.6 | 4.645 | 24 | 14 | 5.664 | 336 |
| 4T07 day 14 vs. 4T07 day 21 | 62.08 | 47.91 | 14.17 | 4.766 | 21 | 14 | 4.204 | 336 |

**Supplementary Table 7** Co-expression tables of two CAF markers in 4T1 and 4T07 tumours at different time points, expanding the analysis to contain the time dimension. First CAF marker = parent gate, Subset CAF marker = child gate. CI95 = 95% confidence interval.

## 4T1

| Day 7 | ① First CAF marker: PDGFRa | | | PDGFRb | | | Podoplanin | | | FAPa | | | aSMA | | | CD26 | | |
|---|---|---|---|---|---|---|---|---|---|---|---|---|---|---|---|---|---|---|
| ② Subset CAF marker | Mean | CI95 low | CI95 high | Mean | CI95 low | CI95 high | Mean | CI95 low | CI95 high | Mean | CI95 low | CI95 high | Mean | CI95 low | CI95 high | Mean | CI95 low | CI95 high |
| PDGFRa | 100.00 | 100.00 | 100.00 | 50.67 | 43.01 | 58.33 | 65.14 | 59.10 | 71.18 | 38.09 | 29.13 | 47.05 | 49.81 | 42.43 | 57.19 | 45.86 | 36.29 | 55.43 |
| PDGFRb | 58.00 | 50.04 | 65.96 | 100.00 | 100.00 | 100.00 | 68.52 | 61.14 | 75.90 | 62.02 | 57.46 | 66.57 | 71.93 | 64.62 | 79.24 | 60.66 | 53.67 | 67.65 |
| Podoplanin | 77.70 | 68.49 | 86.91 | 68.42 | 63.67 | 73.16 | 100.00 | 100.00 | 100.00 | 41.52 | 32.15 | 50.88 | 88.92 | 84.20 | 93.64 | 64.04 | 55.37 | 72.70 |
| FAPa | 34.55 | 29.23 | 39.87 | 55.69 | 49.30 | 62.07 | 34.26 | 28.46 | 40.05 | 100.00 | 100.00 | 100.00 | 26.32 | 19.64 | 32.99 | 49.32 | 46.41 | 52.23 |
| aSMA | 31.62 | 25.84 | 37.40 | 37.84 | 33.57 | 42.11 | 47.37 | 41.18 | 53.56 | 16.07 | 11.27 | 20.86 | 100.00 | 100.00 | 100.00 | 33.11 | 26.95 | 39.27 |
| CD26 | 52.88 | 42.70 | 63.05 | 62.00 | 55.79 | 68.22 | 63.40 | 55.97 | 70.82 | 59.04 | 51.98 | 66.10 | 60.70 | 53.69 | 67.70 | 100.00 | 100.00 | 100.00 |

| Day 14 | ① First CAF marker: PDGFRa | | | PDGFRb | | | Podoplanin | | | FAPa | | | aSMA | | | CD26 | | |
|---|---|---|---|---|---|---|---|---|---|---|---|---|---|---|---|---|---|---|
| ② Subset CAF marker | Mean | CI95 low | CI95 high | Mean | CI95 low | CI95 high | Mean | CI95 low | CI95 high | Mean | CI95 low | CI95 high | Mean | CI95 low | CI95 high | Mean | CI95 low | CI95 high |
| PDGFRa | 100.00 | 100.00 | 100.00 | 26.67 | 19.45 | 33.89 | 40.34 | 33.42 | 47.26 | 15.06 | 10.63 | 19.50 | 25.95 | 20.65 | 31.24 | 22.00 | 14.83 | 29.18 |
| PDGFRb | 62.74 | 56.76 | 68.71 | 100.00 | 100.00 | 100.00 | 66.43 | 60.65 | 72.20 | 52.21 | 45.06 | 59.35 | 70.22 | 65.81 | 74.62 | 46.62 | 38.56 | 54.67 |
| Podoplanin | 84.31 | 79.56 | 89.06 | 55.06 | 47.71 | 62.41 | 100.00 | 100.00 | 100.00 | 27.29 | 21.02 | 33.56 | 88.67 | 86.59 | 90.74 | 43.91 | 34.39 | 53.44 |
| FAPa | 46.95 | 39.76 | 54.14 | 67.37 | 62.77 | 71.98 | 43.09 | 35.59 | 50.60 | 100.00 | 100.00 | 100.00 | 31.62 | 24.98 | 38.26 | 49.72 | 45.16 | 54.28 |
| aSMA | 27.58 | 23.21 | 31.95 | 29.62 | 25.50 | 33.74 | 46.06 | 39.87 | 52.25 | 10.12 | 7.19 | 13.04 | 100.00 | 100.00 | 100.00 | 19.87 | 15.27 | 24.47 |
| CD26 | 71.69 | 66.72 | 76.65 | 67.91 | 63.38 | 72.44 | 71.83 | 66.91 | 76.76 | 57.34 | 54.43 | 60.25 | 62.95 | 58.42 | 67.49 | 100.00 | 100.00 | 100.00 |

| Day 21 | ① First CAF marker: PDGFRa | | | PDGFRb | | | Podoplanin | | | FAPa | | | aSMA | | | CD26 | | |
|---|---|---|---|---|---|---|---|---|---|---|---|---|---|---|---|---|---|---|
| ② Subset CAF marker | Mean | CI95 low | CI95 high | Mean | CI95 low | CI95 high | Mean | CI95 low | CI95 high | Mean | CI95 low | CI95 high | Mean | CI95 low | CI95 high | Mean | CI95 low | CI95 high |
| PDGFRa | 100.00 | 100.00 | 100.00 | 26.74 | 20.16 | 33.31 | 42.85 | 38.55 | 47.15 | 13.88 | 10.64 | 17.13 | 26.07 | 22.21 | 29.94 | 20.66 | 15.69 | 25.63 |
| PDGFRb | 56.32 | 48.39 | 64.25 | 100.00 | 100.00 | 100.00 | 66.96 | 60.11 | 73.80 | 47.74 | 39.82 | 55.67 | 71.59 | 64.28 | 78.91 | 46.83 | 38.40 | 55.26 |
| Podoplanin | 74.90 | 65.76 | 84.03 | 51.43 | 42.87 | 59.99 | 100.00 | 100.00 | 100.00 | 23.23 | 17.89 | 28.58 | 87.63 | 84.68 | 90.58 | 36.34 | 29.21 | 43.47 |
| FAPa | 47.46 | 41.19 | 53.73 | 70.59 | 65.03 | 76.14 | 47.77 | 39.65 | 55.89 | 100.00 | 100.00 | 100.00 | 36.46 | 29.18 | 43.74 | 57.52 | 54.11 | 60.92 |
| aSMA | 25.58 | 19.26 | 31.90 | 29.82 | 22.81 | 36.84 | 45.66 | 40.75 | 50.58 | 10.53 | 6.54 | 14.52 | 100.00 | 100.00 | 100.00 | 18.16 | 13.45 | 22.87 |
| CD26 | 70.56 | 63.85 | 77.28 | 71.27 | 67.12 | 75.43 | 73.88 | 71.09 | 76.66 | 60.35 | 57.60 | 63.11 | 66.14 | 62.42 | 69.86 | 100.00 | 100.00 | 100.00 |

## 4T07

| Day 7 | ① First CAF marker: PDGFRa | | | PDGFRb | | | Podoplanin | | | FAPa | | | aSMA | | | CD26 | | |
|---|---|---|---|---|---|---|---|---|---|---|---|---|---|---|---|---|---|---|
| ② Subset CAF marker | Mean | CI95 low | CI95 high | Mean | CI95 low | CI95 high | Mean | CI95 low | CI95 high | Mean | CI95 low | CI95 high | Mean | CI95 low | CI95 high | Mean | CI95 low | CI95 high |
| PDGFRa | 100.00 | 100.00 | 100.00 | 46.06 | 39.96 | 52.15 | 65.84 | 61.33 | 70.35 | 29.68 | 24.32 | 35.04 | 51.11 | 45.54 | 56.68 | 35.77 | 29.76 | 41.77 |
| PDGFRb | 60.93 | 53.01 | 68.86 | 100.00 | 100.00 | 100.00 | 73.94 | 69.47 | 78.40 | 57.16 | 52.16 | 62.15 | 80.34 | 76.63 | 84.05 | 54.01 | 47.07 | 60.95 |
| Podoplanin | 75.81 | 65.84 | 85.79 | 60.42 | 55.34 | 65.51 | 100.00 | 100.00 | 100.00 | 34.01 | 29.30 | 38.71 | 88.01 | 83.96 | 92.06 | 48.71 | 41.78 | 55.65 |
| FAPa | 41.09 | 36.47 | 45.72 | 63.80 | 58.82 | 68.78 | 46.81 | 40.94 | 52.68 | 100.00 | 100.00 | 100.00 | 38.05 | 31.57 | 44.54 | 51.10 | 48.16 | 54.05 |
| aSMA | 29.10 | 23.46 | 34.75 | 31.99 | 27.61 | 36.38 | 42.25 | 37.88 | 46.62 | 13.47 | 10.43 | 16.51 | 100.00 | 100.00 | 100.00 | 22.20 | 18.48 | 25.93 |
| CD26 | 66.62 | 55.90 | 77.33 | 72.07 | 67.52 | 76.61 | 78.21 | 73.32 | 83.11 | 63.88 | 58.34 | 69.42 | 74.91 | 69.13 | 80.70 | 100.00 | 100.00 | 100.00 |

| Day 14 | ① First CAF marker: PDGFRa | | | PDGFRb | | | Podoplanin | | | FAPa | | | aSMA | | | CD26 | | |
|---|---|---|---|---|---|---|---|---|---|---|---|---|---|---|---|---|---|---|
| ② Subset CAF marker | Mean | CI95 low | CI95 high | Mean | CI95 low | CI95 high | Mean | CI95 low | CI95 high | Mean | CI95 low | CI95 high | Mean | CI95 low | CI95 high | Mean | CI95 low | CI95 high |
| PDGFRa | 100.00 | 100.00 | 100.00 | 35.67 | 28.90 | 42.44 | 58.62 | 52.58 | 64.65 | 19.44 | 13.47 | 25.41 | 50.10 | 44.13 | 56.07 | 23.38 | 17.04 | 29.71 |
| PDGFRb | 43.33 | 36.41 | 50.25 | 100.00 | 100.00 | 100.00 | 54.13 | 47.46 | 60.79 | 38.53 | 31.06 | 46.00 | 66.19 | 61.04 | 71.34 | 36.84 | 28.09 | 45.58 |
| Podoplanin | 66.72 | 56.53 | 76.90 | 46.78 | 39.87 | 53.69 | 100.00 | 100.00 | 100.00 | 22.76 | 17.44 | 28.08 | 88.07 | 83.57 | 92.58 | 31.27 | 24.02 | 38.52 |
| FAPa | 45.55 | 39.42 | 51.68 | 78.53 | 74.81 | 82.25 | 54.76 | 49.91 | 59.61 | 100.00 | 100.00 | 100.00 | 50.95 | 44.35 | 57.54 | 57.50 | 51.22 | 63.78 |
| aSMA | 19.89 | 15.78 | 24.00 | 19.99 | 16.28 | 23.69 | 30.66 | 25.92 | 35.40 | 7.11 | 5.24 | 8.98 | 100.00 | 100.00 | 100.00 | 9.73 | 7.29 | 12.18 |
| CD26 | 62.39 | 50.86 | 73.91 | 71.55 | 66.32 | 76.78 | 73.89 | 66.94 | 80.83 | 55.90 | 50.75 | 61.05 | 66.29 | 59.84 | 72.74 | 100.00 | 100.00 | 100.00 |

| Day 21 | ① First CAF marker: PDGFRa | | | PDGFRb | | | Podoplanin | | | FAPa | | | aSMA | | | CD26 | | |
|---|---|---|---|---|---|---|---|---|---|---|---|---|---|---|---|---|---|---|
| ② Subset CAF marker | Mean | CI95 low | CI95 high | Mean | CI95 low | CI95 high | Mean | CI95 low | CI95 high | Mean | CI95 low | CI95 high | Mean | CI95 low | CI95 high | Mean | CI95 low | CI95 high |
| PDGFRa | 100.00 | 100.00 | 100.00 | 34.42 | 24.70 | 44.14 | 59.55 | 51.54 | 67.57 | 20.27 | 13.43 | 27.10 | 47.31 | 41.62 | 53.00 | 25.21 | 16.60 | 33.82 |
| PDGFRb | 38.98 | 25.05 | 52.91 | 100.00 | 100.00 | 100.00 | 58.09 | 47.43 | 68.74 | 40.88 | 30.98 | 50.77 | 65.45 | 52.47 | 78.43 | 37.64 | 26.88 | 48.40 |
| Podoplanin | 45.10 | 29.38 | 60.82 | 38.22 | 28.35 | 48.09 | 100.00 | 100.00 | 100.00 | 15.72 | 11.82 | 19.63 | 82.57 | 71.31 | 93.84 | 24.73 | 17.50 | 31.96 |
| FAPa | 34.45 | 28.43 | 40.47 | 75.58 | 68.22 | 82.94 | 48.50 | 40.28 | 56.71 | 100.00 | 100.00 | 100.00 | 35.98 | 27.41 | 44.55 | 56.77 | 50.52 | 63.02 |
| aSMA | 14.22 | 7.99 | 20.45 | 15.49 | 10.16 | 20.82 | 29.70 | 21.98 | 37.42 | 4.16 | 2.67 | 5.65 | 100.00 | 100.00 | 100.00 | 6.84 | 4.53 | 9.15 |
| CD26 | 42.43 | 29.01 | 55.85 | 56.74 | 49.29 | 64.18 | 59.70 | 51.50 | 67.90 | 47.12 | 43.03 | 51.21 | 46.53 | 37.30 | 55.76 | 100.00 | 100.00 | 100.00 |

**Supplementary Table 8. Abundance ranking of Boolean CAF subpopulations in 4T1 tumours.** Three independent repeats combined, mean % of CAF+ population. Top 5 subpopulations also presented in main Fig 5.

| | 4T1 D7 | | | | 4T1 D14 | | | | 4T1 D21 | | | |
|---|---|---|---|---|---|---|---|---|---|---|---|---|
| Top | CAF subset | Mean % | SEM | n | CAF subset | Mean % | SEM | n | CAF subset | Mean % | SEM | n |
| 1 | S56 | 8.07 | 1.17 | 21 | S32 | 16.45 | 3.13 | 24 | S32 | 15.09 | 1.96 | 24 |
| 2 | S63 | 7.07 | 2.14 | 21 | S56 | 13.98 | 1.34 | 24 | S56 | 14.55 | 0.76 | 24 |
| 3 | S22 | 6.87 | 0.85 | 21 | S22 | 10.38 | 1.31 | 24 | S22 | 13.21 | 2.03 | 24 |
| 4 | S32 | 5.84 | 1.33 | 21 | S24 | 8.11 | 0.96 | 24 | S24 | 11.35 | 1.23 | 24 |
| 5 | S17 | 5.82 | 0.88 | 21 | S54 | 6.41 | 0.95 | 24 | S54 | 6.19 | 1.25 | 24 |
| 6 | S27 | 4.83 | 1.20 | 21 | S10 | 3.84 | 0.60 | 24 | S10 | 2.97 | 0.52 | 24 |
| 7 | S54 | 4.45 | 0.48 | 21 | S27 | 3.78 | 1.06 | 24 | S17 | 2.72 | 0.32 | 24 |
| 8 | S9 | 4.20 | 0.63 | 21 | S17 | 3.39 | 0.51 | 24 | S18 | 2.09 | 0.28 | 24 |
| 9 | S10 | 3.78 | 0.59 | 21 | S42 | 2.72 | 0.57 | 24 | S25 | 2.02 | 0.38 | 24 |
| 10 | S41 | 3.56 | 0.71 | 21 | S18 | 2.49 | 0.18 | 24 | S2 | 1.97 | 0.21 | 24 |
| 11 | S1 | 3.55 | 0.75 | 21 | S12 | 2.07 | 0.32 | 24 | S27 | 1.92 | 0.43 | 24 |
| 12 | S24 | 3.24 | 0.61 | 21 | S2 | 1.94 | 0.24 | 24 | S42 | 1.79 | 0.33 | 24 |
| 13 | S57 | 3.23 | 0.44 | 21 | S25 | 1.92 | 0.34 | 24 | S1 | 1.66 | 0.35 | 24 |
| 14 | S25 | 2.59 | 0.31 | 21 | S1 | 1.66 | 0.29 | 24 | S9 | 1.50 | 0.24 | 24 |
| 15 | S42 | 2.44 | 0.44 | 21 | S62 | 1.66 | 0.25 | 24 | S62 | 1.47 | 0.14 | 24 |
| 16 | S59 | 2.15 | 0.31 | 21 | S44 | 1.47 | 0.36 | 24 | S63 | 1.45 | 0.34 | 24 |
| 17 | S62 | 2.06 | 0.30 | 21 | S28 | 1.46 | 0.31 | 24 | S12 | 1.35 | 0.23 | 24 |
| 18 | S19 | 2.02 | 0.53 | 21 | S63 | 1.40 | 0.65 | 24 | S20 | 1.17 | 0.17 | 24 |
| 19 | S12 | 1.95 | 0.37 | 21 | S9 | 1.36 | 0.17 | 24 | S44 | 1.13 | 0.25 | 24 |
| 20 | S11 | 1.92 | 0.45 | 21 | S19 | 1.14 | 0.28 | 24 | S19 | 1.10 | 0.21 | 24 |
| 21 | S18 | 1.74 | 0.17 | 21 | S20 | 1.05 | 0.17 | 24 | S48 | 1.06 | 0.27 | 24 |
| 22 | S2 | 1.40 | 0.18 | 21 | S48 | 0.90 | 0.14 | 24 | S55 | 0.82 | 0.23 | 24 |
| 23 | S49 | 1.28 | 0.12 | 21 | S26 | 0.85 | 0.12 | 24 | S60 | 0.69 | 0.10 | 24 |
| 24 | S55 | 1.27 | 0.44 | 21 | S41 | 0.74 | 0.15 | 24 | S28 | 0.64 | 0.08 | 24 |
| 25 | S48 | 1.16 | 0.22 | 21 | S50 | 0.70 | 0.06 | 24 | S26 | 0.64 | 0.11 | 24 |
| 26 | S21 | 1.04 | 0.18 | 21 | S58 | 0.67 | 0.11 | 24 | S11 | 0.59 | 0.12 | 24 |
| 27 | S28 | 1.00 | 0.21 | 21 | S57 | 0.65 | 0.11 | 24 | S23 | 0.57 | 0.08 | 24 |
| 28 | S58 | 0.95 | 0.17 | 21 | S60 | 0.62 | 0.05 | 24 | S52 | 0.55 | 0.08 | 24 |
| 29 | S44 | 0.81 | 0.09 | 21 | S11 | 0.58 | 0.16 | 24 | S41 | 0.53 | 0.08 | 24 |
| 30 | S26 | 0.71 | 0.09 | 21 | S52 | 0.53 | 0.08 | 24 | S50 | 0.53 | 0.05 | 24 |
| 31 | S20 | 0.70 | 0.12 | 21 | S34 | 0.48 | 0.05 | 24 | S57 | 0.52 | 0.08 | 24 |
| 32 | S43 | 0.67 | 0.08 | 21 | S46 | 0.45 | 0.07 | 24 | S58 | 0.50 | 0.09 | 24 |
| 33 | S50 | 0.61 | 0.09 | 21 | S59 | 0.44 | 0.07 | 24 | S4 | 0.46 | 0.14 | 24 |
| 34 | S3 | 0.59 | 0.20 | 21 | S55 | 0.44 | 0.12 | 24 | S59 | 0.46 | 0.10 | 24 |
| 35 | S60 | 0.58 | 0.06 | 21 | S30 | 0.38 | 0.05 | 24 | S34 | 0.45 | 0.06 | 24 |
| 36 | S53 | 0.57 | 0.10 | 21 | S4 | 0.32 | 0.08 | 24 | S30 | 0.43 | 0.04 | 24 |
| 37 | S33 | 0.51 | 0.06 | 21 | S49 | 0.31 | 0.03 | 24 | S21 | 0.38 | 0.06 | 24 |
| 38 | S46 | 0.51 | 0.09 | 21 | S21 | 0.26 | 0.03 | 24 | S31 | 0.37 | 0.18 | 24 |
| 39 | S31 | 0.50 | 0.14 | 21 | S40 | 0.23 | 0.05 | 24 | S40 | 0.31 | 0.09 | 24 |
| 40 | S23 | 0.44 | 0.08 | 21 | S36 | 0.19 | 0.05 | 24 | S49 | 0.31 | 0.03 | 24 |
| 41 | S4 | 0.40 | 0.13 | 21 | S43 | 0.18 | 0.04 | 24 | S3 | 0.29 | 0.09 | 24 |
| 42 | S51 | 0.39 | 0.05 | 21 | S23 | 0.17 | 0.03 | 24 | S36 | 0.28 | 0.09 | 24 |
| 43 | S52 | 0.37 | 0.09 | 21 | S3 | 0.16 | 0.05 | 24 | S43 | 0.25 | 0.07 | 24 |
| 44 | S34 | 0.35 | 0.05 | 21 | S31 | 0.16 | 0.06 | 24 | S46 | 0.24 | 0.03 | 24 |
| 45 | S61 | 0.34 | 0.07 | 21 | S33 | 0.16 | 0.03 | 24 | S33 | 0.22 | 0.06 | 24 |
| 46 | S30 | 0.19 | 0.03 | 21 | S51 | 0.10 | 0.02 | 24 | S16 | 0.21 | 0.06 | 24 |
| 47 | S6 | 0.18 | 0.04 | 21 | S8 | 0.09 | 0.03 | 24 | S8 | 0.21 | 0.05 | 24 |
| 48 | S38 | 0.16 | 0.04 | 21 | S53 | 0.09 | 0.01 | 24 | S6 | 0.15 | 0.02 | 24 |
| 49 | S40 | 0.16 | 0.04 | 21 | S16 | 0.09 | 0.02 | 24 | S51 | 0.13 | 0.03 | 24 |
| 50 | S36 | 0.11 | 0.02 | 21 | S38 | 0.08 | 0.01 | 24 | S38 | 0.11 | 0.02 | 24 |
| 51 | S35 | 0.10 | 0.02 | 21 | S6 | 0.08 | 0.01 | 24 | S53 | 0.10 | 0.02 | 24 |
| 52 | S8 | 0.10 | 0.03 | 21 | S61 | 0.08 | 0.03 | 24 | S61 | 0.07 | 0.02 | 24 |
| 53 | S29 | 0.08 | 0.02 | 21 | S14 | 0.05 | 0.01 | 24 | S35 | 0.05 | 0.02 | 24 |
| 54 | S14 | 0.08 | 0.02 | 21 | S35 | 0.03 | 0.01 | 24 | S29 | 0.04 | 0.01 | 24 |
| 55 | S16 | 0.07 | 0.01 | 21 | S29 | 0.02 | 0.00 | 24 | S14 | 0.03 | 0.01 | 24 |
| 56 | S45 | 0.07 | 0.02 | 21 | S45 | 0.01 | 0.00 | 24 | S47 | 0.03 | 0.01 | 24 |
| 57 | S5 | 0.07 | 0.02 | 21 | S5 | 0.01 | 0.00 | 24 | S5 | 0.02 | 0.01 | 24 |
| 58 | S47 | 0.06 | 0.01 | 21 | S47 | 0.01 | 0.00 | 24 | S37 | 0.01 | 0.00 | 24 |
| 59 | S7 | 0.03 | 0.01 | 21 | S37 | 0.00 | 0.00 | 24 | S39 | 0.01 | 0.00 | 24 |
| 60 | S15 | 0.03 | 0.01 | 21 | S39 | 0.00 | 0.00 | 24 | S7 | 0.01 | 0.00 | 24 |
| 61 | S37 | 0.02 | 0.01 | 21 | S7 | 0.00 | 0.00 | 24 | S45 | 0.01 | 0.00 | 24 |
| 62 | S39 | 0.02 | 0.01 | 21 | S15 | 0.00 | 0.00 | 24 | S15 | 0.01 | 0.00 | 24 |
| 63 | S13 | 0.01 | 0.00 | 21 | S13 | 0.00 | 0.00 | 24 | S13 | 0.00 | 0.00 | 24 |

**Supplementary Table 9. Abundance ranking of Boolean CAF subpopulations in 4T07 tumours.** Three independent repeats combined, mean % of CAF+ population. Top 5 subpopulations also presented in main Fig 5.

| | 4T07 D7 | | | | 4T07 D14 | | | | 4T07 D21 | | | |
|---|---|---|---|---|---|---|---|---|---|---|---|---|
| Top | CAF subset | Mean % | SEM | n | CAF subset | Mean % | SEM | n | CAF subset | Mean % | SEM | n |
| 1 | S32 | 14.44 | 1.94 | 24 | S56 | 17.73 | 1.08 | 21 | S56 | 16.88 | 1.21 | 14 |
| 2 | S56 | 9.48 | 0.82 | 24 | S32 | 17.32 | 2.50 | 21 | S32 | 13.72 | 2.49 | 14 |
| 3 | S22 | 9.39 | 1.05 | 24 | S24 | 12.32 | 1.61 | 21 | S63 | 13.69 | 4.41 | 14 |
| 4 | S24 | 7.23 | 0.94 | 24 | S22 | 9.88 | 1.43 | 21 | S24 | 8.95 | 1.49 | 14 |
| 5 | S63 | 6.49 | 2.62 | 24 | S63 | 6.03 | 3.02 | 21 | S22 | 8.91 | 1.89 | 14 |
| 6 | S17 | 5.33 | 0.48 | 24 | S54 | 4.70 | 1.02 | 21 | S54 | 7.66 | 1.82 | 14 |
| 7 | S54 | 4.52 | 0.52 | 24 | S27 | 3.56 | 0.84 | 21 | S27 | 2.45 | 0.72 | 14 |
| 8 | S9 | 4.48 | 0.60 | 24 | S17 | 3.29 | 0.56 | 21 | S55 | 2.31 | 0.70 | 14 |
| 9 | S25 | 3.55 | 0.52 | 24 | S19 | 1.94 | 0.42 | 21 | S17 | 2.10 | 0.42 | 14 |
| 10 | S1 | 3.15 | 0.59 | 24 | S18 | 1.88 | 0.24 | 21 | S62 | 1.72 | 0.40 | 14 |
| 11 | S10 | 2.97 | 0.57 | 24 | S1 | 1.59 | 0.30 | 21 | S21 | 1.17 | 0.24 | 14 |
| 12 | S27 | 2.96 | 0.52 | 24 | S55 | 1.52 | 0.48 | 21 | S25 | 1.13 | 0.34 | 14 |
| 13 | S18 | 2.07 | 0.21 | 24 | S20 | 1.33 | 0.19 | 21 | S19 | 1.12 | 0.29 | 14 |
| 14 | S2 | 1.78 | 0.16 | 24 | S25 | 1.16 | 0.18 | 21 | S31 | 1.06 | 0.41 | 14 |
| 15 | S57 | 1.70 | 0.31 | 24 | S2 | 1.09 | 0.15 | 21 | S23 | 1.02 | 0.20 | 14 |
| 16 | S19 | 1.52 | 0.22 | 24 | S28 | 0.97 | 0.22 | 21 | S42 | 0.99 | 0.37 | 14 |
| 17 | S62 | 1.36 | 0.12 | 24 | S62 | 0.89 | 0.16 | 21 | S18 | 0.91 | 0.11 | 14 |
| 18 | S55 | 1.18 | 0.46 | 24 | S9 | 0.88 | 0.17 | 21 | S41 | 0.84 | 0.20 | 14 |
| 19 | S41 | 1.04 | 0.20 | 24 | S52 | 0.71 | 0.10 | 21 | S61 | 0.80 | 0.18 | 14 |
| 20 | S59 | 1.02 | 0.17 | 24 | S21 | 0.69 | 0.19 | 21 | S1 | 0.76 | 0.17 | 14 |
| 21 | S11 | 0.94 | 0.19 | 24 | S11 | 0.61 | 0.14 | 21 | S59 | 0.71 | 0.17 | 14 |
| 22 | S28 | 0.88 | 0.13 | 24 | S60 | 0.57 | 0.10 | 21 | S9 | 0.71 | 0.24 | 14 |
| 23 | S26 | 0.87 | 0.19 | 24 | S10 | 0.56 | 0.13 | 21 | S50 | 0.68 | 0.13 | 14 |
| 24 | S49 | 0.77 | 0.09 | 24 | S41 | 0.55 | 0.11 | 21 | S20 | 0.66 | 0.10 | 14 |
| 25 | S21 | 0.76 | 0.09 | 24 | S42 | 0.55 | 0.16 | 21 | S57 | 0.63 | 0.12 | 14 |
| 26 | S42 | 0.74 | 0.12 | 24 | S50 | 0.54 | 0.12 | 21 | S44 | 0.62 | 0.20 | 14 |
| 27 | S20 | 0.72 | 0.09 | 24 | S23 | 0.53 | 0.08 | 21 | S60 | 0.62 | 0.14 | 14 |
| 28 | S12 | 0.69 | 0.11 | 24 | S31 | 0.48 | 0.14 | 21 | S48 | 0.59 | 0.27 | 14 |
| 29 | S48 | 0.66 | 0.12 | 24 | S3 | 0.45 | 0.12 | 21 | S49 | 0.58 | 0.10 | 14 |
| 30 | S23 | 0.62 | 0.08 | 24 | S57 | 0.44 | 0.08 | 21 | S58 | 0.58 | 0.20 | 14 |
| 31 | S53 | 0.52 | 0.17 | 24 | S59 | 0.43 | 0.07 | 21 | S28 | 0.57 | 0.17 | 14 |
| 32 | S61 | 0.51 | 0.11 | 24 | S12 | 0.36 | 0.06 | 21 | S53 | 0.56 | 0.11 | 14 |
| 33 | S58 | 0.44 | 0.10 | 24 | S61 | 0.36 | 0.09 | 21 | S10 | 0.53 | 0.21 | 14 |
| 34 | S30 | 0.43 | 0.07 | 24 | S44 | 0.35 | 0.05 | 21 | S2 | 0.39 | 0.08 | 14 |
| 35 | S31 | 0.43 | 0.09 | 24 | S49 | 0.35 | 0.06 | 21 | S11 | 0.33 | 0.10 | 14 |
| 36 | S50 | 0.41 | 0.06 | 24 | S34 | 0.31 | 0.05 | 21 | S52 | 0.32 | 0.06 | 14 |
| 37 | S6 | 0.40 | 0.15 | 24 | S48 | 0.29 | 0.07 | 21 | S51 | 0.31 | 0.11 | 14 |
| 38 | S60 | 0.35 | 0.07 | 24 | S51 | 0.28 | 0.08 | 21 | S26 | 0.29 | 0.10 | 14 |
| 39 | S51 | 0.31 | 0.07 | 24 | S30 | 0.27 | 0.06 | 21 | S12 | 0.25 | 0.07 | 14 |
| 40 | S3 | 0.27 | 0.06 | 24 | S26 | 0.26 | 0.05 | 21 | S34 | 0.24 | 0.06 | 14 |
| 41 | S46 | 0.27 | 0.03 | 24 | S4 | 0.25 | 0.07 | 21 | S30 | 0.23 | 0.06 | 14 |
| 42 | S44 | 0.26 | 0.05 | 24 | S53 | 0.24 | 0.04 | 21 | S33 | 0.21 | 0.05 | 14 |
| 43 | S33 | 0.25 | 0.03 | 24 | S58 | 0.23 | 0.05 | 21 | S43 | 0.20 | 0.06 | 14 |
| 44 | S34 | 0.24 | 0.03 | 24 | S33 | 0.23 | 0.04 | 21 | S3 | 0.19 | 0.10 | 14 |
| 45 | S4 | 0.22 | 0.05 | 24 | S43 | 0.20 | 0.04 | 21 | S29 | 0.09 | 0.05 | 14 |
| 46 | S52 | 0.22 | 0.03 | 24 | S6 | 0.16 | 0.07 | 21 | S46 | 0.09 | 0.02 | 14 |
| 47 | S43 | 0.22 | 0.04 | 24 | S36 | 0.12 | 0.02 | 21 | S4 | 0.09 | 0.03 | 14 |
| 48 | S38 | 0.18 | 0.08 | 24 | S40 | 0.08 | 0.01 | 21 | S36 | 0.08 | 0.03 | 14 |
| 49 | S40 | 0.17 | 0.04 | 24 | S8 | 0.08 | 0.02 | 21 | S6 | 0.07 | 0.02 | 14 |
| 50 | S29 | 0.14 | 0.03 | 24 | S29 | 0.07 | 0.02 | 21 | S7 | 0.06 | 0.06 | 14 |
| 51 | S8 | 0.09 | 0.02 | 24 | S38 | 0.07 | 0.02 | 21 | S38 | 0.05 | 0.02 | 14 |
| 52 | S16 | 0.08 | 0.01 | 24 | S35 | 0.06 | 0.02 | 21 | S40 | 0.05 | 0.02 | 14 |
| 53 | S36 | 0.08 | 0.01 | 24 | S46 | 0.06 | 0.02 | 21 | S47 | 0.05 | 0.03 | 14 |
| 54 | S14 | 0.06 | 0.01 | 24 | S16 | 0.05 | 0.02 | 21 | S5 | 0.05 | 0.03 | 14 |
| 55 | S35 | 0.04 | 0.01 | 24 | S47 | 0.02 | 0.01 | 21 | S16 | 0.04 | 0.02 | 14 |
| 56 | S45 | 0.03 | 0.01 | 24 | S39 | 0.01 | 0.01 | 21 | S8 | 0.04 | 0.01 | 14 |
| 57 | S47 | 0.02 | 0.00 | 24 | S14 | 0.01 | 0.00 | 21 | S35 | 0.02 | 0.01 | 14 |
| 58 | S5 | 0.02 | 0.00 | 24 | S7 | 0.01 | 0.00 | 21 | S14 | 0.02 | 0.01 | 14 |
| 59 | S15 | 0.01 | 0.00 | 24 | S5 | 0.01 | 0.00 | 21 | S45 | 0.02 | 0.00 | 14 |
| 60 | S13 | 0.01 | 0.00 | 24 | S15 | 0.01 | 0.00 | 21 | S15 | 0.01 | 0.01 | 14 |
| 61 | S37 | 0.01 | 0.00 | 24 | S45 | 0.01 | 0.00 | 21 | S39 | 0.01 | 0.00 | 14 |
| 62 | S7 | 0.01 | 0.00 | 24 | S13 | 0.00 | 0.00 | 21 | S37 | 0.01 | 0.00 | 14 |
| 63 | S39 | 0.00 | 0.00 | 24 | S37 | 0.00 | 0.00 | 21 | S13 | 0.00 | 0.00 | 14 |

**Supplementary Table 10. Comparing Boolean CAF subpopulations between tumour types.** Percentages from three independent repeats combined. Multiple t-test without assuming equal variance and with FDR correction for multiple comparisons (Q=1%). Significant results reported in main Table 2.

| Multiple t-tests 4T1 vs 4T07 D7 | | | | | | | | |
|---|---|---|---|---|---|---|---|---|
| CAF subset | Discovery? | P value | Mean1 4T1 | Mean2 4T07 | Difference | SE of difference | t ratio | df |
| S1 | | 0.670128 | 3.55 | 3.15 | 0.41 | 0.94 | 0.429 | 43 |
| S2 | | 0.115340 | 1.40 | 1.78 | -0.38 | 0.24 | 1.607 | 43 |
| S3 | | 0.113867 | 0.59 | 0.27 | 0.31 | 0.19 | 1.614 | 43 |
| S4 | | 0.197771 | 0.40 | 0.22 | 0.17 | 0.13 | 1.308 | 43 |
| S5 | | 0.006119 | 0.07 | 0.02 | 0.05 | 0.02 | 2.884 | 43 |
| S6 | | 0.194937 | 0.18 | 0.40 | -0.22 | 0.17 | 1.317 | 43 |
| S7 | | 0.018639 | 0.03 | 0.01 | 0.02 | 0.01 | 2.445 | 43 |
| S8 | | 0.775064 | 0.10 | 0.09 | 0.01 | 0.03 | 0.288 | 43 |
| S9 | | 0.750949 | 4.20 | 4.48 | -0.28 | 0.87 | 0.319 | 43 |
| S10 | | 0.332852 | 3.78 | 2.97 | 0.80 | 0.82 | 0.979 | 43 |
| S11 | | 0.041256 | 1.92 | 0.94 | 0.98 | 0.47 | 2.104 | 43 |
| S12 | * | 0.001220 | 1.95 | 0.69 | 1.26 | 0.36 | 3.463 | 43 |
| S13 | | 0.708238 | 0.01 | 0.01 | 0.00 | 0.01 | 0.377 | 43 |
| S14 | | 0.384671 | 0.08 | 0.06 | 0.02 | 0.02 | 0.878 | 43 |
| S15 | | 0.149474 | 0.03 | 0.01 | 0.01 | 0.01 | 1.468 | 43 |
| S16 | | 0.574243 | 0.07 | 0.08 | -0.01 | 0.02 | 0.566 | 43 |
| S17 | | 0.620748 | 5.82 | 5.33 | 0.48 | 0.97 | 0.498 | 43 |
| S18 | | 0.237856 | 1.74 | 2.07 | -0.33 | 0.28 | 1.197 | 43 |
| S19 | | 0.369887 | 2.02 | 1.52 | 0.50 | 0.55 | 0.906 | 43 |
| S20 | | 0.871153 | 0.70 | 0.72 | -0.02 | 0.15 | 0.163 | 43 |
| S21 | | 0.159622 | 1.04 | 0.76 | 0.28 | 0.20 | 1.431 | 43 |
| S22 | | 0.072977 | 6.87 | 9.39 | -2.53 | 1.37 | 1.838 | 43 |
| S23 | | 0.123313 | 0.44 | 0.62 | -0.17 | 0.11 | 1.572 | 43 |
| S24 | * | 0.001268 | 3.24 | 7.23 | -3.99 | 1.16 | 3.450 | 43 |
| S25 | | 0.134095 | 2.59 | 3.55 | -0.96 | 0.63 | 1.527 | 43 |
| S26 | | 0.463034 | 0.71 | 0.87 | -0.16 | 0.22 | 0.740 | 43 |
| S27 | | 0.142609 | 4.83 | 2.96 | 1.87 | 1.25 | 1.494 | 43 |
| S28 | | 0.631965 | 1.00 | 0.88 | 0.12 | 0.24 | 0.482 | 43 |
| S29 | | 0.085995 | 0.08 | 0.14 | -0.06 | 0.03 | 1.757 | 43 |
| S30 | | 0.006813 | 0.19 | 0.43 | -0.23 | 0.08 | 2.843 | 43 |
| S31 | | 0.665055 | 0.50 | 0.43 | 0.07 | 0.16 | 0.436 | 43 |
| S32 | * | 0.000918 | 5.84 | 14.44 | -8.60 | 2.41 | 3.561 | 43 |
| S33 | * | 0.000139 | 0.51 | 0.25 | 0.26 | 0.06 | 4.183 | 43 |
| S34 | | 0.082846 | 0.35 | 0.24 | 0.10 | 0.06 | 1.776 | 43 |
| S35 | | 0.015673 | 0.10 | 0.04 | 0.06 | 0.02 | 2.516 | 43 |
| S36 | | 0.203425 | 0.11 | 0.08 | 0.03 | 0.02 | 1.292 | 43 |
| S37 | | 0.032380 | 0.02 | 0.01 | 0.01 | 0.01 | 2.211 | 43 |
| S38 | | 0.782050 | 0.16 | 0.18 | -0.03 | 0.10 | 0.278 | 43 |
| S39 | | 0.039842 | 0.02 | 0.00 | 0.01 | 0.01 | 2.120 | 43 |
| S40 | | 0.788940 | 0.16 | 0.17 | -0.02 | 0.06 | 0.269 | 43 |
| S41 | * | 0.000789 | 3.56 | 1.04 | 2.52 | 0.70 | 3.612 | 43 |
| S42 | * | 0.000322 | 2.44 | 0.74 | 1.69 | 0.43 | 3.911 | 43 |
| S43 | * | 0.000008 | 0.67 | 0.22 | 0.45 | 0.09 | 5.074 | 43 |
| S44 | * | 0.000001 | 0.81 | 0.26 | 0.55 | 0.10 | 5.590 | 43 |
| S45 | | 0.044907 | 0.07 | 0.03 | 0.04 | 0.02 | 2.066 | 43 |
| S46 | | 0.010461 | 0.51 | 0.27 | 0.24 | 0.09 | 2.677 | 43 |
| S47 | | 0.002508 | 0.06 | 0.02 | 0.03 | 0.01 | 3.211 | 43 |
| S48 | | 0.043457 | 1.16 | 0.66 | 0.51 | 0.24 | 2.081 | 43 |
| S49 | * | 0.001231 | 1.28 | 0.77 | 0.51 | 0.15 | 3.460 | 43 |
| S50 | | 0.050778 | 0.61 | 0.41 | 0.20 | 0.10 | 2.010 | 43 |
| S51 | | 0.388689 | 0.39 | 0.31 | 0.08 | 0.09 | 0.871 | 43 |
| S52 | | 0.097134 | 0.37 | 0.22 | 0.15 | 0.09 | 1.696 | 43 |
| S53 | | 0.785562 | 0.57 | 0.52 | 0.06 | 0.21 | 0.274 | 43 |
| S54 | | 0.914057 | 4.45 | 4.52 | -0.08 | 0.71 | 0.109 | 43 |
| S55 | | 0.893838 | 1.27 | 1.18 | 0.09 | 0.64 | 0.134 | 43 |
| S56 | | 0.322404 | 8.07 | 9.48 | -1.40 | 1.40 | 1.001 | 43 |
| S57 | | 0.006089 | 3.23 | 1.70 | 1.53 | 0.53 | 2.885 | 43 |
| S58 | | 0.010897 | 0.95 | 0.44 | 0.51 | 0.19 | 2.661 | 43 |
| S59 | | 0.002165 | 2.15 | 1.02 | 1.13 | 0.35 | 3.263 | 43 |
| S60 | | 0.017022 | 0.58 | 0.35 | 0.23 | 0.09 | 2.483 | 43 |
| S61 | | 0.207491 | 0.34 | 0.51 | -0.17 | 0.14 | 1.280 | 43 |
| S62 | | 0.031101 | 2.06 | 1.36 | 0.70 | 0.31 | 2.229 | 43 |
| S63 | | 0.868287 | 7.07 | 6.49 | 0.57 | 3.45 | 0.167 | 43 |

| Multiple t-tests 4T1 vs 4T07 D14 | | | | | | | | |
|---|---|---|---|---|---|---|---|---|
| CAF subset | Discovery? | P value | Mean1 4T1 | Mean2 4T07 | Difference | SE of difference | t ratio | df |
| S1 | | 0.862189 | 1.66 | 1.59 | 0.07 | 0.42 | 0.175 | 43 |
| S2 | | 0.005825 | 1.94 | 1.09 | 0.85 | 0.29 | 2.902 | 43 |
| S3 | | 0.018291 | 0.16 | 0.45 | -0.29 | 0.12 | 2.453 | 43 |
| S4 | | 0.521111 | 0.32 | 0.25 | 0.07 | 0.11 | 0.647 | 43 |
| S5 | | 0.641022 | 0.01 | 0.01 | 0.00 | 0.00 | 0.470 | 43 |
| S6 | | 0.225144 | 0.08 | 0.16 | -0.08 | 0.07 | 1.231 | 43 |
| S7 | | 0.066966 | 0.00 | 0.01 | -0.01 | 0.00 | 1.879 | 43 |
| S8 | | 0.672961 | 0.09 | 0.08 | 0.02 | 0.04 | 0.425 | 43 |
| S9 | | 0.053233 | 1.36 | 0.88 | 0.48 | 0.24 | 1.988 | 43 |
| S10 | * | 0.000009 | 3.84 | 0.56 | 3.28 | 0.65 | 5.047 | 43 |
| S11 | | 0.899971 | 0.58 | 0.61 | -0.03 | 0.22 | 0.126 | 43 |
| S12 | * | 0.000014 | 2.07 | 0.36 | 1.70 | 0.35 | 4.905 | 43 |
| S13 | | 0.439272 | 0.00 | 0.00 | 0.00 | 0.00 | 0.781 | 43 |
| S14 | * | 0.000159 | 0.05 | 0.01 | 0.04 | 0.01 | 4.140 | 43 |
| S15 | | 0.218424 | 0.00 | 0.01 | 0.00 | 0.00 | 1.255 | 43 |
| S16 | | 0.192132 | 0.09 | 0.05 | 0.04 | 0.03 | 1.325 | 43 |
| S17 | | 0.894037 | 3.39 | 3.29 | 0.10 | 0.76 | 0.134 | 43 |
| S18 | | 0.042731 | 2.49 | 1.88 | 0.62 | 0.30 | 2.088 | 43 |
| S19 | | 0.108571 | 1.14 | 1.94 | -0.80 | 0.49 | 1.639 | 43 |
| S20 | | 0.285773 | 1.05 | 1.33 | -0.28 | 0.26 | 1.081 | 43 |
| S21 | | 0.022461 | 0.26 | 0.69 | -0.43 | 0.18 | 2.368 | 43 |
| S22 | | 0.798111 | 10.38 | 9.88 | 0.50 | 1.94 | 0.257 | 43 |
| S23 | * | 0.000080 | 0.17 | 0.53 | -0.36 | 0.08 | 4.359 | 43 |
| S24 | | 0.025527 | 8.11 | 12.32 | -4.21 | 1.82 | 2.314 | 43 |
| S25 | | 0.069355 | 1.92 | 1.16 | 0.76 | 0.41 | 1.863 | 43 |
| S26 | * | 0.000081 | 0.85 | 0.26 | 0.59 | 0.13 | 4.356 | 43 |
| S27 | | 0.870810 | 3.78 | 3.56 | 0.23 | 1.38 | 0.164 | 43 |
| S28 | | 0.218536 | 1.46 | 0.97 | 0.49 | 0.39 | 1.249 | 43 |
| S29 | | 0.002540 | 0.02 | 0.07 | -0.05 | 0.02 | 3.206 | 43 |
| S30 | | 0.148259 | 0.38 | 0.27 | 0.11 | 0.07 | 1.472 | 43 |
| S31 | | 0.030607 | 0.16 | 0.48 | -0.32 | 0.14 | 2.236 | 43 |
| S32 | | 0.832101 | 16.45 | 17.32 | -0.87 | 4.08 | 0.213 | 43 |
| S33 | | 0.119713 | 0.16 | 0.23 | -0.08 | 0.05 | 1.588 | 43 |
| S34 | | 0.018420 | 0.48 | 0.31 | 0.17 | 0.07 | 2.450 | 43 |
| S35 | | 0.194486 | 0.03 | 0.06 | -0.03 | 0.02 | 1.318 | 43 |
| S36 | | 0.235245 | 0.19 | 0.12 | 0.07 | 0.06 | 1.204 | 43 |
| S37 | | 0.202029 | 0.00 | 0.00 | 0.00 | 0.00 | 1.296 | 43 |
| S38 | | 0.477959 | 0.08 | 0.07 | 0.01 | 0.02 | 0.716 | 43 |
| S39 | | 0.195444 | 0.00 | 0.01 | -0.01 | 0.01 | 1.315 | 43 |
| S40 | | 0.013852 | 0.23 | 0.08 | 0.15 | 0.06 | 2.566 | 43 |
| S41 | | 0.321236 | 0.74 | 0.55 | 0.19 | 0.19 | 1.003 | 43 |
| S42 | * | 0.001271 | 2.72 | 0.55 | 2.18 | 0.63 | 3.449 | 43 |
| S43 | | 0.743423 | 0.18 | 0.20 | -0.02 | 0.05 | 0.329 | 43 |
| S44 | | 0.006553 | 1.47 | 0.35 | 1.12 | 0.39 | 2.858 | 43 |
| S45 | | 0.225895 | 0.01 | 0.01 | 0.01 | 0.00 | 1.229 | 43 |
| S46 | * | 0.000007 | 0.45 | 0.06 | 0.39 | 0.08 | 5.109 | 43 |
| S47 | | 0.046365 | 0.01 | 0.02 | -0.02 | 0.01 | 2.051 | 43 |
| S48 | * | 0.000469 | 0.90 | 0.29 | 0.61 | 0.16 | 3.787 | 43 |
| S49 | | 0.618470 | 0.31 | 0.35 | -0.03 | 0.06 | 0.502 | 43 |
| S50 | | 0.229197 | 0.70 | 0.54 | 0.16 | 0.13 | 1.220 | 43 |
| S51 | | 0.027193 | 0.10 | 0.28 | -0.18 | 0.08 | 2.287 | 43 |
| S52 | | 0.165022 | 0.53 | 0.71 | -0.18 | 0.13 | 1.412 | 43 |
| S53 | * | 0.001104 | 0.09 | 0.24 | -0.15 | 0.04 | 3.498 | 43 |
| S54 | | 0.226255 | 6.41 | 4.70 | 1.71 | 1.40 | 1.228 | 43 |
| S55 | | 0.025723 | 0.44 | 1.52 | -1.08 | 0.47 | 2.311 | 43 |
| S56 | | 0.038064 | 13.98 | 17.73 | -3.74 | 1.75 | 2.140 | 43 |
| S57 | | 0.149292 | 0.65 | 0.44 | 0.20 | 0.14 | 1.468 | 43 |
| S58 | * | 0.001065 | 0.67 | 0.23 | 0.43 | 0.12 | 3.510 | 43 |
| S59 | | 0.926590 | 0.44 | 0.43 | 0.01 | 0.10 | 0.093 | 43 |
| S60 | | 0.615706 | 0.62 | 0.57 | 0.05 | 0.10 | 0.506 | 43 |
| S61 | | 0.002566 | 0.08 | 0.36 | -0.28 | 0.09 | 3.202 | 43 |
| S62 | | 0.014361 | 1.66 | 0.89 | 0.77 | 0.30 | 2.552 | 43 |
| S63 | | 0.118327 | 1.40 | 6.03 | -4.63 | 2.91 | 1.594 | 43 |

| Multiple t-tests 4T1 vs 4T07 D21 | | | | | | | | |
|---|---|---|---|---|---|---|---|---|
| CAF subset | Discovery? | P value | Mean1 4T1 | Mean2 4T07 | Difference | SE of difference | t ratio | df |
| S1 | | 0.066789 | 1.66 | 0.76 | 0.90 | 0.47 | 1.890 | 36 |
| S2 | * | 0.000003 | 1.97 | 0.39 | 1.58 | 0.29 | 5.503 | 36 |
| S3 | | 0.506516 | 0.29 | 0.19 | 0.09 | 0.14 | 0.671 | 36 |
| S4 | | 0.045289 | 0.46 | 0.09 | 0.38 | 0.18 | 2.074 | 36 |
| S5 | | 0.223138 | 0.02 | 0.05 | -0.03 | 0.02 | 1.240 | 36 |
| S6 | | 0.011930 | 0.15 | 0.07 | 0.09 | 0.03 | 2.648 | 36 |
| S7 | | 0.211691 | 0.01 | 0.06 | -0.05 | 0.04 | 1.272 | 36 |
| S8 | | 0.017864 | 0.21 | 0.04 | 0.17 | 0.07 | 2.482 | 36 |
| S9 | | 0.038304 | 1.50 | 0.71 | 0.79 | 0.37 | 2.151 | 36 |
| S10 | | 0.001291 | 2.97 | 0.53 | 2.44 | 0.70 | 3.491 | 36 |
| S11 | | 0.156885 | 0.59 | 0.33 | 0.26 | 0.18 | 1.446 | 36 |
| S12 | * | 0.000914 | 1.35 | 0.25 | 1.10 | 0.31 | 3.614 | 36 |
| S13 | | 0.839347 | 0.00 | 0.00 | 0.00 | 0.00 | 0.204 | 36 |
| S14 | | 0.209145 | 0.03 | 0.02 | 0.01 | 0.01 | 1.279 | 36 |
| S15 | | 0.203697 | 0.01 | 0.01 | -0.01 | 0.01 | 1.295 | 36 |
| S16 | | 0.038017 | 0.21 | 0.04 | 0.17 | 0.08 | 2.154 | 36 |
| S17 | | 0.243631 | 2.72 | 2.10 | 0.62 | 0.52 | 1.185 | 36 |
| S18 | | 0.003042 | 2.09 | 0.91 | 1.18 | 0.37 | 3.178 | 36 |
| S19 | | 0.955422 | 1.10 | 1.12 | -0.02 | 0.35 | 0.056 | 36 |
| S20 | | 0.034774 | 1.17 | 0.66 | 0.51 | 0.23 | 2.194 | 36 |
| S21 | * | 0.000303 | 0.38 | 1.17 | -0.79 | 0.20 | 3.998 | 36 |
| S22 | | 0.164183 | 13.21 | 8.91 | 4.29 | 3.02 | 1.420 | 36 |
| S23 | | 0.020824 | 0.57 | 1.02 | -0.45 | 0.19 | 2.417 | 36 |
| S24 | | 0.230951 | 11.35 | 8.95 | 2.40 | 1.97 | 1.219 | 36 |
| S25 | | 0.124650 | 2.02 | 1.13 | 0.89 | 0.56 | 1.572 | 36 |
| S26 | | 0.039129 | 0.64 | 0.29 | 0.35 | 0.16 | 2.141 | 36 |
| S27 | | 0.502708 | 1.92 | 2.45 | -0.53 | 0.78 | 0.677 | 36 |
| S28 | | 0.686840 | 0.64 | 0.57 | 0.07 | 0.17 | 0.406 | 36 |
| S29 | | 0.167042 | 0.04 | 0.09 | -0.06 | 0.04 | 1.410 | 36 |
| S30 | | 0.004817 | 0.43 | 0.23 | 0.21 | 0.07 | 3.005 | 36 |
| S31 | | 0.088126 | 0.37 | 1.06 | -0.68 | 0.39 | 1.753 | 36 |
| S32 | | 0.669595 | 15.09 | 13.72 | 1.38 | 3.20 | 0.430 | 36 |
| S33 | | 0.853266 | 0.22 | 0.21 | 0.02 | 0.08 | 0.186 | 36 |
| S34 | | 0.028409 | 0.45 | 0.24 | 0.21 | 0.09 | 2.283 | 36 |
| S35 | | 0.214440 | 0.05 | 0.02 | 0.03 | 0.02 | 1.264 | 36 |
| S36 | | 0.095830 | 0.28 | 0.08 | 0.20 | 0.12 | 1.710 | 36 |
| S37 | | 0.375778 | 0.01 | 0.01 | 0.01 | 0.01 | 0.897 | 36 |
| S38 | | 0.053825 | 0.11 | 0.05 | 0.06 | 0.03 | 1.994 | 36 |
| S39 | | 0.531654 | 0.01 | 0.01 | 0.00 | 0.01 | 0.632 | 36 |
| S40 | | 0.026662 | 0.31 | 0.05 | 0.26 | 0.11 | 2.311 | 36 |
| S41 | | 0.104877 | 0.53 | 0.84 | -0.31 | 0.19 | 1.664 | 36 |
| S42 | | 0.126384 | 1.79 | 0.99 | 0.81 | 0.52 | 1.565 | 36 |
| S43 | | 0.686055 | 0.25 | 0.20 | 0.04 | 0.10 | 0.407 | 36 |
| S44 | | 0.162772 | 1.13 | 0.62 | 0.51 | 0.36 | 1.425 | 36 |
| S45 | | 0.060755 | 0.01 | 0.02 | -0.01 | 0.00 | 1.936 | 36 |
| S46 | | 0.001589 | 0.24 | 0.09 | 0.15 | 0.05 | 3.416 | 36 |
| S47 | | 0.500512 | 0.03 | 0.05 | -0.02 | 0.02 | 0.681 | 36 |
| S48 | | 0.252844 | 1.06 | 0.59 | 0.48 | 0.41 | 1.162 | 36 |
| S49 | | 0.004597 | 0.31 | 0.58 | -0.27 | 0.09 | 3.023 | 36 |
| S50 | | 0.227580 | 0.53 | 0.68 | -0.15 | 0.12 | 1.228 | 36 |
| S51 | | 0.066808 | 0.13 | 0.31 | -0.17 | 0.09 | 1.890 | 36 |
| S52 | | 0.067913 | 0.55 | 0.32 | 0.22 | 0.12 | 1.882 | 36 |
| S53 | * | 0.000005 | 0.10 | 0.56 | -0.46 | 0.09 | 5.354 | 36 |
| S54 | | 0.497700 | 6.19 | 7.66 | -1.47 | 2.14 | 0.685 | 36 |
| S55 | | 0.019206 | 0.82 | 2.31 | -1.49 | 0.61 | 2.452 | 36 |
| S56 | | 0.095990 | 14.55 | 16.88 | -2.32 | 1.36 | 1.709 | 36 |
| S57 | | 0.402479 | 0.52 | 0.63 | -0.12 | 0.14 | 0.847 | 36 |
| S58 | | 0.695238 | 0.50 | 0.58 | -0.08 | 0.19 | 0.395 | 36 |
| S59 | | 0.170736 | 0.46 | 0.71 | -0.26 | 0.18 | 1.398 | 36 |
| S60 | | 0.657369 | 0.69 | 0.62 | 0.08 | 0.17 | 0.447 | 36 |
| S61 | * | 0.000009 | 0.07 | 0.80 | -0.73 | 0.14 | 5.158 | 36 |
| S62 | | 0.478906 | 1.47 | 1.72 | -0.25 | 0.35 | 0.716 | 36 |
| S63 | * | 0.000849 | 1.45 | 13.69 | -12.24 | 3.36 | 3.640 | 36 |
